# Spatially resolved human kidney multi-omics single cell atlas highlights the key role of the fibrotic microenvironment in kidney disease progression

**DOI:** 10.1101/2022.10.24.513598

**Authors:** Amin Abedini, Jonathan Levinsohn, Konstantin A Klötzer, Bernhard Dumoulin, Ziyuan Ma, Julia Frederick, Poonam Dhillon, Michael S Balzer, Rojesh Shrestha, Hongbo Liu, Steven Vitale, Kishor Devalaraja-Narashimha, Paola Grandi, Tanmoy Bhattacharyya, Erding Hu, Steven S. Pullen, Carine M Boustany-Kari, Paolo Guarnieri, Anil Karihaloo, Daniel Traum, Hanying Yan, Kyle Coleman, Matthew Palmer, Lea Sarov-Blat, Lori Morton, Christopher A. Hunter, Klaus H Kaestner, Mingyao Li, Katalin Susztak

**Affiliations:** Renal, Electrolyte, and Hypertension Division, Department of Medicine, University of Pennsylvania, Perelman School of Medicine, Philadelphia, PA 19104, USA; Institute for Diabetes, Obesity, and Metabolism, University of Pennsylvania, Perelman School of Medicine, Philadelphia, PA 19104, USA; Penn/CHOP Kidney Innovation Center, University of Pennsylvania, Perelman School of Medicine, Philadelphia, PA 19104, USA; Department of Genetics, University of Pennsylvania, Perelman School of Medicine, Philadelphia, PA 19104, USA; Cardiovascular, Renal and Fibrosis Research, Regeneron Pharmaceuticals Inc., Tarrytown, NY 10591, USA; Research and Development, GSK, Cellzome GmbH, Genomic Sciences, GSK, Heidelberg, Germany; Research and Development, GSK, Crescent Drive, Philadelphia, Pennsylvania, USA; Department of Cardiometabolic Diseases Research, Boehringer Ingelheim Pharmaceuticals, Ridgefield, CT, USA; Novo Nordisk Research Center Seattle, Inc., Seattle, USA; Department of Epidemiology and Biostatistics, University of Pennsylvania Perelman School of Medicine, Philadelphia, PA, USA; Department of Pathology and Laboratory Medicine, University of Pennsylvania, Perelman School of Medicine, Philadelphia, Pennsylvania, USA; Department of Pathobiology, University of Pennsylvania School of Veterinary Medicine, Philadelphia, PA, USA

## Abstract

Kidneys possess one of the most intricate three-dimensional cellular structures in the body, yet the spatial and molecular principles of kidney health and disease remain inadequately understood. Here, we have generated high-quality datasets for 81 samples, including single cell (sc), single nuclear (sn), spot level (Visium) and single cell resolution (CosMx) spatial (sp)-RNA expression, and sn open chromatin, capturing cells from healthy, diabetic, and hypertensive diseased human kidneys. By combining the snRNA, snATAC and scRNA sequencing we identify cell types and map these cell types to their locations within the tissue. Unbiased deconvolution of the spatial data identifies 4 distinct spatial microenvironments: glomerular, immune, tubule and fibrotic. We describe the complex, heterogenous cellular and spatial organization of human microenvironments in health and disease. Further, we find that the fibrotic microenvironment spatial gene signature is not only able to molecularly classify human kidneys, but it also offers an improved prognosis prediction compared to traditional histopathological analysis. We provide a comprehensive spatially resolved molecular roadmap of the human kidney and the fibrotic process, demonstrating the clinical utility of spatial transcriptomics.

## Introduction

Human kidneys filter over 140 liters of plasma daily, reabsorb vital nutrients, excrete water and electrolytes, and eliminate toxins to maintain the internal milieu^1,2^. Kidney disease is defined by a decline in glomerular filtration. Chronic kidney disease (CKD) is the 9^th^ leading cause of death^3,4^ in the United States, affecting 14% of the population. Diabetes and hypertension are responsible for more than 75% of all CKD^5^.

More than 30 specialized cell types including epithelial, endothelial, interstitial, and immune cells, have been identified in the kidney^6,7^. The development of single cell and single nuclear RNA sequencing (scRNAseq, snRNAseq, respectively) as well as single nuclei Assay for Transposase-Accessible Chromatin sequencing (snATACseq) has provided an unprecedented insight into the molecular and cellular composition of healthy mouse and human kidneys, including changes during development and disease^8–12^. These methods use dissociated cells or nuclei isolated from kidney tissue samples. Despite the significant cellular diversity of the kidney, cell types could be accurately identified in scRNAseq and snRNAseq data, as their specialized cellular function matches with the cellular gene expression signatures^13^.

The lack of spatial information impedes the accurate mapping of known cell types only described by their anatomical location. This limitation hampers the interrogation of local gene expression changes and cell-cell communication, both of which play a vital role in maintaining cellular health and can be dysregulated in disease. The field of spatial omics is rapidly evolving, but currently available spatial datasets either lack single cell resolution information, unable to provide genome-scale gene expression data, or only evaluate a small spatial area. Clearly, there is a pressing need for large-scale spatial omics datasets to better understand kidney health and disease.

Chronic kidney disease (CKD) is associated with a complex change in the kidney’s cellular structure^14^. Some histologic changes are specific for disease subtypes; for instance, diffuse thickening of glomerular basement membrane is observed in diabetic kidney disease (DKD)^15^. However, fibrosis is a common manifestation across all forms of progressive CKD. The histologic hallmark of fibrosis is the accumulation of extracellular matrix^16,17^, which has been the primary focus of most prior studies. Matrix accumulation can cause organ stiffness, which is implicated in organ failure in cases of pulmonary and heart fibrosis^18–20^. The role of tissue elasticity in the regulation of kidney function is not clear^21^, and thus, the mechanism by which matrix accumulation (or fibrosis) influences kidney function remains a subject of controversy^22,23^.

One of the most significant clinical problems is identifying patients who will develop end-stage renal failure, requiring either a transplant or dialysis. Clinical parameters, including eGFR and albuminuria, can predict kidney function decline with modest accuracy for patients with advanced CKD stages ^24–26^. However, for patients in the earlier stages of disease, accurate prognosis remains a challenge^25^. Although some studies suggest that the severity of fibrosis can serve as a predictor, this finding has not been consistently reproduced^27–29^.

Here, we generate both spot (Visium) and single cell level (CosMx) spatial transcriptomic data for both healthy and diseased human kidneys in conjunction with sn/scRNAseq and snATACseq. By combining spatial gene expression with high quality single cell expression and open chromatin information, we resolve the identity of cells previously only known by their spatial localization. Moreover, we perform a detailed spatial and cellular characterization of tissue fibrosis. We demonstrate the cellular heterogeneity of the fibrotic stroma, which includes not only fibroblasts and myofibroblasts but also endothelial cells and immune cells that follow the organization of a lymphoid organ and are anatomically close to injured proximal tubule cells. We define various tissue microenvironments, including the fibrotic microenvironment (FME). Ultimately, our data show that the FME gene signature can classify kidney samples and predict future kidney function decline. These findings demonstrate the potential clinical utility of spatial transcriptomics.

## Results

### Spatially resolved multi-omics single cell survey of healthy and diseased human kidneys defines expression, gene regulation and spatial location of >100 cell types and states

We generated a comprehensive human kidney single cell atlas with spatial resolution by analyzing 81 human kidney tissue samples from 58 subjects (62% male, and age of 64.1 ± 14.0 years). Samples were divided into two groups: (i) healthy control (N = 36) determined by estimated glomerular filtration rate (eGFR) > 60 ml/min/1.73 *m*^2^ and fibrosis score of < 10 % (ii) chronic kidney disease (CKD) (N = 45) determined by (eGFR) < 60 ml/min/1.73 *m*^2^ or kidney fibrosis score of > 10%, including 20 with the clinical histopathological diagnosis of diabetic kidney disease (DKD) and 25 with the clinical diagnosis of hypertensive-attributed chronic kidney disease. **Supplementary Table 1** shows the demographic, clinical, and histological characteristics of the included samples.

We performed droplet-based single cell analysis using 10X Chromium Next GEM for sc/snRNAseq (N = 47), snATACseq (N = 18). After standard processing and meticulous quality control (**Supplementary** Fig. 1 and **Supplementary Table 2**), during which we removed low-quality cells, we included 338,565 cells/nuclei in our final atlas. We used SCVI tools to generate a single unified comprehensive integrated human kidney atlas including all different modalities and disease states^30^. Overall, we identified six cell super families, including endothelial cells, stromal cells, tubule epithelial cells, immune cell types, glomerular cells, and neural/Schwann cells. Within these large families we identified 44 main and 114 distinct cell sub-types or states in healthy and diseased human kidneys (**Fig. 1a**). Key cluster-specific gene markers are shown in **Supplementary** Fig. 2 and **Supplementary Tables 3 to 6**. Our sc and sn human kidney atlas captured kidney cell types in healthy and diseased states in all anatomical regions of the kidney. The main identified cell types were: podocytes, different types of proximal tubule segments 1-3 (PT_S1, S2, S3, and injured), descending thin loop of Henle, ascending thin loop of Henle, cortical and medullary thick ascending loop of Henle (C_TAL and M_TAL), distal convoluted tubule (DCT), connecting tubule (CNT), principal cells of collecting duct (PC), intercalated cells type alpha and beta (IC_A and IC_B), stromal, endothelial, and different types of immune cells.

**Fig 1.**
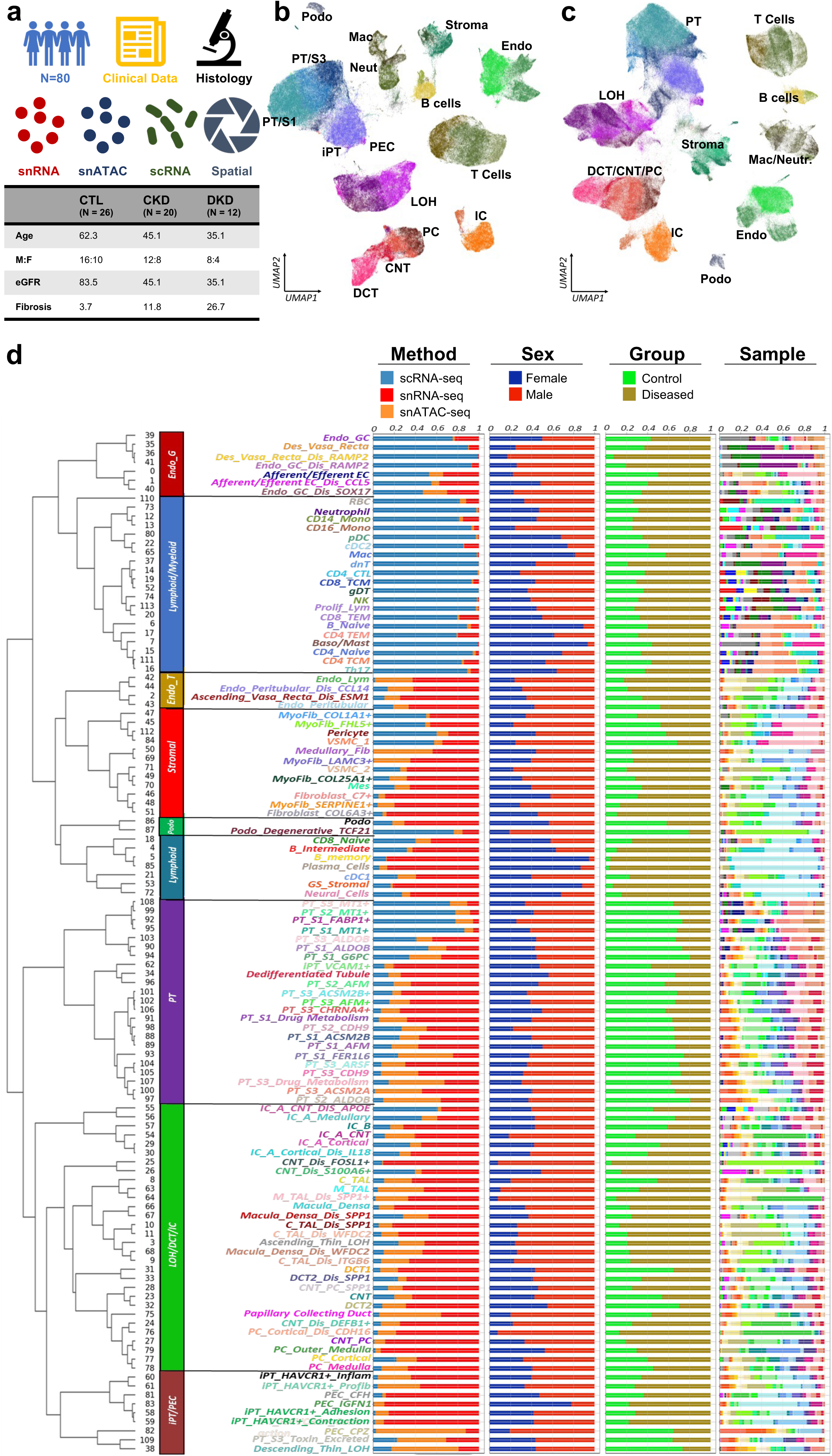
Comprehensive integrated multimodal human kidney single cell atlas. **(a).** Overview of the multimodal analysis. Basic clinical characteristics of the samples. **(b)** UMAP of 338,565 integrated human kidney sn/scRNAseq and snATACseq data generated in this study. Annotated cell types are indicated on the plot. **(c)** UMAP of 588,425 integrated human kidney sn/scRNAseq and snATACseq data of the present study and kidney precision medicine project (KPMP). **(d)** Hierarchical sub-clustering identified 114 distinct cell types or cell states. The bar charts depict the relative abundance of each group (method, sex, group, and samples) contributing to the cluster. Endo_G; endothelial cells of glomerular capillary tuft, Endo_peritubular; endothelial cells of peritubular vessels, Endo_lymphatic; endothelial cells of lymphatic vessels, Mes; mesangial cells, GS_Stromal; glomerulosclerosis-specific stromal cells, VSMC/Pericyte; vascular smooth muscle cells/pericyte, PEC; parietal epithelial cells, Podo; podocyte, PT_S1; proximal tubule segment 1, PT_S2; proximal tubule segment 2, PT_S3; proximal tubule segment 3, iPT; injured proximal tubule cells, DTL; descending thin loop of Henle, C_TAL; cortical thick ascending loop of Henle, M_TAL; medullary thick ascending loop of Henle, DCT; distal convoluted tubule, CNT; connecting tubule cells, PC; principal cells of collecting duct, IC_A; Type alpha intercalated cells, IC_B; Type beta intercalated cells, NK; natural killer cells, CD4T; T lymphocytes CD4+, CD8T; T lymphocytes CD8+, dnT; double negative T cells, Prolif_Lym; proliferative lymphocyte, Th17; T helper 17 lymphocyte, gDT; gamma delta T cells, TCM; T Cell memory, TEM; T effector memory, Naive B lymphocyte, B_memory; memory B lymphocyte, RBC; red blood cells, Baso/Mast; basophil or mast cells, pDC; plasmacytoid dendritic cells, cDC; classical dendritic cells, Mac; macrophage, CD14_Mono; monocyte CD14+, CD16_Mono; monocyte CD16+, CTL; control, HKD; hypertensive kidney disease, DKD; diabetic kidney disease.

The combination of single cell and single nuclear methods, the large number of analyzed cells, the high-quality dataset, and the inclusion of samples with different degrees of kidney disease severity in our kidney atlas enabled the capture of rare and novel cell types. We captured 11 different types of endothelial cells, including afferent, efferent arterioles (*FBLN5*+) and vasa recta (*MCTP1*+) cells (**Fig. 1b, c, and** **Supplementary** Fig. 2,3). We could also identify rare erythropoietin producing cells within the stromal cell population (**Supplementary** Fig. 3). We captured proximal tubule (PT) cells expressing high levels of *SLC47A2,* a gene specific for toxin excretion (**Supplementary** Fig. 2**, 3 and Supplementary Table 6)** and tubule epithelial subtypes mostly seen in diseased kidneys that were positive for *IL18*, *WFDC2*, *SPP1*, and *ITGB6*. Our atlas provides a comprehensive reference for human kidney immune cells. We identified lymphoid (CD4T, CD8T, natural killer cells, double-negative T cells, Th17, B_Naive, B_intermediate B_memory, plasma_cells) and myeloid cells (neutrophil, basophil/mast cells, CD14_monocyte, CD16_monocyte, macrophage, classical and plasmacytoid dendritic cells). We provide comprehensive hierarchical clustering, including information on sex, disease status, sample identity, and analytical methods (**Fig. 1d).** Furthermore, we created an easy-to-use web interface to examine gene expression enabling users to identify cell type-specific and disease state-specific changes in the human kidney (www.susztaklab.com/hk_genemap/)

To further improve and validate our cell type and state identification, we integrated our human kidney atlas with the human kidney precision medicine project (KPMP) dataset^31^ (**Fig. 1c, Supplementary** Fig. 4,5). This combined fully integrated atlas contains ∼600K cells or nuclei using three different analytical modalities validating the consistency of the cell type annotations between datasets (**Supplementary** Fig. 4**, 5).**

In addition to the gene expression data, the snATACseq of 57,847 human kidney nuclei provided us opportunities to identify transcription factors (TF) and enriched TF motifs in each cell type. Cell gene-expression markers, a comprehensive list of cell-type differentially accessible regions and transcription factors can be found in **Supplementary** Fig. 6 and **Supplementary Table 7** and include *WT1* for podocyte and parietal epithelial cells (PEC), *HNF4A* for PT cell types, *FOSL2* for injured_PT (iPT), and *TFAP2A* for C_TAL (**Supplementary** Fig. 6).

### High-resolution spatially resolved human kidney atlas

A key limitation to cell type identification has been the lack of high-resolution, spatially resolved cell transcriptomics information. To overcome this limitation, we used the new Visium FFPE platform and generated 14 spRNAseq data sets, including 3 control (healthy) and 11 diseased samples (7 DKD, 4 HKD) (**Supplementary Table 2, Supplementary** Fig. 7-9).

We then leveraged our dissociated sn/sc data to identify cell types within our spatial dataset. Firstly, for the Visium data we used the Cell2location^32^ package, which is a machine learning method that estimates the contribution of each cell type to the observed gene expression profiles. We then employed CellTrek^33^, which uses the dissociated data to impute the spatial location of cell types to near single cell level (149,717 datapoints after imputation). The high-resolution data enabled the projection of all identified cell types from the dissociated datasets to their spatial location (**Fig. 2a, b**). We could also identify markers for cell types previously only known by their anatomical location; for instance, PEC cells expressing *CFH*, and *WT1,* as well as mesangial cells expressing *ITGA8* and *POSTN*. The gene expression-based spatial map was consistent with the Human Protein Atlas^34^ data (**Supplementary** Fig. 10). Careful examination of the Cell2Location and CellTrek-based cell type prediction showed the expected anatomical localization (i.e. glomerular cells colocalizing at morphologic glomeruli, immune cells localizing with histologic immune infiltrate) (**Supplementary** Fig. 11 to 14).

**Fig 2.**
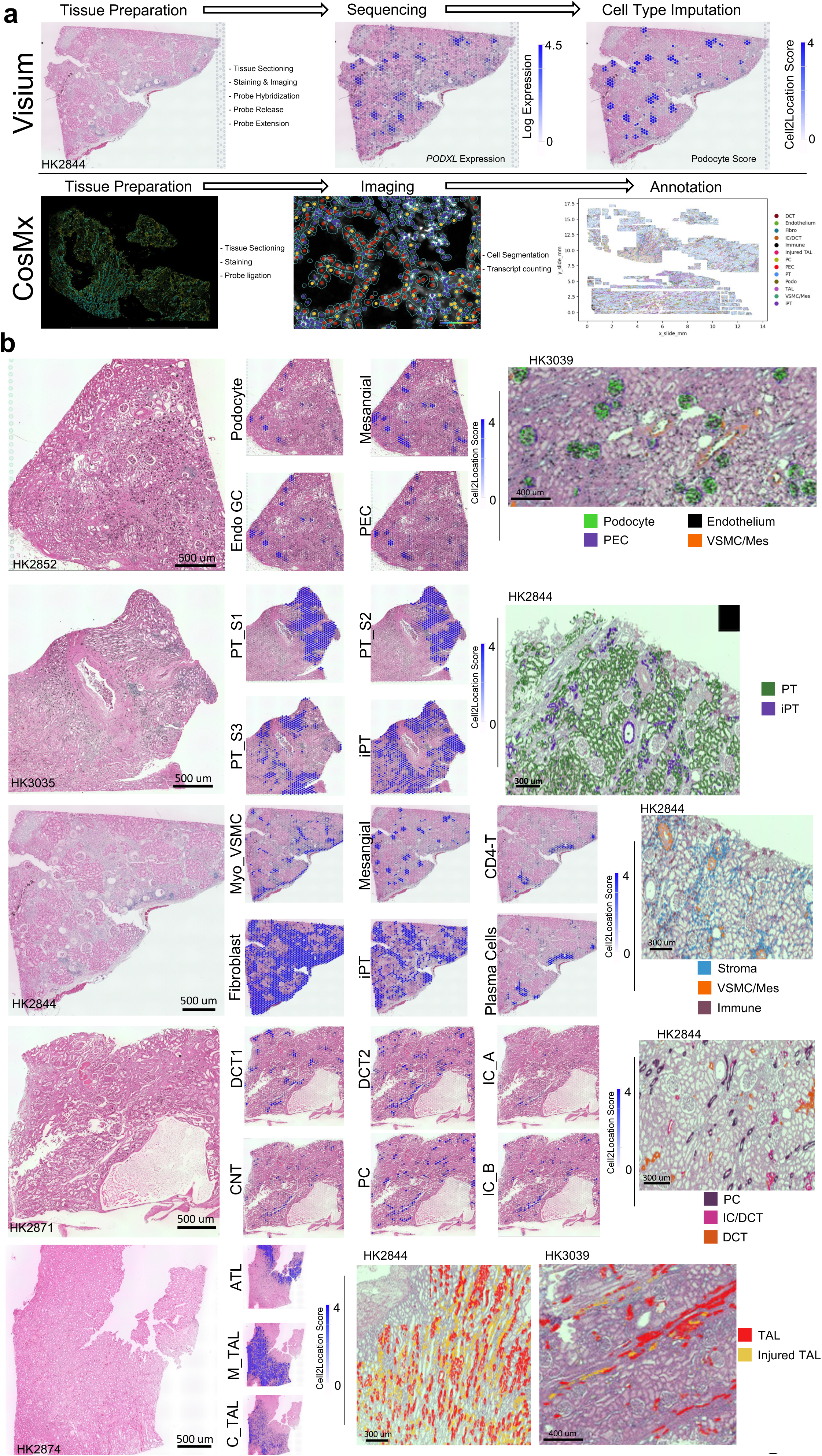
Spatially resolved human kidneys. **(a)** Spatial transcriptomics data was generated from human formalin fixed paraffin embedded kidney samples using two platforms. Visium (top) employs a spot-based approach with each spot of 55 um, with the ability to detect >18,000 genes able for detection and requires deconvolution to identify presence of individual cell types. CosMx (bottom) imaging generates single cell level data and identifies 1000 genes, and which permits annotation of cell types based on the expression patterns of these genes. **(b)** Spatial location and marker gene expression of identified cell types using both Visium and CosMx. H&E sections shown on the left of the figure show individual tissue histology of our Visium sections. Using our calculated Cell2Location scores for each of these tissues, we imputed the presence of cell types on each of these sections with cell type scores shown in blue overlaying the H&E. We identify glomerular cell types, proximal tubular cell types, immune and fibrotic cell types, distal tubular cell types and loop of Henle cell types. To the right of these images, we validated the location of these cell types using the CosMx assay, with allows for individual cell type annotations, which are overlayed on the H&E image of the assayed tissue. Endo_GC; endothelial cells of glomerular capillary tuft, Mes; mesangial cells, GS_Stromal; glomerulosclerosis-specific stromal cells, Myo_VSMC; myofibroblast/vascular smooth muscle cells PEC; parietal epithelial cells, Podocyte; podocyte, PT_S1; proximal tubule segment 1, PT_S2; proximal tubule segment 2, PT_S3; proximal tubule segment 3, iPT; injured proximal tubule cells, ATL: ascending thin limb of loop of Henle, C_TAL; cortical thick ascending loop of Henle, M_TAL; medullary thick ascending loop of Henle, DCT; distal convoluted tubule, CNT; connecting tubule cells, PC; principal cells of collecting duct, IC_A; Type alpha intercalated cells, IC_B; Type beta intercalated cells, CD4T; T lymphocytes CD4+, Plasma cell; plasma cell.

Finally, we then used the CosMx (NanoString) platform to assay human kidney samples (1 healthy, 1 DKD), at true single cell resolution analyzing the expression of 1,000 genes (**Supplementary** Fig. 7**, 15) and Supplementary Table 2**). The method generated high quality datasets and distinct UMAP cell clusters that corresponded to specific kidney cell types (**Supplementary** Fig. 16). The CosMx dataset seamlessly integrated with our single nuclei RNA dissociated cell data, further confirming the proper annotation of cell types (**Supplementary** Fig. 17). Furthermore, the gene expression-based cell type prediction fully matched with anatomical and histological annotation of the slide (**Fig. 2b**). The processed spatial transcriptomics (Visium and CosMx) dataset is fully available for the community and on our interactive website (www.susztaklab.com/hk_genemap/). https://susztaklab.com/samui Overall, we constructed an integrated high-quality spatially resolved human kidney multi-omics atlas, which allowed the spatial mapping of cellular gene expression and gene regulatory information in healthy and diseased states.

### Stromal cells heterogeneity and matrix production in human kidney disease

We next analyzed kidney fibrosis associated changes, as fibrosis is the common manifestation of all progressive chronic kidney disease. We created an extracellular matrix (ECM) score by combining the expression of collagens, glycoproteins, and proteoglycans^35,36^. **Fig. 3a** shows that myofibroblasts, followed by fibroblasts, and then vascular smooth muscle cells (VSMC)/pericytes had the highest ECM score. We noted that disease samples have a higher frequency of myofibroblasts compared to healthy samples **(Fig. 3b**) and myofibroblasts were located within high ECM score area (**Fig. 3c**).

**Fig 3.**
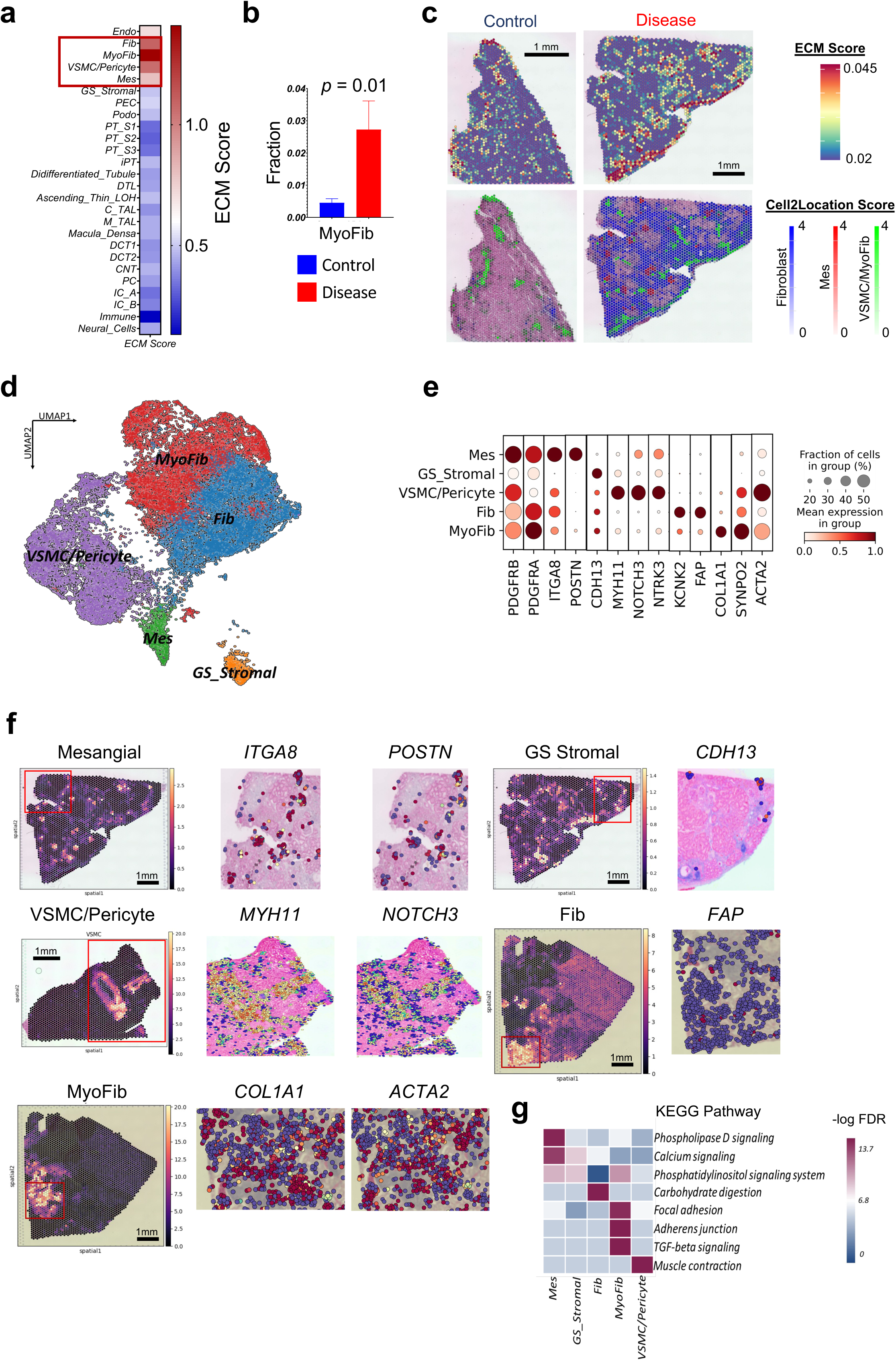
Human kidney stromal cell atlas. **(a)** Extracellular matrix (ECM) gene expression score in different kidney cell types in the integrated sn/scRNAseq and snATACseq. ECM score was calculated based on the expression of collagen, proteoglycan, and glycoprotein genes. **(b)** The fraction of myofibroblasts in control and CKD samples. The bars indicate SEM. Independent t-test was used to compare the fractions between two groups. **(c)** The ECM score in the spatial transcriptomics datasets of healthy and CKD samples (upper panel). Red indicates higher ECM gene expression (upper panel). The lower panel indicates the location of mesangial cells, fibroblasts, myofibroblasts and VSCMC/pericytes in healthy and CKD samples based on their Cell2Location score. **(d)** UMAP representation of sub-clustering of 32,706 stomal cells in the integrated dataset -- present study and Kidney Precision Medicine Project (KPMP). **(e)** The dot plots of marker genes used for stromal cell type annotation in the integrated dataset. The size of the dot indicates the percent positive cells, and the darkness of the color indicates average expression. **(f)** The spatial location and stromal cell subtypes and specific marker genes. The left panel shows the relative abundance of each cell type using Cell2location. The right panel shows relative gene expression using CellTrek (red higher). **(g)** The heatmap indicates the -Log_10_ (FDR) enrichment of the top KEGG pathways in each stromal cell type. Mes; mesangial cells, GS_Stromal; glomerulosclerois-specific stromal cells, Fib; fibroblast, VSMC; vascular smooth muscle cells, MyoFib; myofibroblast.

Our combined dataset enabled the identification of stromal cell subtypes that were consistent with prior publications and datasets (**Supplementary** Fig. 17,18) and with correct spatial location (**Supplementary** Fig. 19**-20)**. As previously known, stromal cells were positive for *PDGFRB* expression. We were able to identify markers for mesangial cells (*ITGA8*, *POSTN*), VSMC/pericytes (*MYH11*, *NOTCH3*, *NTRK3*), fibroblast (*KCNK2*, *FAP*), and myofibroblast (*COL1A1*, *SYNPO2*) (**Fig. 3d, e, Supplementary** Fig. 21). We validated the consistency of stromal subtypes with prior publications using the MetaNeighbor tool^37^ (**Supplementary** Fig. 22) and we verified their spatial location (**Fig. 3f**). The CosMx dataset separated 2 main stromal clusters, one of which comprised VSMC/pericyte/mesangial cells and the other fibroblasts/myofibroblasts. (**Supplementary** Fig. 21).

Further sub-clustering analysis of stromal cells distinguished 12 different cell subtypes (**Supplementary** Fig. 22). We captured medullary fibroblasts expressing *SYT1* and *NCAM1* and four different myofibroblasts marked by *COL1A1*+, *CLMP*+, *FGF7*+, or *ITGBL1*+ expression (**Supplementary** Fig. 22). We could discriminate VSMC from myofibroblasts based on the expression of *MYH11*, *NOTCH3, CNN1*, and *NTRK3* (**Fig 3f, Supplementary** Fig. 21). These annotations were also consistent with protein expression in the Human Protein Atlas (**Supplementary** Fig. 23) and snATACseq analysis (**Supplementary** Fig. 24). Gene ontology analysis highlighted important differences between the different stromal cells, such as myofibroblasts showed enrichment for focal adhesion, and TGFB-signaling pathway genes (**Fig. 3g)**. Weighted gene co-expression network analysis (WGCNA) ^38,39^ indicated enrichment for *GUCY1A2*, *LAMA2*, and *MYO1D* in the fibroblast module and *COL1A1*, *ZEB2*, *SVEP1*, and *PTEN* in the myofibroblast module in disease states (**Supplementary** Fig. 21**, Supplementary Table 8)**. In the snATACseq data, we could identify enrichment for *NPAS2*, *TEAD2*, and *TCF7* motifs in open chromatin areas in myofibroblasts and *USF2* transcription factor motifs in VSMC (**Supplementary** Fig. 24).

In summary, we have generated a high-resolution, spatially resolved stromal cell atlas and identified ECM producing cells in diseased human kidneys.

### The kidney fibrotic microenvironment is established by the complex interaction of injured epithelial cells, immune cells, and endothelial cells

Our newly generated spRNAseq dataset is uniquely suited to defining microenvironments (ME) in the human kidney. We ran nonnegative matrix factorization (NMF) on the Visium datasets using the STutility^40^ package. We found four major MEs in the human kidney, that visually corresponded to glomerular, tubule, fibrotic, and immune regions. Gene ontology enrichment analysis of genes detected in each microenvironment was consistent with their anatomical annotation (**Supplementary** Fig. 25). The computationally defined fibrotic microenvironment (FME) strongly correlated with kidney ECM gene expression (**Supplementary** Fig. 26) and our pathologist’s assessment of fibrosis (r=0.75, *p*=0.003) (**Fig. 4a, Supplementary** Fig. 26).

**Fig 4.**
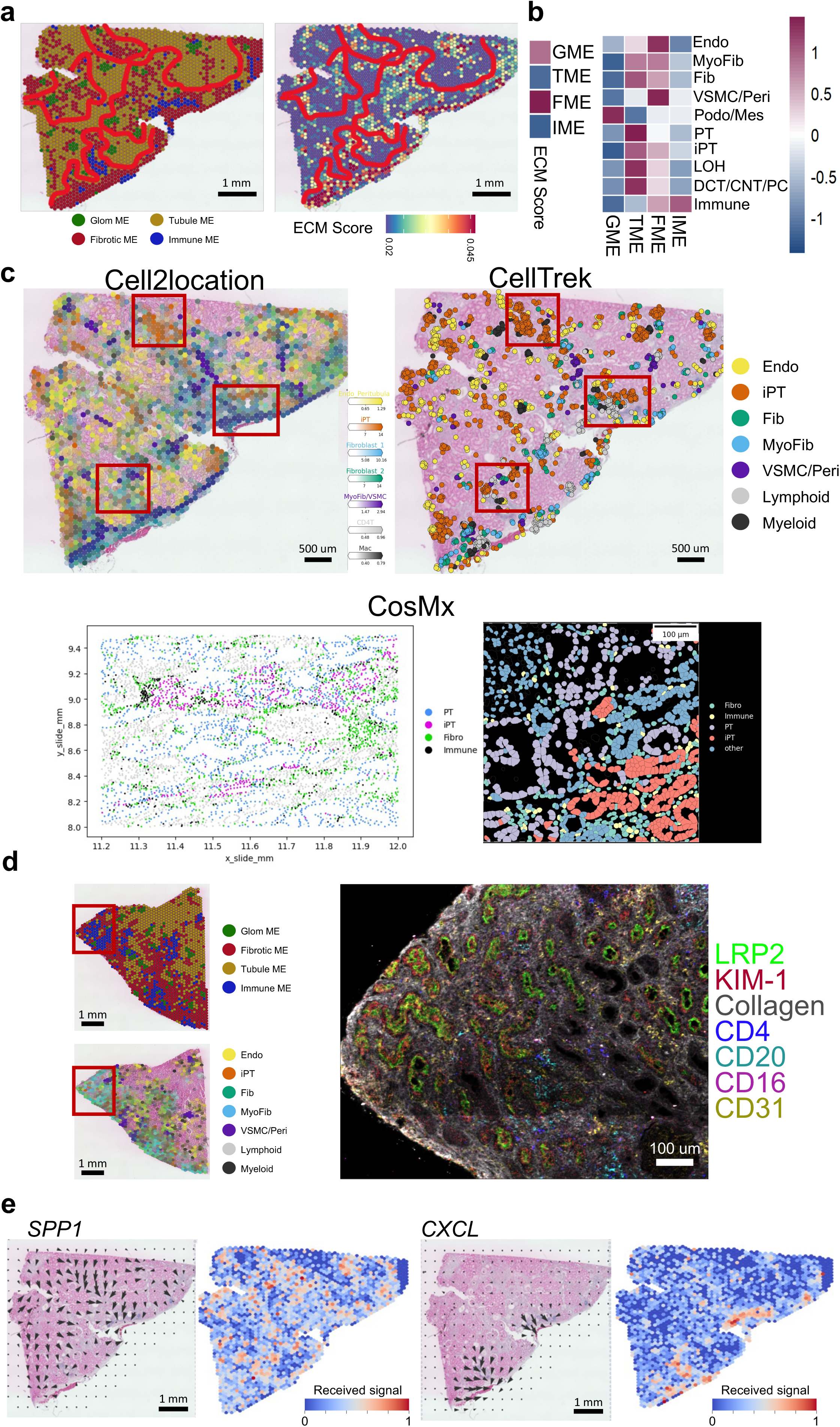
Human kidney fibrotic microenvironment. **(a)** Human kidney microenvironments defined by non-negative matrix factorization (NMF) of the spRNAseq. Glomerular (green), tubule (brown), fibrotic (red), immune (blue). The spatial distribution of the calculated ECM score (right panel). Red indicates higher ECM gene expression. **(b)** Correlation of microenvironment with ECM score (left) and cell type correlation with each microenvironment (right). **(c)** Cell2location (left), Celltrek (right) cell type imputations of the Visium spatial data and CosMx (below) annotations (below) showing the location of different cell types in the fibrotic microenvironment in a diseased kidney sample. CosMx annotations of iPT, PT, fibroblasts and immune cells are shown at 2 different magnifications. **(d)** *In situ* mass spec image of CKD kidney (from panel E) labeled with LRP2 (PT) KIM1 (iPT), CD4 (T-cell) CD20 (B-cell), CD16 (myeloid) and CD31 (endo) in a fibrotic human kidney sample. **(e)** The spatial location of the identified cell-cell interaction pathway (*SPP1* and *CXCL*). The arrows indicate the source and targets of the identified pathways, and the color indicates the received signals (red higher). The results were obtained using COMMOT package.

We also identified a specific immune ME. The immune ME regions were located within the FME, but with patchy distribution. The immune ME consisted of dendritic cells, plasma cells, and B-and T-lymphocytes (**Fig. 4b, c**, **Supplementary** Fig. 26**, 27**). The immune ME organizations resembled early tertiary lymphoid structures^41^. Immunostaining studies with cell type specific antibodies validated the presence of these specific immune cells and immune cell aggregates (**Supplementary** Fig. 28). Furthermore, we have performed *in situ* mass spectrometry (IMC) to understand protein expression levels. Consistent with the Visium data, we confirmed the protein expression of HAVCR1+-positive iPT cells, B-cells, CD4T-cells, plasma, myeloid and endothelial cells by IMC (**Fig. 4d**, **Supplementary** Fig. 29).

To further understand cell interactions in FMEs, we implemented CellChat^42^ on the integrated sc/snRNAseq and snATACseq datasets as well as communication analysis by Optimal Transport (COMMOT)^43^ on the spRNAseq datasets. We found enrichment for *SPP1*, *CXCL12*, *CCL19*, *CCL21, PDGFB,* and *TGFB1* and their receptors in FME regions (**Fig. 4e, Supplementary** Fig. 30**, Supplementary Table 10**). The iPT cells in the FMEs expressed *SPP1* and *PDGFB* and showed a strong interaction with stromal cells. Also, we found endothelial cells expressing *CD34* and *CDH5* in the FME regions (**Supplementary** Fig. 30). The stromal cells in FME were enriched for chemotactic factors including *CXCL12*, *CCL19*, *CCL21*, while their receptors were expressed in different immune cells, suggesting that stromal cells might signal to immune cells. We observed *PDGFB* and *TGFB1*, known mediators of fibrosis, in FME associated immune aggregates (**Supplementary** Fig. 30)^43^.

Overall, using unbiased computational tools, we were able to identify spatial regions in the kidney, including glomerular, tubular regions in healthy samples and fibrotic and immune regions in diseased kidneys. Most importantly, FMEs were not only characterized by matrix-producing fibroblasts, but also associated iPT-stromal ligand-receptor interactions.

### Injured tubule cells in human kidney fibrosis

Cell type enrichment analysis of the fibrotic microenvironment indicated fibroblast, myofibroblast, immune cells, endothelial cells and injured tubule cells including injured proximal tubule cells (iPT) in FMEs (**Fig. 4e**). iPT cells have not been well characterized, but they have been identified in diseased mouse and human kidney samples ^10,11,44,45^. iPT cells were physically close to proximal tubule cells **(Fig. 5a, 4c, Supplementary** Fig. 33). Further, we note that PT cells number correlated with eGFR, fibrosis and disease state within our cohort (**Fig 5b, c**) and showed the most drastic decline in CKD **(Supplementary** Fig. 32). In contrast, we observed a clear enrichment of iPT cells in the fibrotic microenvironment and increased number in diseased samples **(Fig. 5d).** Upon careful sub-clustering, of our CosMx data, which we verified by examining our annotations in space, We note that *VCAM1* expression in both PEC and iPT, while *HAVCR1* was specific for iPTs **(Fig. 5e, Supplementary** Fig. 20a).

**Fig 5.**
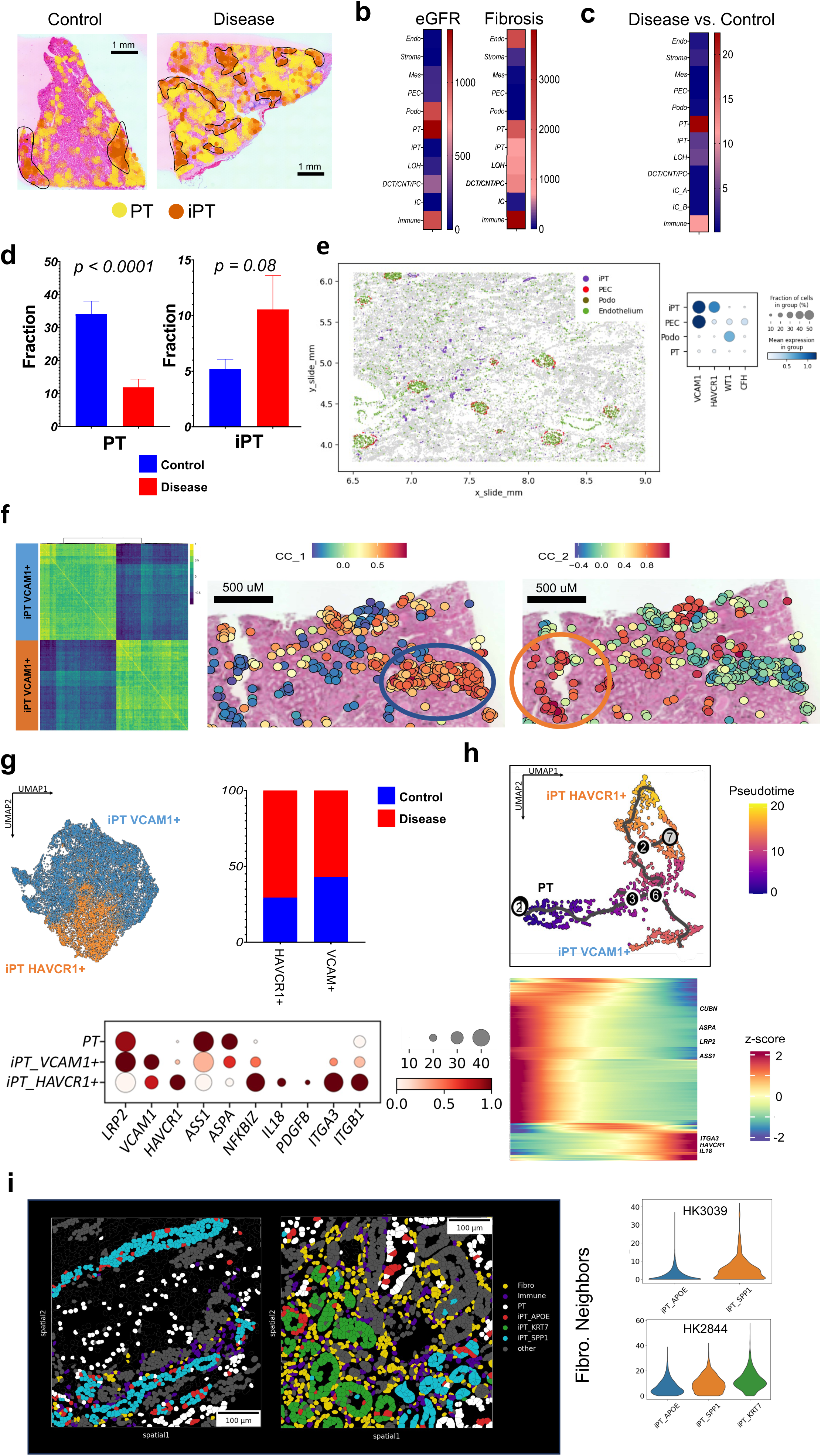
Injured proximal tubule cells in diseased human kidneys. **(a)** Localization of iPT and PT cells within control and diseased samples in the human kidney. iPT is scattered in a patchy pattern, and colocalize amongst themselves. **(b)** Heatmap of number of genes correlated with eGFR and renal fibrosis in each cell type. The red shows higher number of genes and blue indicates lower number of genes. **(c)** Heatmap of number of genes comparing prevalence in healthy vs. diseased samples in each cell type. PT had the highest changes between control and diseased samples, with red indicating more genes. **(d)** The fractions of PT and iPT in control and diseased samples type in the integrated sn/scRNAseq and snATACseq dataset. Bar indicated the SEM. For the comparison between two groups, t-test was used. **(e)** CosMx spatial transcriptomics shows that annotation of iPT and PECs localize to tubules and glomeruli, respectively. Dot plot shows *VCAM1* is expressed in both populations, while *HAVCR1* and *CFH* appear to be more specific for iPT and PECs, respectively. **(f)** Gene co-expression network analysis of human kidney spRNAseq data indicates two modules expressing *VCAM1* or *HAVCR1*. The right panels indicate the spatial location of *VCAM1* or *HAVCR1* (KIM1) positive injured PT cells. The color indicates gene expression of iPT modules (red higher expression). **(g)** Sub-clustering of iPT cells from integrated atlas. The fraction of VCAM1 or HAVCR1 positive iPT cells in control and diseased kidneys, shown on UMAP and quantified in frequency. The dot plots show the expression marker genes in iPT and PT cells. The size of the dot indicates the percent positive cells, and the darkness of the color indicates average expression. **(h)** Cell trajectory analysis (Monocle) representation of PT and iPT cells (Upper panels). The heatmap shows the differentially expressed genes along the trajectory, cells ordered by pseudotime. Red indicates higher expression. **(i)** Sub-clustering of iPT within the CosMx data shows 3 subtypes of iPT. These iPT subtypes are located near PT cells and are visualized in specific fields of view—one in each sample. These iPT subtypes have different neighboring cells (within a 50 micron radius) with iPT_KRT7 having most frequent fibroblast and immune neighbors within our diseased sample (*p*-value = 1e-177, 1e-17 for fibroblasts and immune cells respectively within our disease sample by Wilcoxon rank sum test). iPT_APOE had fewer immune (*p*-value = 0, *p*-value = 0, for HK3039 and HK2844) and fibroblast neighbors than the iPT_SPP1 subtype. (*p*-value = 5e-52, *p*-value = 1e-52, for HK3039 and HK2844). Endo; endothelial cells Mes; mesangial cells, Stroma; stromal cells, PEC; parietal epithelial cells, Podo; podocyte, PT; proximal tubule cells, iPT; injured proximal tubule cells, LOH; loop of Henle, DCT; distal convoluted tubule, CNT; connecting tubule cells, PC; principal cells of collecting duct, IC_A; Type alpha intercalated cells, IC_B; Type beta intercalated cells.

We then sought to examine iPT heterogeneity, starting with our Visium data. We used the single cell co-expression (SCoexp) module of CellTrek^33^. We identified two different iPT modules, corresponding to two iPT subtypes in diseased samples (**Fig. 5f**) and only one iPT type in healthy samples (**Supplementary** Fig. 34). Moving back to the rich integrated atlas, we found that one of the iPT cluster expressed *VCAM1*, *ACSL1*, *ASS1*, and *ASPA*; genes playing role in cellular metabolism. We called this cluster iPT_VCAM1+. This cluster was present in healthy samples and this cluster was close to other injured PT cells in the integrated dataset expressing mitochondrial genes such as *MT1* (**Fig. 5f**). The second iPT cluster expressed *HAVCR1* (or *KIM1*), *NFKBIZ*, *IL18*, *ITGA3*, *PDGFB*, and *ITGB1* and was enriched for the expression of genes associated with cell adhesion and matrix (iPT-HAVCR1+) (**Supplementary** Fig. 34). Both in the Visium spatial and dissociated datasets, most iPT-HAVCR1+ cells were in CKD samples (**Fig. 5g**). Pseudo-time analysis of PT and iPT cells using Monocle, indicated a continuous differentiation path of PT cells to iPT_VCAM1+ and iPT_HAVCR1+ (**Fig. 5h, Supplementary Table 11)**. Studies going back to a decade ago have identified HAVCR1+ as an injured tubule marker^10^. These two different iPT cell subtypes were also present in a recently published mouse DKD snRNAseq^45^ dataset (**Supplementary** Fig. 35) as well as in our snATACseq data (**Supplementary** Fig. 36). We identified *HNF1A* and *BACH2* as enriched TFs for iPT_VCAM1+ and iPT_HAVCR1+, respectively (**Supplementary** Fig. 36).

We then subclustered iPT cells using our positionally verified CosMx data (Fig. 5i, Supplementary Fig. 31, 34). We noted upon sub-clustering iPT heterogeneity that corresponded with neighboring cell identity (Supplementary Fig. 31), supporting our Visium data that iPT subpopulations co-localize and may be related to the underlying microenvironment. In the CosMx dataset, APOE, SPP1 and KRT7 (**Supplemental Figure 31b**) marked these subclusters. Our identification of these specific markers, as opposed to other iPT markers, may be related to the relative paucity of transcripts used by the CosMx platform compared to whole transcriptome approaches.

In summary, the different types of single cell omics (snRNA, scRNA, snATAC) data indicated highly plastic PT cell population, including iPT cells. Our spatial transcriptomics analysis highlighted different types of iPT cells (*VCAM1*+ and *HAVCR1*+). We observed *VCAM1*+ iPT and PEC cells in non-CKD adult kidney samples, while HAVCR+ cells were enriched in the fibrotic microenvironment and diseased kidneys.

### Fibrotic microenvironment gene signature successfully predicts disease prognosis in a large cohort of human kidney samples

At present, kidney disease classification is performed by an expert renal pathologist. We wanted to understand whether our spRNAseq can aid disease classification and prognosis evaluation. We first generated an FME gene signature (FME-GS) (**Supplementary Table 12**) and used this signature to analyze gene expression data from a large external kidney cohort containing 292 human kidneys (**Fig. 6a**), including healthy samples and samples with varying severity of CKD (attributed to diabetes and hypertension). Our FME-GS successfully discriminated samples with more severe disease (higher fibrosis and lower eGFR) from healthier samples (**Fig. 6b, Supplementary Table 13**) despite the fact, that these parameters were not included in the clustering algorithm. Next, we wanted to understand whether FME signature can predict kidney disease prognosis. We found that cluster 1 subgroup, samples with the highest FME-GS load, had 4x higher chance of reaching endpoint, defined as 40% change in eGFR or reaching end stage renal failure (eGFR< 15 ml/min) **(Fig. 6c)**. Subgroup analysis however, indicated that samples with the highest FME-GS has statistically higher degree of fibrosis 12.1% vs 6.51%. While this difference is not clinically meaningful the differences was statistically significant.

**Fig 6.**
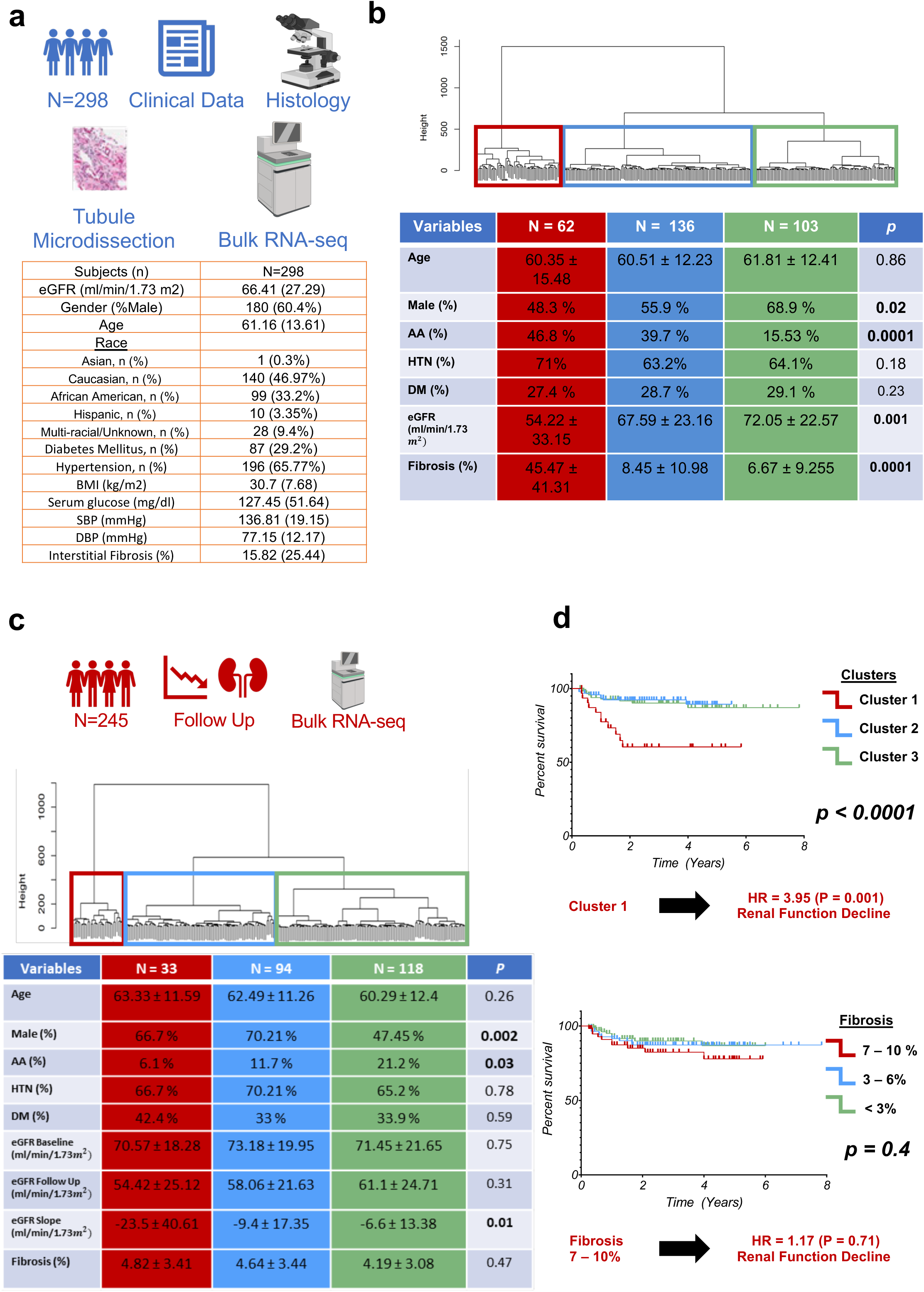
Fibrotic microenvironment gene signature successfully predicts disease prognosis in a large cohort of human kidney samples. **(a)** Clinical characteristics of 292 human kidney tubule RNA samples. **(b)** Unbiased cluster dendrogram of 292 human kidney tubule bulk RNAseq samples based on FME gene signature. Clinical characteristics of each cluster. Chi-square test for categorical variables and one-way ANOVA for continuous variables were used to compare groups. **(c)** Unbiased cluster dendrogram of 245 human kidney tubule bulk RNAseq samples with fibrosis <10% based on expression of FME genes. Clinical characteristics of each cluster are shown in the table. Chi-square test for categorical variables and one-way ANOVA for continuous variables were used to compare groups. **(d)** Kaplan-Meier analysis of 292 kidney samples based on FME gene signature. Renal survival was defined as cases reaching end stage renal disease or greater than 40% eGFR decline (above). Kaplan-Meier analysis of 245 kidney samples based on pathologist defined kidney fibrosis degree (<3%, 3-6% or 7-10% as defined by an expert pathologist) (below). Renal survival was defined as cases reaching end stage renal disease (eGFR of 15 ml/min/1.73m2) or greater than 40% eGFR decline. FME-GS; fibrotic microenvironment gene signature, HR; hazard ratio, HTN: hypertension, DM: diabetes, AA: African American.

Finally, we wanted to understand whether the FME signature could also predict renal disease progression in patients with preserved eGFR (mean eGFR 70 ml/min/1.73*m*^!^) and no structural damage (interstitial fibrosis less than 10%). These parameters are used to define the lack of structural and functional kidney damage (CKD/fibrosis)^46^ **(Fig. 6c)**. The pathologist reported fibrosis score could not predict the outcome for these samples (**HR = 1.17, 95%CI: 0.49 – 2.83**) **(Fig. 6d)**. Kaplan-Meier analysis indicated that cluster 1, with the highest FME-GS load, had significantly higher hazard ratio to reach the endpoint (**HR = 3.95, 95%CI: 1.71 – 9.12**) **(Fig. 6d)**. While the entire FME gene set better predicted outcome, employing only the top 10 genes from LASSO regression still showed a statistically significant enrichment for renal failure (**Supplementary** Fig. 37**, p=0.021, Supplementary Table 14**). Clustering samples using a random gene set did not correlate with outcome (**Supplementary** Fig. 37). Analyzing the relationship between cell types and kidney disease progression, we found that genes that correlated with eGFR slope were enriched in iPT and stromal cells, potentially implying their causal role (**Supplementary** Fig. 37).

In summary, our spatially derived FME-GS can identify subjects with progressive kidney function decline in a large cohort.

## Discussion

Here we present the spatial molecular principles of kidney health and disease by generating a comprehensive and spatially resolved human kidney atlas and combining single cell omics data (scRNAseq, snRNAseq, and snATACseq) from a large number of human kidney tissue samples with varying degree of disease severity. Our work fills a critical knowledge gap by characterizing the cellular gene expression program of cells previously only defined by their spatial location, showing the anatomical location of cells only observed in dissociated single cell expression data, and defining cell-type specific gene expression changes in diseased states. We define the cellular complexity of the fibrotic microenvironment as the intricate interaction of many different cell types. We furthermore demonstrate the clinical prognostic value of spatial transcriptomics even for samples where the current pathological analysis loses sensitivity.

Previous single cell analyses, focusing on dissociated human and mouse kidney datasets, have generated gene expression and regulatory matrices for a variety of kidney cell types^8–12,47^. As kidney cell types have been functionally well characterized, most identified cell types have been matched back to a more than half-century old functional cell type definition^6^. A key limitation of these analyses has been the identification and molecular characterization of anatomically defined cell types, such as mesangial cells, PECs, and fibroblasts. Here, we demonstrate that a joint approach, which combines single cell and single nuclear expression, open chromatin, and single cell and spot level spRNAseq data from many diverse samples and a large number of cells, is critical to achieving this goal. The orthogonal analytical tools provide unique opportunities for validation, as each method suffers from specific technological biases. Here, we have not only been able to resolve and validate previously anatomically-known cell types but also identify novel cell types such as specific stromal cells for glomerulosclerosis (expressing *CDH13)*^48^. Furthermore, we clarify genes expressed by cells previously only identified by their anatomical location, such as PECs.

Fibrotic diseases are responsible for close to 40% of all deaths^49^. Kidney fibrosis is the final common pathway to end stage kidney failure^50^. Combining snRNA and spatial information and multiple datasets we characterize stromal cell subtypes, and we validate their consistency between different datasets. We could conclusively discriminate VSMC and mesangial cells from myofibroblasts that are anatomically distinct but share gene expression signatures in dissociated sc/snRNAseq data^51–53^. We could identify novel markers and, ultimately, new stromal cell types and determine their spatial location. This information could be important in the field of finding therapeutic candidates for renal fibrosis. We noted a cluster of FAP-positive fibroblasts in diseased human kidneys^54–56^. FAP targeted cellular and RNA therapies have been developed and shown to have efficacy in animal models of cardiac fibrosis^54–56^. Our data suggests that these therapeutics may be helpful for treating kidney fibrosis.

Here, we demonstrate the cellular and architectural complexity of kidney fibrosis. We propose the use of the fibrotic microenvironment to characterize these lesions, to not only focus on matrix accumulation but on the elaborate cellular complexity of these lesions. We show that FME signature contains tubule cells including injured PT. While iPT cells have been observed in dissociated single cell data, here we show that these cells can be consistently identified in all datasets and contain consistent markers. We furthermore identify spatially defined iPT subtypes ^10^. We show that in humans *VCAM1*+ cells can be observed in subjects with preserved kidney function. This might be consistent with observations that we show that some parietal epithelial cells express VCAM1 or that low degree of fibrosis is observed in some healthy subjects or during aging. We identify HAVCR1 positive iPT cells that almost exclusively present in CKD samples. *HACVR1* has been identified as marker of acute and chronic kidney disease.^57^

Immune cell clusters have long been observed in fibrotic kidney samples, even in patients with non-immune-mediated kidney disease, such as diabetes and hypertension^50,58^. Here, we resolve these regions both spatially and at a cell type level. The interaction of a large number of cell types, including iPT, immune, stromal, and endothelial cells, establishes the FME. In patient samples, there is a strong interaction between stromal and immune cells and also signaling by immune and stromal cells to iPT, which might play a role in maintaining their injured PT state. Our kidney scRNAseq data was enriched for immune cells and enabled us to spatially resolve immune cell types and determine the distributions of immune cells in the kidney. We show that immune cell clusters (the immune microenvironment) are localized mostly within some FMEs and to some degree they show a resemblance to the tertiary lymphoid structures (TLS). TLS are organized aggregates of immune cells that form postnatally in nonlymphoid tissues, usually as a persistent antigen production^59^, and generate autoreactive effector cells. TLS have been earlier described in mouse kidney tissue samples^60–63^.

One of the most devastating complications of CKD is its progression to ESRD, which requires life-sustaining dialysis or transplantation ^64^. Predicting which patients will have progressive disease is of major clinical importance. This has been classically done using eGFR and the extent of fibrosis noted on biopsy for subjects with more advanced disease (eGFR<60 ml/min/1.73m2). While some studies indicated that including the degree of tissue fibrosis can improve kidney prognosis estimates, this has not been uniformly observed. By using the FME-GS score, we were able to improve upon this. We identified subjects at risk of ESRD in a large external dataset of human kidney tissue samples even for samples where traditional fibrosis score had limited predictive value. These results establish FME-GS as a key biomarker and potentially as a causal mechanism of progression. Future studies shall work on optimizing and validating the predictive gene signature, but our preliminary data indicates that a relatively small number of genes might be able to discriminate those patients who progress to renal failure.

Our study has several limitations. Here, we present the initial interpretation of one of the largest dissociated and spatially resolved human kidney datasets. Here, we used the clinical and histological annotation for disease definition, however, hypertensive kidney disease might not be a single disease entity, where hypertension might not even be the single causal injury. While we applied a variety of tools and methods, future studies will be essential for a more comprehensive analysis of the presented information. Our datasets indicate the presence and key role of injured PT cells and immune cells in disease progression; however, their functional role should be studied in mechanistic experiments using cells and animal models. Furthermore, the FME gene signature will need to be validated in other cohorts.

In summary, we develop a spatially defined molecular human kidney cellular atlas, characterize the fibrotic microenvironment, and indicate its role as a clinically meaningful prognostic disease biomarker, demonstrating the utility of spRNAseq for the investigation of complex diseases.

## Methods

### Single nuclei RNA sequencing

Nuclei were isolated using lysis buffer (Tris-HCl, NaCl, MgCl2, NP40 10%, and RNAse inhibitor (40 U/ul)). 10-30 mg of frozen kidney tissue was minced with a razor blade into 1-2 mm pieces in 1 ml of lysis buffer. The chopped tissue was transferred into a gentleMACS C tube, and homogenized in 2 ml of lysis buffer using a gentleMACS homogenizer with programs of Multi_E_01 and Multi_E_02 for 45 seconds. The homogenized tissue was filtered through a 40 µm strainer (08-771-1, Fisher Scientific) and the strainer was washed with 4 ml wash buffer. Nuclei were centrifuged at 500xg for 5 minutes at 4°C. The pellet was resuspended in wash buffer (PBS 1X + BSA 10% (50 mg/ml), + RNAse inhibitor (40 U/ul)), filtered through a 40 µm Flowmi cell strainer (BAH136800040-50EA, Sigma Aldrich). Nuclear quality was checked, and nuclei were counted. In accordance with the manufacturer’s instructions, 30,000 cells were loaded into the Chromium Controller (10X Genomics, PN-120223) on a Chromium Next GEM chip G Single Cell Kit (10X Genomics, PN-1000120) generate single cell gel beads in the emulsion (10X Genomics, PN-1000121). The Chromium Next GEM Single Cell 3ʹ GEM Kit v3.1 (10X Genomics, PN-1000121) and Single Index Kit T Set A (10X Genomics, PN-120262) were used in accordance with the manufacturer’s instructions to create the cDNA and library. Libraries were subjected to quality control using the Agilent Bioanalyzer High Sensitivity DNA kit (Agilent Technologies, 5067-4626). Libraries were sequenced using the Illumina Novaseq 6000 system with 2 × 150 paired-end kits. Demultiplexing was as follows: 28 bp Read1 for cell barcode and UMI, 8 bp I7 index for sample index, and 91 bp Read2 for transcript.

### Single nuclei ATAC sequencing

The procedure described above for single nuclei RNA sequencing was used to isolate the nuclei for ATAC sequencing. The resuspension was performed in diluted Nuclei Buffer (10X GEM). Nuclei quality and concentration were measured in Countess AutoCounter (Invitrogen, C10227). Diluted nuclei were loaded and incubated in chromium single cell ATAC library & gel bead kit’s transposition mix (10X Genomics, PN-1000110). Chromium Chip E (10X Genomics, PN-1000082) in the Chromium Controller was utilized to capture the GEMs. The Chromium Single Cell ATAC Library & Gel Bead Kit and Chromium i7 Multiplex Kit N Set A (10X Genomics, PN-1000084) were then used to create snATAC libraries in accordance with the manufacturer’s instructions. Library quality was examined using an Agilent Bioanalyzer High Sensitivity DNA kit. After sequencing on an Illumina Novaseq system using two 50 bp paired-end kits, libraries were demultiplexed as follows: 50 bp Read1 for DNA fragments, 8 bp i7 index for sample index, 16 bp i5 index for cell barcodes, and 50 bp Read2 for DNA fragments.

### Single Cell RNAseq

Fresh human kidneys (up to 0.5 gr) collected in RPMI were minced into approximately 2-4 mm cubes using a razor blade. The minced tissue was then transferred to a gentleMACS C tube containing Multi Tissue dissociation kit 1 (Miltenyi, #130-110-201). The tissue was homogenized using the Multi_B program of the gentleMACS dissociator. The tube, containing 100ul of Enzyme D, 50ul of Enzyme R, and 12.5ul of Enzyme A in 2.35 ml of RPMI, was incubated for 30 mins at 37 °C. Second homogenization was performed using Multi_B program on gentleMACS dissociator. The solution was then passed through a 70 um cell strainer. After centrifugation at 1,200 RPM for 7 mins, cell pellet was incubated with 1ml of RBC lysis buffer on ice for 3mins. The reaction was stopped by adding 10 ml PBS. Next, the solution centrifuged at 1,000 RPM for 5 minutes. Finally, after removing the supernatant, the pellet was resuspended in PBS. Cell number and viability were analyzed using Countess AutoCounter (Invitrogen, C10227). This method generated a single cell suspension with greater than 80% viability. Next, 30,000 cells were loaded into the Chromium Controller (10X Genomics, PN-120223) on a Chromium Next GEM chip G Single Cell Kit (10X Genomics, PN-1000120) to generate single cell gel beads in the emulsion (GEM) according to the manufacturer’s protocol (10X Genomics, PN-1000121). The cDNA and library were made using the Chromium Next GEM Single Cell 3ʹ GEM Kit v3.1 (10X Genomics, PN-1000121) and Single Index Kit T Set A (10X Genomics, PN-120262) according to the manufacturer’s protocol. Quality control for the libraries was performed using Agilent Bioanalyzer High Sensitivity DNA kit (Agilent Technologies, 5067-4626). Libraries were sequenced on Illumina Novaseq 6000 system with 2 × 150 paired-end kits using the following demultiplexing: 28 bp Read1 for cell barcode and UMI, 8 bp I7 index for sample index and 91 bp Read2 for transcript.

### Visium FFPE sample preparation

The RNA quality of human kidney FFPE sample was checked by RNeasy FFPE kit (Qiagen-Cat #73504) according to the manufacturer’s protocol. RNA quality was examined using Agilent bioanalyzer, and samples with DV200>50% were selected. Then a 5 µm tissue sample was cut onto the 10x Visium Spatial Gene Expression Slide. After deparaffinization, H & E staining was performed. We used Keyence 1266 BZ-X810 microscope for whole slide imaging. After scanning, de-crosslinking, probe hybridization, probe release, and extension, library preparation was performed using a Single Index Kit TS Set A (10X Genomics, PN-3000511) according to the manufacturer’s protocol. Quality control for the libraries was performed using the Agilent Bioanalyzer High Sensitivity DNA kit (Agilent Technologies, 5067-4626). Libraries were sequenced using the Illumina Novaseq 6000 system with 2 × 150 paired-end kits. Demultiplexing was as follows: 28 bp Read1 for cell barcode and UMI, 10 bp I7 index, 10bp i5 index, and 50 bp Read2 for transcript.

### CosMx Sample preparation

Samples were prepared according to manufacturer specifications. Tissue sections were cut at 5 um thickness and prepared according to the manufacturer specifications (NanoString Technologies). We used the human universal cell characterization RNA probes, plus 50 additional custom probes which were designed to the following gene transcripts: *ESRRB, SLC12A1, UMOD, CD247, SLC8A1, SNTG1, SLC12A3, TRPM6, ACSL4, SCN2A, SATB2, STOX2, EMCN, MEIS2, SEMA3A, PLVAP, NEGR1, SERPINE1, CSMD1, SLC26A7,SLC22A7, SLC4A9, SLC26A4, CREB5, HAVCR1, REN, AP1S3, LAMA3, NOS1, PAPPA2, SYNPO2, RET, LHX1, SIX2, CITED1, WNT9B, AQP2, SCNN1G, ALDH1A2, CFH, NTRK3, WT1, NPHS2, PTPRQ, CUBN, LRP2, SLC13A3, ACSM2B, SLC4A4, PARD3, XIST,UTY*. Samples were imaged with configuration A. After imaging, the flowcell was kept in xylene overnight, the coverslip was removed and the slide was stained with hematoxylin and eosin.

### Human Sample Acquisition

Left-over kidney samples were irreversibly deidentified, and no personal identifiers gathered. Therefore, they were exempt from IRB review (category 4). We engaged an external honest broker who was responsible for clinical data collection without disclosing personal identifiable information. The University of Pennsylvania institutional review board (IRB) gave its approval for the collection of human kidney tissue.

Cortical and outer medullary area of the human kidney was used for scRNAseq, snRNAseq, and snATACseq. In addition, a portion of the tissue was formalin-fixed, paraffin-embedded, and stored for spatial trascriptomics, as well as stained with periodic acid-Schiff for pathology scoring. A local renal pathologist performed objective pathological scoring.

### Immunostaining

Paraffin blocks were sectioned and deparaffinized. For blocking, 1% bovine serum albumin was used. Slides were then incubated overnight with diluted primary antibodies (CD4 CST [Catalogue #48374], IGKC: Biolegend [Catalogue #392702], and CD79A Abcam [Catalogue #ab79414]). After washing the sections with PBS three times, secondary antibodies were applied for 1 hour at room temperature. After DAPI staining, slides were imaged with an OLYMPUS BX43 Microscope. Positive cells in ten randomly selected fields were counted on each slide.

### Imaging Mass Spectrometry

Formalin-fixed paraffin-embedded (FFPE) tissue sections were deparaffinized and rehydrated using standard protocols. Antigen retrieval was performed using citrate buffer (pH 6.0), and sections were blocked with a buffer containing 5% bovine serum albumin (BSA). Primary antibodies against target proteins were conjugated with specific metal isotopes (mention which isotopes) using the Maxpar Antibody Labeling Kit (Fluidigm). Tissue sections were incubated with the antibody-metal conjugates overnight at 4°C. After washing, secondary antibodies, conjugated with a different metal isotope (mention which isotopes), were added to amplify the signal. Finally, the tissue sections were stained with an iridium intercalator for nuclear staining. IMC imaging was performed using a Hyperion imaging system (Fluidigm) equipped with a 20X objective lens. The tissue sections were ablated using a laser at a 1μm pixel size, and the resulting metal signals were detected by a time-of-flight mass spectrometer.

## Bioinformatic analysis

### Primary single nuclei and cell RNAseq data processing

FASTQ files from each 10X single nuclei run were processed using Cell Ranger v6.0.1 (10X Genomics). Gene expression matrices for each cell were produced using the human genome reference GRCh38.

### Data Processing and Computational Analyses

Ambient RNA was corrected using SoupX^65^ and doublets were removed by DoubletFinder^66^ with default parameters. Seurat objects were created from the aligned outputs of multiple samples, retaining genes expressed in more than three cells and cells with at least 300 genes. Further, a merged Seurat object was obtained using the merge function in Seurat v4.0.366. The QC filters applied were: (a) cells with n_feature counts more than 3000 or less than 200, (b) cells with more than 15% mitochondrial counts (for snRNAseq data) and more than 50% mitochondrial counts (for scRNAseq data), and (c) cells with more than 15% ribosomal gene counts.

### Data Normalization and Cell Population Identification

Initially, highly variable genes were identified using the vst method. The data was then natural log-transformed and scaled. The scaled values underwent principle component analysis (PCA) for linear dimension reduction. We used the RunHarmony function in the harmony^67^ package for batch effect correction. A shared nearest neighbor network was created based on Euclidean distances between cells in a multidimensional PC space (the first 50 PCs were used for both snRNAseq and scRNAseq). This network was used to generate a 2-dimensional Uniform Manifold Approximation and Projection (UMAP) for visualization.

To identify cell-type markers, we used the FindAllMarkers function in Seurat. This method calculates log fold changes, percentages of expression within and outside a group, and p-values of the Wilcoxon-Rank Sum test comparing a group to all cells outside that specific group, including adjustment for multiple testing. A log-fold-change threshold of 0.25 and FDR<0.05 was considered significant. These steps were performed on the snRNAseq and scRNAseq datasets separately. Clusters expressing multiple cell types of specific marker genes were excluded as potential doublets. The dataset was further refined: first, clusters were annotated based on well-known marker genes, then the dataset was projected to a published human kidney single cell^44^ reference using the CCA method. The FindTransferAnchors function was used to discover matching genes between the two datasets by using shared correlation patterns in the gene activity matrix and snRNAseq dataset. Predicted labels within the two datasets were identified using the TransferData method. Cells were considered doublets and eliminated from the dataset if the original annotation was incompatible with the predicted cell type.

### Single nuclei ATACseq analysis

Raw FASTQ files were aligned to the GRCh38 reference genome and quantified using Cell Ranger ATAC (v. 1.1.0). `O`utputs `from 19` snATACseq datasets were embedded using Signac (v.1.3.0)^68^ to generate Signac object. Low-quality cells were removed from each snATAC object using the following criteria: peak_region_fragments < 3000 & peak_region_fragments > 20000 & pct_reads_in_peaks < 15 & nucleosome_signal > 4 & TSS.enrichment < 2). The filtered cells in `19` objects were merged in Seurat. Dimension reduction was `performed` by singular value decomposition (SVD) of the TFIDF matrix and UMAP. `B`atch effect was corrected using Harmony^67^ via the RunHarmony function in Seurat. A KNN graph was `created` to cluster cells using the Louvain algorithm. Peaks observed in at least 20% of cells were evaluated for differentially accessible chromatin regions (DARs) between different cell types using a likelihood ratio test, a log-fold-change threshold of 0.25, and an FDR of 0.05 with the Signac FindMarkers function. To annotate the genomic regions harboring snATACseq peaks, ChIPSeeker (v1.24.0)70 was used^69^.

### Removing the doublets from snATACseq dataset

The GeneActivity tool in Signac was used to create a gene activity matrix after clustering the twenty integrated snATACseq datasets using protein-coding genes annotated in the Ensembl database (this technique counts the ATAC peaks inside the gene body and 2 kb upstream of TSS). Log normalization was applied to the gene activity matrix.

A published snRNAseq^44^ dataset was utilized as a reference, and the FindTransferAnchors function was used to discover matching genes between the snRNAseq and snATACseq datasets by using shared correlation patterns in the gene activity matrix and snRNAseq dataset. Subsequently, the predicted labels within two datasets were identified using the TransferData method. If the original annotation was incompatible with the predicted cell type, the cell was considered a doublet and was eliminated from the dataset.

### Motif Enrichment Analysis and Motif Activities

First, we used the AddMotifs function of Signac to perform motif enrichment analysis. This was done by creating a matrix of positional weights for motif candidates from JASPAR2020. To identify potentially important cell-type-specific regulatory sequences and motifs, we evaluated the differentially accessible peaks between each cell type versus other cell types using the ‘LR’ test and min.pct = 0.05. Subsequently, a hypergeometric test was used to determine the likelihood of observing a specific motif at a particular frequency by chance. We compared this observation with a group of peaks that have the same GC content as the motif, which we considered the background set. This analysis was performed using the Find Motifs function. We used the RunChromVAR function in conjunction with chromVAR (v.1.6.0) ^70^ to ascertain transcription factor activity.

### DARs between groups

We used the FindMarkers function after selecting DefaultAssay as peaks to identify DARs in each cell type and diseased and healthy conditions, with a log-fold-change threshold of 0.25 and FDR<0.05. Peaks translated to related genes using ChIPSeeker (v1.21.1)^69^.

### Integration of snRNAseq, scRNAseq, and snATACseq datasets

We used SCVI^30,71^ to integrate scRNAseq, snRNAseq, and snATACseq data from multiple samples. Briefly, we used the preprocessed and clean datasets previously described and then combined them into a single AnnData object using the Scanpy v1.4.4 package. After normalization and identification of the 3000 highly variable genes in the integrated dataset, we ran SCVI on the combined dataset to learn a joint latent representation of the data, using the sample, nFeature_RNA, nCount_RNA, and tech factors as covariates. We utilized the scanorama setting of SCVI to account for batch effects across samples. We incorporated four layers in the model and reduced the dimensionality of the data to 45 dimensions using the SCVI’s dimensionality reduction method. Finally, we used the Scanpy package to perform downstream analysis on the integrated dataset, including clustering, differential expression analysis, and visualization.

### Ligand–receptor interactions in integrated sc/snRNAseq and snATACseq

The CellChat^42^ repository (v1.4.00 was used to assess cellular interactions between different cell types and to infer cell–cell communication networks from integrated sc/snRNAseq and snATACseq data. Only receptors and ligands expressed in more than 10 cells in each cluster were considered. Probability and *P* values were measured for each interaction.

### DEGs between diseased and healthy groups

To identify DEGs between groups, we utilized the FindMarkers tool for each cell type and condition, a log-fold-change threshold of 0.25, and an FDR 0.05. DEGs in the integrated dataset of snRNAseq, scRNAseq, and snATACseq were calculated using model.differential_expression of SCVI tool with the default parameters.

### Single nuclei RNAseq trajectory analysis

Proximal tubules (PT) and Injured PT cells were subclustered for the trajectory analysis. The trajectory analysis was done in two steps. Different sub-types of iPT cells with equal numbers were randomly subsampled and cell dataset object (CDS) was generated using Monocle3^72,73^. After preprocessing and batch effects correction, the dataset was embedded for dimension reduction and pseudotemporal ordering. We used the order_cell function and indicated the PT as start point for pseudotime analysis. The track genes algorithm was used to identify the DEGs along the trajectories, and genes with q values of 0.05 or higher were considered significant.

## MetaNeighbor analysis

MetaNeighbor^37^ is a versatile computational tool designed to quantify the degree of cell type replicability between datasets by assessing cell type-specific gene expression patterns. The method involves constructing a cell-cell similarity network using a shared, reduced gene space and subsequently calculating the area under the receiver operating characteristic curve (AUROC) to estimate cell type-specific gene expression conservation. High AUROC values indicate a high degree of cell type replicability, whereas low values suggest limited conservation. We followed the original protocol and used default settings for parameter selection, including the 50 nearest neighbors for network construction and 1,000 most variable genes for gene space reduction.

### Single nuclei WGCNA

We used the R packages hdWGCNA and WGCNA to perform WGCNA on the fibroblast and myofibroblast. We used the hdWGCNA function to construct metacells to aggregate transcriptionally neighboring single cells into pseudo-bulk metacells. Then, the metacells were processed by regular Seurat functions including NormalizeMetacells, ScaleMetacells, RunPCAMetacells, RunHarmonyMetacells, and RunUMAPMetacells were performed. Then, the expression matrix was set for fibroblasts and myofibroblasts separately. Next, after selecting the appropriate soft power by running TestSoftPowers, the co-expression network was created. In the next step harmonized module eigengenes were calculated to summarize the gene expression profile for each gene and in each sample. Then, the module connectivity was calculated, genes assigned to each module were extracted, and the results were shown as the average gene expression in all clusters. Also, the module eigengenes for each group were calculated and shown by heatmap.

### Visium data analysis

Data was aligned using Space Ranger (v1.0.0) with the reference genome GRCh38 and the human probe dataset (Visium_Human_Transcriptome_Probe_Set_v1.0_GRCh38). To correct for technical noise and improve the quality of our spatial transcriptomics dataset, we used the SpotClean method^74^. SpotClean is a computational method that leverages the negative control spots on the spatial transcriptomics array to remove technical noise and improve the accuracy of the expression measurements. First, we preprocessed the spatial transcriptomics data by filtering out lowly expressed genes and normalizing the expression values using the SCTransform method. We then used the SpotClean package to clean the data using the negative control spots. Finally, we performed downstream analysis on the cleaned expression matrix. This step was done for all 14 samples, which we merged, using the merge function of Seurat. The data were subjected to principal component analysis (PCA) for linear dimension reduction and Harmony was used to integrate the datasets. A shared nearest neighbor network was created based on Euclidean distances between cells in a multidimensional PC space (30 PCs were used) and a fixed number of neighbors per cell. This network was used to generate a 2-dimensional Uniform Manifold Approximation and Projection (UMAP) for visualization.

To identify spot specific markers, we used Seurat’s FindAllMarkers function. This method calculates log-fold changes, percentages of expression within and outside a group, and p-values of the Wilcoxon-Rank Sum test comparing a group to all cells outside that specific group. This also includes adjustments for multiple testing. A log-fold-change threshold of 0.25 and FDR < 0.05 was considered significant. The basic functions of Seurat were used for visualization.

### Cell2location analysis

To identify cell types and their spatial locations within a tissue, we used the Cell2location tool^32^. Cell2location is a Bayesian probabilistic model that integrates transcriptomic and spatial information to estimate cell type-specific gene expression patterns and their spatial distribution. We first preprocessed the spatial transcriptomics data by filtering out lowly expressed genes and normalizing the expression values. We then mapped the spatial locations of the cells onto a two-dimensional coordinate space using the cell coordinates provided by the dataset. We subsequently trained the Cell2location model on the preprocessed data using our integrated sc/snRNAseq and snATACseq data and prior information on cell type co-localization. This prior information, obtained from imaging data, was used to constrain the spatial distribution of each cell type.

Finally, we used the trained model to estimate cell type-specific gene expression patterns and their spatial distribution in the tissue. The resulting cell type-specific expression maps and spatial distributions were visualized on a spatial map of the tissue to identify the spatial organization of different cell types.

### Deconvolution of Visium Dataset

To deconvolve the spots in our spatial transcriptomics dataset, we utilized the Cell2location method. We began by preprocessing the raw data using standard quality control and filtering steps to remove low-quality spots and genes with low expression. We then normalized the data using the total-count normalization method implemented in the Cell2location package. Next, we trained the Cell2location model using our integrated sc/snRNAseq and snATACseq atlas and our spatial transcriptomics data. The single-cell RNA sequencing data were used to define the cell types present in the tissue and to estimate their gene expression signatures, while the spatial transcriptomics data were used to estimate the spatial distribution of the cell types within the tissue. We estimated the model parameters using the variational Bayesian inference algorithm implemented in the Cell2location package, running the algorithm for 5,000 iterations with a convergence threshold of 0.01 for the change in the lower bound on the marginal likelihood of the observed data.

Once the model parameters were estimated, we used the Cell2location package to predict the expression of each gene in each cell type at each spot in the tissue. This allowed us to identify the cell types present in each spot and visualize their spatial distribution within the tissue. The Cell2location method proved highly effective in deconvolving the spots in our spatial transcriptomics dataset, providing insights into the spatial organization of cells in the tissue.

### Mapping sn/scRNAseq to spatial location

To map back the cell types identified in the dissociated data (sn/scRNAseq datasets), we used the Celltrek^33^ package. We first downsampled the integrated sc/snRNAseq and snATACseq datasets to 20,000 cells. Then, using the traint function, sn/scRNAseq datasets were co-embedded with spRNAseq datasets. Subsequently, using the random forest model, single cells were mapped to their spatial locations. This analysis was performed by merging snRNAseq and immune cell types to enrich the dataset for immune cells. Regarding colocalization, the sColoc function of CellTrek was used. To identify the different cell type modules in the spRNAseq, we performed a spatial-weighted gene co-expression analysis.

## Ligand–Receptor Interactions in Spatial Transcriptomics Data

We used the CellChat v1.4.0 package to predict cell type-specific ligand–receptor interactions (1939 interactions) between cells in the spRNAseq data. Only receptors and ligands expressed in more than 10 cells in each cluster were considered. To compute the interactions, we used the computeCommunProb function with the following parameters: distance.use = TRUE, interaction.length = 200, and scale.distance = 0.01. Probability and P-values were measured for each interaction.

In addition, we utilized the COMMOT package, which provides a spatially informed approach to account for cell type locations in our data. After preprocessing each spRNAseq data, we used the ct.tl.spatial_communication function with default parameters to calculate the communication scores between each pair of cell types based on the expression levels of ligand and receptor genes. We visualized the results using the plotComm functions to generate a network.

## Finding microenvironments in spRNAseq

To identify microenvironments on the merged spatial dataset after SC transformation (SCT) we performed an NMF reduction using the STUtility package^40^. The dataset was then clustered using the first 20 factors from the NMF reduction. To identify MEs specific markers, we used Seurat’s FindAllMarkers function. Identified MEs were then annotated based on marker genes. To find correlations between the number of spots of each MEs and clinical characteristics, we extracted the number of spots for each MEs per sample and then performed a Pearson correlation analysis with clinical findings.

### ECM production score

To calculate the extracellular matrix production (ECM) score, we calculated the proportion of the expressions of collagen, proteoglycan, and glycoprotein^36^ genes in each cell types.

## CosMx Data Analysis

CosMx data was exported and analyzed in python using the scanpy and squidpy packages. Probe counts were normalized and log transformed. Cells with fewer than 30 reads were removed. Principal components were calculated which were used to determine 15 neighbors for each cell. These neighbors were used to create a UMAP and Leiden clusters (resolution 1.0). Annotation of leiden clusters was manually performed—the iPT/PEC cluster was subclustered and annotated to resolve these two populations. Annotation locations were manually inspected after annotation based on gene expression. Neighborhoods were calculated using the Squidpy function, using a neighborhood size of 20 microns. The data was integrated with our single nuclei data using SCVI, using 4 layers and 30 latent variables.

### Bulk RNAseq Analysis

The human bulk kidney RNAseq data were downloaded from GEO (<u>GSE115098</u> and <u>GSE173343)</u>. We used previously published pipeline for bulk RNAseq analysis ^75^. Briefly, the FASTQC was used to check the QC of the sequencing results. The adapters and low-quality bases were trimmed using TrimGalore (v0.4.5). The trimmed FASTQ files were aligned to the human genome (hg19/GRCh37) using STAR (v2.7.3a)^76,77^ based on GENCODE v19 annotations^76,77^. Gene expression was quantified using RSEM by calculating uniquely mapped reads as transcripts per million (TPM). To ensure accurate microdissection, we projected the TPM matrix of the tubule and glomerular compartments onto a PCA plot and samples that did not separate clearly were removed.

### Hierarchical clustering of microdissected human kidney tubule samples based on FME-gene signature

Hierarchical clustering was performed on the scaled TPM matrix of microdissected human tubules datasets using the FME-GS list as the base. Ward’s method with Euclidean distances was used to cluster the datasets. The optimal number of clusters was determined by average silhouette method.

### Kaplan Meier and Cox-regression analysis

To compare the probability of reaching the outcome; end stage kidney disease (eGFR < 15 ml/min/1.73 *m^2^*) or 40% eGFR decline, Kaplan-Meier life table survival analysis was performed. The log-rank test was used to compare the survival probability in different clusters. Cox proportional hazard ratio analysis was applied to estimate the hazard ratio of outcome of the subjects in different groups.

### Linear regression analysis

To perform the linear regression analysis between fibrosis, eGFR or eGFR decline with gene expression levels in each cell type, cells were subset based on the frequency of each cell type with a maximum number of 1000 cells proportional to its original fraction. A regression model adjusted for age, sex, race, diabetes mellitus, hypertension, and QC parameters (nFeature_Count, nCount_RNA, and percent_mt) was employed. FDR < 0.05 was considered as significant threshold.

## Statistics

Data were expressed as means ± SEM. Independent sample t-test was used to compare the continuous variable in two groups and one-way ANOVA was used to compare continuous parameters between more than two groups followed by Bonferroni post hoc test for subgroup comparisons. A *p* < 0.05 was considered as a significant. LASSO regression was performed with scikit-learn’s LASSO CV function, 10 n_splits with 3 repeats were used for parameter tuning.

## Data Availability

Raw data, processed data, and metadata from the snRNAseq, scRNAseq, snATACseq, and spRNAseq have been deposited in Gene Expression Omnibus (GEO) with the accession code of GSE211785 (**reviewer token: srahoicgfzqbfkj**) The human bulk kidney RNAseq data are available under following accession numbers: <u>GSE115098</u> and <u>GSE173343</u>. The single cell and nuclear expression and open chromatin and spatial data is also available at www.susztaklab.com (https://susztaklab.com/hk_genemap/).

## Code Availability

All the codes used for the analysis were deposited on GitHub (https://github.com/amin69upenn/Human_Kidney_Multiomics_and_Spatial_Atlas_and https://github.com/jlevins2010/FME_atlas)

## Acknowledgement

Work in the Susztak lab is supported by the NIH P50DK114786, DK076077, DK087635, DK132630 and DK105821. The study is supported by GSK, Regeneron, Boehringer Ingelheim, and Novo Nordisk. The funders have no influence on the reported results. The authors thank the Molecular Pathology and Imaging Core (P30-DK050306) and Diabetes Research Center (P30-DK19525) at University of Pennsylvania for their services.

## Author Contributions

AA, JL, ZM, JF, RS, PD, GP, DT, and TB performed experiments. AA, JL, KK, MSB, HL, SV, MSB, HY, and KC performed computational analysis. KD, BD, LM, EH, LSB, CAH, AK, PG, CMB, GP, KHK and ML offered experimental and analytical suggestions. KS was responsible for overall design and oversight of the experiments. MP performed pathological scorings. KS supervised the experiment. AA and KS wrote the original draft. All authors contributed to and approved the final version of the manuscript.

## Supplementary material

### Supplementary Tables

**Supplementary Table 1. Demographic and clinical characteristics of the subjects in the study.**

**Supplementary Table 2. QC metrics of the snRNAseq, scRNAseq, snATACseq, and spRNAseq data.**

**Supplementary Table 3. Cell type marker genes in the snRNAseq data.**

**Supplementary Table 4. Cell type marker genes in the scRNAseq data.**

**Supplementary Table 5. Cell type specific accessible regions and genes in the snATACseq data.**

**Supplementary Table 6. Cell type specific markers genes in the integrated snRNAseq, snATACseq, and scRNAseq.**

**Supplementary Table 7. Cell type specific transcription factor motif enrichment.**

**Supplementary Table 8. Module specific genes and scores in fibroblast and myofibroblast (WGCNA)**

**Supplementary Table 9. Differentially expressed genes between diseased and control samples in each cell type in the integrated snRNAseq, snATACseq, and scRNAseq.**

**Supplementary Table 10. CellChat interaction scores.**

**Supplementary Table 11. Differentially expressed genes along injured PT trajectory in the integrated snRNAseq, snATACseq, and scRNAseq.**

**Supplementary Table 12. Fibrotic microenvironment specific genes.**

**Supplementary Table 13. Clinical characteristics of the 298 and 245 tubule bulk RNAseq cohorts. Supplementary Table 14. Lasso coefficients for each gene of the FME gene signature against eGFR.**

### Supplementary Figures

Supplementary Fig. 1. Quality metrics of the sn/scRNAseq, snATACseq and spatial transcriptomics data.

(a) QC parameters of the scRNAseq. The plots depict the distribution of nFeature_Count (number of genes per cell), nCount_RNA (number of reads per cell), and mitochondrial percentage (fraction of mitochondrial genes per cell) in each sample. X axis indicates the individual sample and y-axis shows the values.

(b) QC parameters of the snRNAseq. The plots depict the distribution of nFeature_Count (number of genes per cell), nCount_RNA (umber of reads per cell), and mitochondrial percentage (fraction of mitochondrial genes per cell) in each sample. X axis indicates the individual sample and y-axis shows the values.

(c) QC parameters of the snATACseq. The plots depict the distribution of pct_reads in peaks, nFeature_ATAC per sample. X-axis indicates the individual sample and y-axis shows the values.

Supplementary Fig. 2. Multimodal single cell atlas

(a) UMAP of snRNAseq, scRNAseq, and snATACseq datasets before integration.

(b) UMAP of integrated snRNAseq, scRNAseq, and snATACseq datasets of 338,565 cells and nuclei using the SCVI tool.

(c) Annotations of cell types on integrated UMAP.

(d) The dot plots of marker genes used to annotation of 44 main cell types in the integrated dataset. The size of the dot indicates the percent positive cells, and the darkness of the color indicates average expression.

Endo_G; endothelial cells of glomerular capillary tuft, Endo_peritubular; endothelial cells of peritubular vessels, Endo_lymphatic; endothelial cells of lymphatic vessels, Mes; meseangial cells, GS_Stromal; glomerulosclerosis-specific stromal cells, VSMC/Pericyte; vascular smooth muscle cells/pericyte, PEC; parietal epithelial cells, Podo; podocyte, PT_S1; proximal tubule segment 1, PT_S2; proximal tubule segment 2, PT_S3; proximal tubule segment 3, Injured_PT; injured proximal tubule cells, DLOH; thin descending loop of Henle, C_TAL; cortical thick ascending loop of Henle, M_TAL; medullary thick ascending loop of Henle, DCT; distal convoluted tubule, CNT; connecting tubule cells, PC; principal cells of collecting duct, IC_A; Type alpha intercalated cells, IC_B; Type beta intercalated cells, NK; natural killer cells, CD4T; T lymphocytes CD4+, CD8T; T lymphocytes CD8+, dnT; double negative T cells, Prolif_Lym; proliferative lymphocyte, Th17; T helper 17 lymphocyte, B_Naive; Naive B lymphocyte, B_memory; memory B lymphocyte, RBC; red blood cells, Baso/Mast; basophil or mast cells, pDC; plasmacytoid dendritic cells, cDC; classical dendritic cells, Mac; macrophage, CD14_Mono; monocyte CD14+, CD16_Mono; monocyte CD16+.

Supplementary Fig. 3. High resolution clustering of the integrated data.

(a) UMAP of integrated snRNAseq, scRNAseq, and snATACseq datasets of 338,565 cells and nuclei and identification of 114 clusters. Right panel shows the original source data.

(b) The dot plots of marker genes used for the annotation of 144 fine clusters in the integrated dataset. The size of the dot indicates the percent positive cells, and the darkness of the color indicates average expression.

(c) The dot plot of erythropoietin (EPO) expression in the integrated snRNAseq, scRNAseq, and snATACseq datasets. The size of the dot indicates the percent positive cells, and the darkness of the color indicates average expression. Stromal cells are the main source of EPO in the kidney.

Endo_G; endothelial cells of glomerular capillary tuft, Endo_peritubular; endothelial cells of peritubular vessels, Endo_lymphatic; endothelial cells of lymphatic vessels, Mes; mesangial cells, GS_Stromal; glomerulosclerosis-specific stromal cells, VSMC/Myofib; vascular smooth muscle cells/myofibroblast, PEC; parietal epithelial cells, Podo; podocyte, PT_S1; proximal tubule segment 1, PT_S2; proximal tubule segment 2, PT_S3; proximal tubule segment 3, Injured_PT; injured proximal tubule cells, DLOH; thin descending loop of Henle, C_TAL; cortical thick ascending loop of Henle, M_TAL; medullary thick ascending loop of Henle, DCT; distal convoluted tubule, CNT; connecting tubule cells, PC; principal cells of collecting duct, IC_A; Type alpha intercalated cells, IC_B; Type beta intercalated cells, NK; natural killer cells, CD4T; T lymphocytes CD4+, CD8T; T lymphocytes CD8+, B_Naive; Naive B lymphocyte, B_memory; memory B lymphocyte, RBC; red blood cells, Baso/Mast; basophil or mast cells, pDC; plasmacytoid dendritic cells, cDC; classical dendritic cells, Mac; macrophage, CD14_Mono; monocyte CD14+, CD16_Mono; monocyte CD16+.

Supplementary Fig. 4. Integrations of snRNAseq, scRNAseq, and snATACseq datasets from multiple sources

(a) UMAP of integrated snRNAseq, scRNAseq, and snATACseq datasets (n=588,425 cells and nuclei) from Susztaklab and KPMP using the SCVI tool. Left panel shows the new annotation after integration the dataset and right panel indicates the original annotation used by KPMP.

(b) Bar charts showing cell abundance in each cell clusters (method, lab, present study annotation and KPMP annotation).

Endo_G; endothelial cells of glomerular capillary tuft, Endo_peritubular; endothelial cells of peritubular vessels, Endo_lymphatic; endothelial cells of lymphatic vessels, Mes; mesangial cells, GS_Stromal; glomerulosclerosis-specific stromal cells, VSMC/Myofib; vascular smooth muscle cells/myofibroblast, PEC; parietal epithelial cells, Podo; podocyte, PT_S1; proximal tubule segment 1, PT_S2; proximal tubule segment 2, PT_S3; proximal tubule segment 3, Injured_PT; injured proximal tubule cells, DLOH; thin descending loop of Henle, C_TAL; cortical thick ascending loop of Henle, M_TAL; medullary thick ascending loop of Henle, DCT; distal convoluted tubule, CNT; connecting tubule cells, PC; principal cells of collecting duct, IC_A; Type alpha intercalated cells, IC_B; Type beta intercalated cells, NK; natural killer cells, CD4T; T lymphocytes CD4+, CD8T; T lymphocytes CD8+, B_Naive; Naive B lymphocyte, B_memory; memory B lymphocyte, RBC; red blood cells, Baso/Mast; basophil or mast cells, pDC; plasmacytoid dendritic cells, cDC; classical dendritic cells, Mac; macrophage, CD14_Mono; monocyte CD14+, CD16_Mono; monocyte CD16+.

Supplementary Fig. 5. Integrations of snRNAseq, scRNAseq, and snATACseq datasets from multiple sources

(a) Dot plots of marker genes used for the annotation of 39 main cell types in the integrated dataset. The size of the dot indicates the percent positive cells, and the darkness of the color indicates average expression.

(b) Cell type consistency of the different datasets using MetaNeighbor. The heatmap displays the Area Under the Receiver Operating Characteristic (AUROC) scores among different cell subtypes in two different data sources (this study vs KPMP). Dendrograms were created via hierarchical clustering, based on Euclidean distances, and used average linkage for this purpose. Each row and each column are a cell type in one of the source data.

Endo_G; endothelial cells of glomerular capillary tuft, Endo_peritubular; endothelial cells of peritubular vessels, Endo_lymphatic; endothelial cells of lymphatic vessels, Mes; mesangial cells, GS_Stromal; glomerulosclerosis-specific stromal cells, VSMC/Pericyte; vascular smooth muscle cells/pericyte, PEC; parietal epithelial cells, Podo; podocyte, PT_S1; proximal tubule segment 1, PT_S2; proximal tubule segment 2, PT_S3; proximal tubule segment 3, Injured_PT; injured proximal tubule cells, DLOH; thin descending loop of Henle, C_TAL; cortical thick ascending loop of Henle, M_TAL; medullary thick ascending loop of Henle, DCT; distal convoluted tubule, CNT; connecting tubule cells, PC; principal cells of collecting duct, IC_A; Type alpha intercalated cells, IC_B; Type beta intercalated cells, NK; natural killer cells, CD4T; T lymphocytes CD4+, CD8T; T lymphocytes CD8+, dnT; double negative T cells, Prolif_Lym; proliferative lymphocyte, Th17; T helper 17 lymphocyte, B_Naive; Naive B lymphocyte, B_memory; memory B lymphocyte, RBC; red blood cells, Baso/Mast; basophil or mast cells, pDC; plasmacytoid dendritic cells, cDC; classical dendritic cells, Mac; macrophage, CD14_Mono; monocyte CD14+, CD16_Mono; monocyte CD16+.

Supplementary Fig. 6. Human kidney single nuclear open chromatin (snATACseq) data.

(a) UMAP of 58,155 nuclei after QC filtering in snATACseq dataset. 26 clusters were identified.

(b) Heatmap of top 10 differentially accessible regions per cells in the snATACseq dataset. Rows indicate the top 10 differentially accessible regions and their degree of accessibility pre cluster and columns show the cell types.

(c) Accessibility of marker genes across annotations. Reads at each locus are shown as a with a higher peak indicating more open chromatin.

(d) Enriched motif and their activity per cluster in snATACseq dataset. The table shows the enriched motif in each cell type.

(e) The dot plot indicates the motifs activity in main clusters of snATACseq dataset. The size of the dot indicates the percent positive cells, and the darkness of the color indicates average activity.

Supplementary Fig. 7. Histological images of the Visium data.

H&E imaging of the 3 control and 11 diseased samples. Each sample is H&E stained and the number of Visium spots with significant RNA quantity is indicated below. Scale bar is 1 mm.

Supplementary Fig. 8. Quality Control metrics of spatial transcriptomic data.

(a) QC parameters of the Visium spatial RNA assay. We show number of features and number of RNA counts for each individual sample.

(b) QC parameters for the CosMx analysis represented as histograms, which are parsed by sample. False codes indicates reads that do not correspond to a known gene barcode, and negative probe counts corresponds to reads to a probe that is designed not to anneal to a human RNA. Counts were aggregated by field of view (FOV) to inspect if a particular region of tissue performed poorly. These metrics are then shown in space on the slide. Lastly, we also show the location of each sample on the CosMx slide (below).

Supplementary Fig. 9. Human kidney Visium spatial transcriptomics data

(a) UMAP of the 37,436 spots, across all 14 samples. Spots were annotated based on markers shown in B, though multiple cell types likely exist within each spot.

(b) The dot plots of marker genes used for spots annotation in the spRNAseq. The size of the dot indicates the percent positive cells, and the darkness of the color indicates average expression.

(c) The heatmap of the top 10 differentially expressed genes in the identified spots in spRNAseq. Rows indicate the top 10 differentially expressed genes and their degree of accessibility pre cluster and columns show the cell types.

Endo_G; endothelial cells of glomerular capillary tuft, Endo_peritubular; endothelial cells of peritubular vessels, Endo_lymphatic; endothelial cells of lymphatic vessels, Mes; mesangial cells, GS_Stromal; glomerulosclerosis-specific stromal cells, Fib; Fibroblast, MyoFib; Myofibroblast, VSMC/Pericyte; vascular smooth muscle cells/Pericyte, PEC; parietal epithelial cells, Podo; podocyte, PT_S1; proximal tubule segment 1, PT_S2; proximal tubule segment 2, PT_S3; proximal tubule segment 3, Injured_PT; injured proximal tubule cells, DLOH; thin descending loop of Henle, C_TAL; cortical thick ascending loop of Henle, M_TAL; medullary thick ascending loop of Henle, DCT; distal convoluted tubule, CNT; connecting tubule cells, PC; principal cells of collecting duct, IC_A; Type alpha intercalated cells, IC_B; Type beta intercalated cells, NK; natural killer cells, CD4T; T lymphocytes CD4+, CD8T; T lymphocytes CD8+, B_Naive; Naive B lymphocyte, B_memory; memory B lymphocyte, RBC; red blood cells, Baso/Mast; basophil or mast cells, pDC; plasmacytoid dendritic cells, cDC; classical dendritic cells, Mac; macrophage, CD14_Mono; monocyte CD14+, CD16_Mono; monocyte CD16+.

Supplementary Fig. 10. Protein expression of human kidney cell type markers in the Human Protein Atlas Images were taken https://www.proteinatlas.org/.

Supplementary Fig. 11. Glomerular, proximal and distal tubule cells in each kidney sample (Cell2Location was used to map cells).

Supplementary Fig. 12. Immune and stromal cells in each kidney sample (Cell2Location was used to map cells).

Supplementary Fig. 13. Proximal tubule, glomerular, stromal and LOH cells in each kidney sample (CellTrek was used to map cells).

Supplementary Fig. 14. Distal tubule and immune cells in each kidney sample (CellTrek was used to map cells).

Supplementary Fig. 15. Tissue staining of the CosMx slides.

(a) Immunostained CosMx slide. CosMx slides were stained with DAPI, Pan-Cytokeratin, CD-45, and membrane stain.

(b) After imaging the slide, the flowcell (sample) was removed and the slides were then H&E stained and re-imaged.

Supplementary Fig. 16. CosMx Cell populations.

(a) Original UMAP of all cells that passed QC along with their annotations. The “PT?” population appeared to be similar to PTs but lacked classic markers.

(b) Cell Populations with annotatable markers of the CosMx data on UMAP.

(c) UMAP of all CosMx cells by sample.

(d) CosMx cell annotations across the UMAP with a single population being shown for a given UMAP demonstrating that these clusters are indeed relatively localized within the UMAP.

(e) Dot plot showing markers for each annotated CosMx population

(f) Annotation Frequency for each cell type.

Supplementary Fig. 17. CosMx SCVI integration with snRNAseq data.

After integrating with the snRNAseq data, we compared annotations from our CosMx analysis and the separately annotated snRNAseq.

(a) UMAP of integrated data demonstrating technology type of each cell within the UMAP.

(b) Comparison of CosMx annotations and snRNAseq annotations, demonstrating concordance of location within the integrated UMAP.

Supplementary Fig. 18. Location of CosMx annotated cell types within the slide. Location of annotated cell types within the two tissue sections.

Location of glomerular cell subtypes

Location of injured PT, fibroblasts and immune cells

Supplementary Fig. 19. Microanatomy of the CosMx slide

Location of glomerular cell types within a subsection of tissue, and in a single field of view (right)

Location of injured thick ascending limb, healthy injured thick ascending limb, principal cells and immune cell types within a subsection of tissue, and in a single field of view (right).

Location of distal nephron cell subtypes within a subsection of tissue and in a single field of view (right) Single field of view showing many cell types.

Supplementary Fig. 20. Neighborhood characteristics of CosMx slide

Relative type cell frequency between each sample. Orange indicates HK3039 (healthy) and blue indicates HK2844 (diseased), and frequency of neighbor annotations for each cell type for a 20-micron neighborhood (right).

Neighborhood enrichment by permuting annotations for the 20-micron neighborhood size. Lighter color indicates higher enrichment and co-localization of a given population.

Dot plots for iPT, PEC, Podocytes and PT cells expression of iPT and PEC markers across genomics modalities and protein staining of VCAM1 in PECs from the human protein atlas: https://www.proteinatlas.org/.

Supplementary Fig. 21. Sub-clustering of stromal cells in the integrated dataset

(a) UMAP of 41,422 stromal cells in the integrated dataset.

(b) The dot plots of marker genes used for cell type annotation. The size of the dot indicates the percent positive cells, and the darkness of the color indicates average expression.

(c) The single cell Weighted correlation network analysis (WGCNA) on fibroblast and myofibroblast identified specific modules in control and disease samples. The heatmaps shows the identified modules in each cell types and their scores. The lower panel shows the representative genes with high scores in each cell type.

Mes; Mesangial cells, GS_Stromal; glomerulosclerosis associated stromal cells, JGC; juxta-glomerular cells, VSMC; vascular smooth muscle cell, Fib; fibroblast, MyoFib; myofibroblast.

Supplementary Fig. 22. The integration of stromal cells with prior studies

(a) UMAP of integration of 86,481 stromal cells of current study prior papers by KPMP and Kramann et al. The color indicates the data source.

(b) Heatmap display of MetaNeighbor analysis of the similarities of the identified stromal cell types. The heatmap displays the Area Under the Receiver Operating Characteristic (AUROC) scores among different cell subtypes in three different sources of datasets, which were determined using a set of highly variable genes (HVG). Dendrograms were created via hierarchical clustering, based on Euclidean distances, and used average linkage for this purpose. Each row and column is a cell type in the source data.

Supplementary Fig. 23. Expression of stromal markers in the Human Protein Atlas kidney samples. Images were taken https://www.proteinatlas.org/.

Supplementary Fig. 24. Stromal sub-clustering in human snATACseq data.

(a) UMAP of 5,822 stromal cells in human snATACseq dataset.

(b) The dot plot of gene markers used to annotate the different stromal cell types in snATACseq dataset. The dot plot shows the gene activity in different cell types. The size of the dot indicates the percent positive cells, and the darkness of the color indicates average expression.

(c) The dot plot of motif activity of enriched motifs in each identified stromal cluster. The size of the dot indicates the percent positive cells, and the darkness of the color indicates average expression.

Mes; Mesangial cells, GS_Stromal; glomerulosclerosis associated stromal cells, JGC; juxta-glomerular cells, VSMC; vascular smooth muscle cell, Fib; fibroblast, MyoFib; myofibroblast.

Supplementary Fig. 25. Gene ontology analysis of microenvironment gene signature

(a) Gene ontology analysis (DAVID) of the microenvironment gene signature. The X axis shows the -Log10 (FDR) of the identified pathways and Y axis lists the pathways.

(b) Person correlation of the number of spots assigned to each microenvironment and eGFR and fibrosis.

Supplementary Fig. 26. The spatial microenvironments in the human kidney. Four spatial microenvironments were identified in human kidney samples using matrix factorization: glomerular (green), tubule (brown), fibrotic (red) and immune (blue). The ECM score (calculated by aggregate expression collagen and other glycoprotein, and proteoglycan) in Visium data shown next to the tissue microenvironment information.

Supplementary Fig. 27. Different cell types in kidney microenvironment.

Left panel kidney microenvironment in tissue samples, right panel shows cell abundance mapped using Cell2location (endothelial cell, injured_PT cells, stromal cells and immune cells)

Supplementary Fig. 28. Immunostaining of the lymphocyte and plasma cells in regions of fibrosis-associated immune cells aggregates.

Immunofluorescent staining of T-cell markers (CD4) and plasma cells (IGKC) in diseased human kidney samples.

Spatial RNAseq (10x Visium with CellTrek imputation) and immunohistochemistry staining of T-cell, B-cell, and plasma cells in diseased human kidney samples in the regions of fibrosis-associated immune cells aggregates.

Supplementary Fig. 29. *In situ* mass spectrometry of the control, and diseased samples to verify cell types in fibrotic microenvironments.

The left panel shows the Visium 10x data cellular microenvironment. The right panel shows the IMC staining of the area, markers and colors are listed on the right.

Supplementary Fig. 30. Cell-cell interaction analysis in the spRNAseq dataset in fibrotic microenvironment.

(a) Weighted total interaction strength of the CXCL, SPP1, TGFb, and PDGF pathways in control and diseased samples in the fibrotic microenvironment (left panel). The spatial location of the identified cell-cell communications pathways (CXCL, SPP1, TGFb, and PDGF) in control and diseased sample in the fibrotic microenvironment were shown in the right panel. The arrows indicate the source and targets of the identified pathways.

(b) Expression of CD34 and CDH5 as the markers of high endothelial venules (HEVs) in the fibrotic microenvironments.

(c) Volcano plot of differentially expressed genes from CosMx data. Cells with an immune neighbor within 20 microns were compared against cells without an immune neighbor for both the PT and iPT population. Logfold change > 0 indicates increased expression in cell with an immune neighbor, while logfoldchange < 0 indicates increased expression in cells without an immune neighbor. Log10(pvalue) is indicated by the y-axis. Genes with an adjusted p-value < 0.01 are marked in orange.

(d) The dot plot of expression of ligands and receptors in regions of FME in integrated sn/scRNAseq and snATACseq data. The size of the dot indicates the percent positive cells and the darkness of the color indicates average expression (right panel). The gray indicates control and red indicates diseased group.

Endo; endothelial cells, MyoFib; myofibroblast, Fib; fibroblast, VSMC/Pericyte; vascular smooth muscle cells and pericyte, Podo; podocyte, Mes; mesangial cells, PT; proximal tubule cells, iPT; injured proximal tubule cells, LOH; loop of Henle, DCT; distal convoluted tubule, CNT; connecting tubule, PC; principal cells of the collecting duct.

Supplementary Fig. 31. Spatial characteristics of injured PT subclusters on CosMx.

(a) The CosMx iPT population for each sample was subclustered using a leiden algorithm with resolution of 0.3.

(b) The top 10 differentially expressed genes for each subcluster.

(c) iPT subcluster localization within the entire slide. Views of specific regions indicated by inset boxes are shown in Supplementary Fig. 31d.

(d) iPT subclusters visualized on H&E. Subset images showing populations on H&E. Blue cells correspond with cluster 0 (iPT_APOE), orange with cluster 1 (iPT_SPP1) and green with cluster 2 (iPT_KRT7).

(e) Frequency of immune and fibroblast neighbors for each iPT subtype within each sample within 20 microns is shown below. We performed testing using a Wilcoxon rank sum test between each population within a sample. These subtypes had significantly different immune neighbors and fibroblast neighbors with each sample. HK3039 fibroblasts: iPT_APOE vs iPT_SPP1, p-value = 7e-39. HK3039 immune cells: iPT_APOE vs iPT_SPP1, p-value = 4e-67. HK2844 fibroblasts iPT_APOE vs iPT_SPP1, p-value = 9e-9, iPT_KRT7 vs iPT_SPP1, p-value = 9e-46. HK2844 immune cells iPT_APOE vs iPT_SPP1, p-value = 3e-16, iPT_KRT7 vs iPT_SPP1, p-value = 9e-11.

(f) iPT subcluster neighborhood enrichment within a 20-micron neighborhood size. Lighter color indicates higher enrichment and co-localization of a given population.

Supplementary Fig. 32. Comparison of control and diseased samples in the integrated sn/scRNAseq and snATACseq.

(a) UMAP distribution of control and diseased samples in the integrated snRNA/scRNAseq dataset.

(b) Heatmaps of cell type distribution in the control and diseased conditions.

(c) Heat of gene expression changes correlating with eGFR and fibrosis in each cell type. Red indicates larger number of correlating cells.

Endo; endothelial cells Mes; mesangial cells, Stroma; stromal cells, PEC; parietal epithelial cells, Podo; podocyte, PT; proximal tubule cells, Injured_PT; injured proximal tubule cells, LOH; loop of Henle, DCT; distal convoluted tubule, CNT; connecting tubule cells, PC; principal cells of collecting duct, IC_A; Type alpha intercalated cells, IC_B; Type beta intercalated cells.

Supplementary Fig. 33. Cell type correlations and distances of Cell Trek data.

(a) Identification of neighborhoods based on spatial location of the CellTrek imputations.

(b) Spatial cell proximity index of CellTrek cell types

(c) Neighborhoods parsed by Control and Disease sample types

(d) Cell type distances between different cell types. Glomerular, proximal tubular and distal tubular neighborhoods demonstrated by red boxes.

Supplementary Fig. 34. Injured PT cells in the control and diseased human kidney samples.

(a) Gene ontology enrichment for top pathways in iPT-VCAM1+ and iPT-HAVCR1+ cells.

(b) The spatial location of iPT_VCAM1+ (blue) and iPT_HAVCR1+ (red) cells in a diseased kidney sample based on CellTrek imputation.

(c) Expression of HAVCR1 and VCAM1 (color indicates level of expression).

(d) The gene co-expression network indicates one type of injured_PT in control human kidneys, upper panel shows the heatmap of co-expressed genes.

(e) Expression of *CC1* in imputed iPT from CellTrek, and raw *VCAM1*, and *HAVCR1* expression in a healthy human sample.

Supplementary Fig. 35. Injured PT cells in mouse diabetic kidney disease samples.

(a) UMAP of (n=28,579) of proximal convoluted tubule and two types of injured PT cell nuclei (left panel).

(b) The dot plots of marker genes used for the annotation of PCT, iPT_VCAM1+, and iPT_HAVCR1+ in the integrated dataset. The size of the dot indicates the percent positive cells, and the darkness of the color indicates average expression.

(c) Monocle cell trajectory of PT and iPT cells sub-clustering trajectory from PT to iPT-VCAM1+ and iPT-HAVCR1+ in mouse snRNAseq (left panel). Cells are colored by pseudotime (right panel).

(d) Expression of the *Vcam1* and *Havcr1* along the trajectory.

Supplementary Fig. 36. Injured PT cells in the snATACseq data.

(a) UMAP of 24,256 injured PT sub-cluster in human snATACseq dataset.

(b) Violin plot of expressions of *VCAM1*, *HAVCR1* and other genes in in two types of iPTs in snATACseq dataset.

(c) Monocle trajectory analysis of PT and injured_PT cells in the snATACseq dataset. The heatmap shows trajectory analysis based on chromatin accessibility.

(d) Feature plots of transcription factor enriched motif in iPT_HAVCR1+ cell type.

Supplementary Fig. 37. FME gene expression predicts kidney outcomes

Hierarchical clustering of 245 human kidney tubule samples based on the expression of 1,100 randomly picked genes.

Kaplan Meier analysis with Log-rank test was used to compare the survival of the 3 different clusters. Renal survival was defined as cases reaching end stage renal disease or greater than 40% eGFR decline.

Single cell expression enrichment of genes associated with eGFR decline. The heatmap shows the cell type enrichment of genes associated with eGFR decline (red indicates more genes with cell type expression). Endo; endothelial cells, Stroma; stromal cells, PEC; parietal epithelial cells, Podo; podocyte, PT; proximal tubule cells, LOH; loop of Henle, DCT; distal convoluted tubule, CNT; connecting tubule, PC; principal cells of collecting duct, IC_A; Type alpha intercalated cells. Spatial expression and microenvironment enrichment of genes associated with eGFR decline. GME: glomerular, TME: tubule, FME: fibrosis, IME: immune microenvironment.

Using the LASSO regression of all FME genes against eGFR, Kaplan Meier Analysis was re-performed using clustering of subsets of gene-- those with a Lasso coefficient that is non-zero and the genes with the most negative coefficients (Supplementary Table 14).

