## supplemental figures for "Spatially resolved human kidney multi-omics single cell atlas highlights the key role of the fibrotic microenvironment in kidney disease progression"

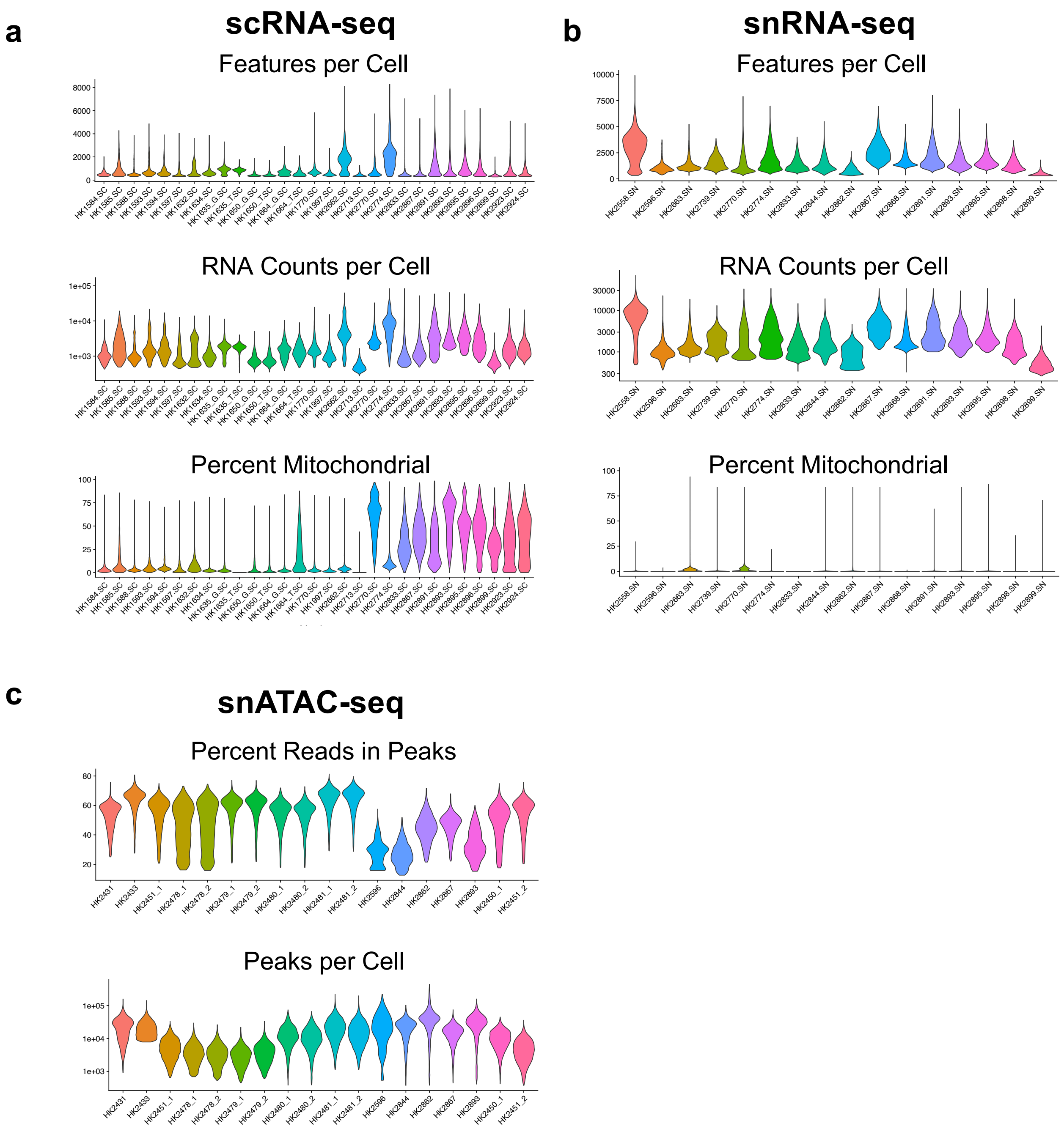


Supplementary Fig. 1

**Supplementary Fig. 1. Quality metrics of the sn/scRNAseq, snATACseq and spatial transcriptomics data.**

**(a)** QC parameters of the scRNAseq. The plots depict the distribution of nFeature_Count (number of genes per cell), nCount_RNA (number of reads per cell), and mitochondrial percentage (fraction of mitochondrial genes per cell) in each sample. X axis indicates the individual sample and y-axis shows the values.

**(b)** QC parameters of the snRNAseq. The plots depict the distribution of nFeature_Count (number of genes per cell), nCount_RNA (umber of reads per cell), and mitochondrial percentage (fraction of mitochondrial genes per cell) in each sample. X axis indicates the individual sample and y-axis shows the values.

**(c)** QC parameters of the snATACseq. The plots depict the distribution of pct_reads in peaks, nFeature_ATAC per sample. X-axis indicates the individual sample and y-axis shows the values.

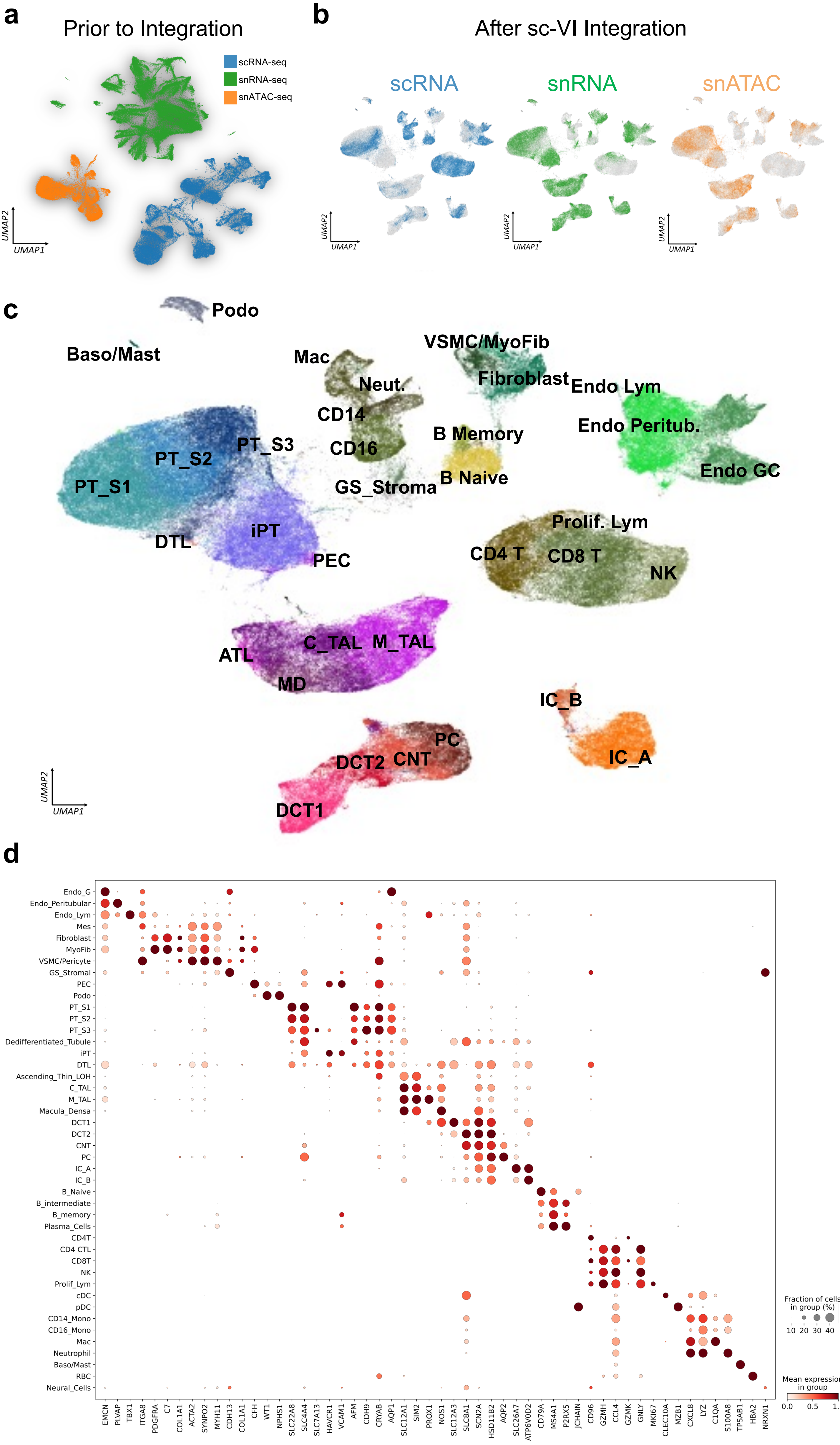


**Supplementary Fig. 2**

**Supplementary Fig. 2. Multimodal single cell atlas**
**(a)** UMAP of snRNAseq, scRNAseq, and snATACseq datasets before integration.
**(b)** UMAP of integrated snRNAseq, scRNAseq, and snATACseq datasets of 338,565 cells and nuclei using the SCVI tool.
**(c)** Annotations of cell types on integrated UMAP.
**(d)** The dot plots of marker genes used to annotation of 44 main cell types in the integrated dataset. The size of the dot indicates the percent positive cells, and the darkness of the color indicates average expression.

Endo_G; endothelial cells of glomerular capillary tuft, Endo_peritubular; endothelial cells of peritubular vessels, Endo_lymphatic; endothelial cells of lymphatic vessels, Mes; meseangial cells, GS_Stromal; glomerulosclerosis-specific stromal cells, VSMC/Pericyte; vascular smooth muscle cells/pericyte, PEC; parietal epithelial cells, Podo; podocyte, PT_S1; proximal tubule segment 1, PT_S2; proximal tubule segment 2, PT_S3; proximal tubule segment 3, Injured_PT; injured proximal tubule cells, DLOH; thin descending loop of Henle, C_TAL; cortical thick ascending loop of Henle, M_TAL; medullary thick ascending loop of Henle, DCT; distal convoluted tubule, CNT; connecting tubule cells, PC; principal cells of collecting duct, IC_A; Type alpha intercalated cells, IC_B; Type beta intercalated cells, NK; natural killer cells, CD4T; T lymphocytes CD4+, CD8T; T lymphocytes CD8+, dnT; double negative T cells, Prolif_Lym; proliferative lymphocyte, Th17; T helper 17 lymphocyte, B_Naive; Naive B lymphocyte, B_memory; memory B lymphocyte, RBC; red blood cells, Baso/Mast; basophil or mast cells, pDC; plasmacytoid dendritic cells, cDC; classical dendritic cells, Mac; macrophage, CD14_Mono; monocyte CD14+, CD16_Mono; monocyte CD16+.

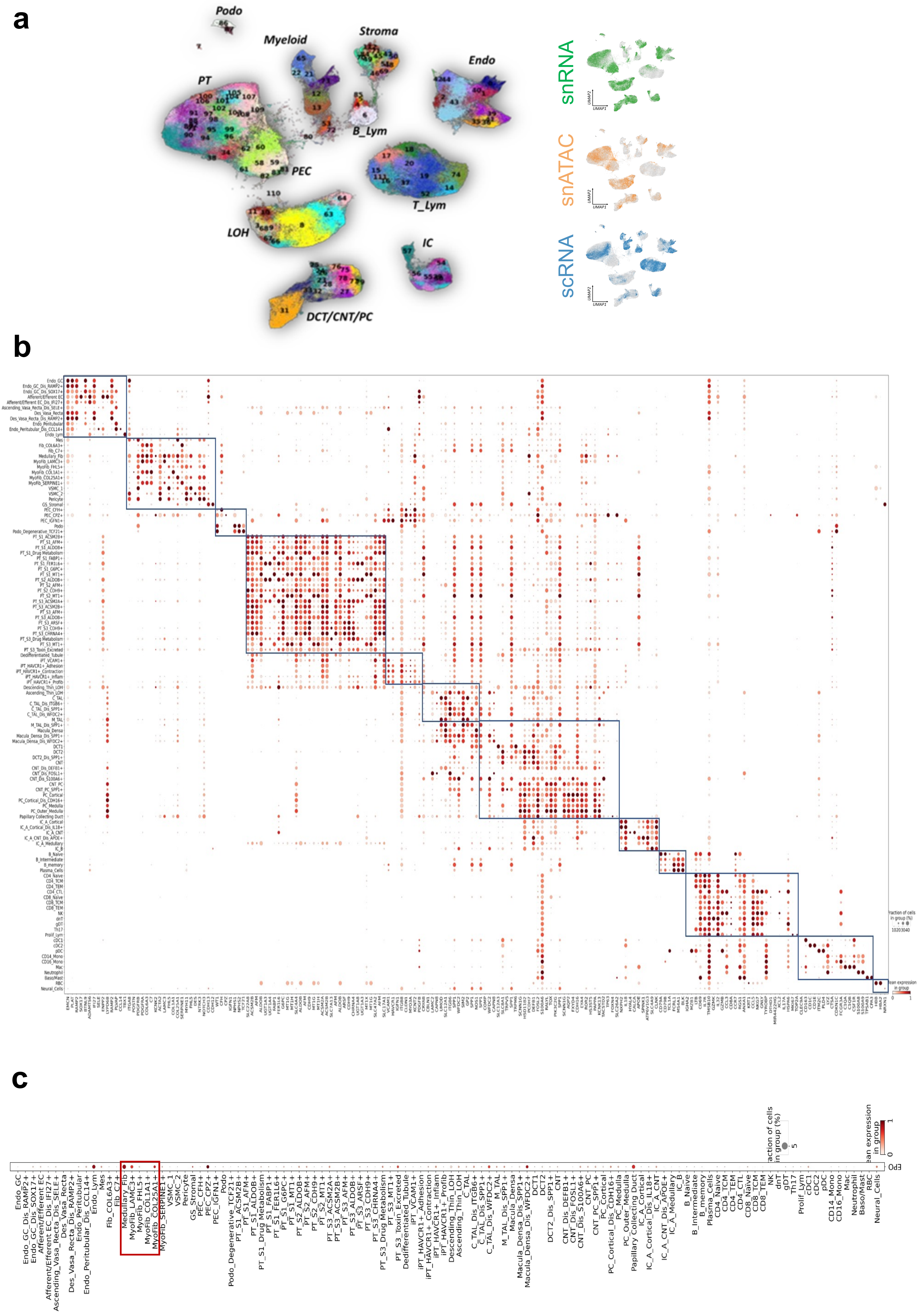


**Supplementary Fig. 3**

**Supplementary Fig. 3. High resolution clustering of the integrated data.**
**(a)** UMAP of integrated snRNAseq, scRNAseq, and snATACseq datasets of 338,565 cells and nuclei and identification of 114 clusters. Right panel shows the original source data.
**(b)** The dot plots of marker genes used for the annotation of 144 fine clusters in the integrated dataset. The size of the dot indicates the percent positive cells, and the darkness of the color indicates average expression.
**(c)** The dot plot of erythropoietin (*EPO*) expression in the integrated snRNAseq, scRNAseq, and snATACseq datasets. The size of the dot indicates the percent positive cells, and the darkness of the color indicates average expression. Stromal cells are the main source of *EPO* in the kidney.

Endo_G; endothelial cells of glomerular capillary tuft, Endo_peritubular; endothelial cells of peritubular vessels, Endo_lymphatic; endothelial cells of lymphatic vessels, Mes; mesangial cells, GS_Stromal; glomerulosclerosis-specific stromal cells, VSMC/Myofib; vascular smooth muscle cells/myofibroblast, PEC; parietal epithelial cells, Podo; podocyte,  PT_S1; proximal tubule segment 1, PT_S2; proximal tubule segment 2, PT_S3; proximal tubule segment 3, Injured_PT; injured proximal tubule cells, DLOH; thin descending loop of Henle, C_TAL; cortical thick ascending loop of Henle, M_TAL; medullary thick ascending loop of Henle, DCT; distal convoluted tubule, CNT; connecting tubule cells, PC; principal cells of collecting duct, IC_A; Type alpha intercalated cells, IC_B; Type beta intercalated cells, NK; natural killer cells, CD4T; T lymphocytes CD4+, CD8T; T lymphocytes CD8+, B_Naive; Naive B lymphocyte, B_memory; memory B lymphocyte, RBC; red blood cells, Baso/Mast; basophil or mast cells, pDC; plasmacytoid dendritic cells, cDC; classical dendritic cells, Mac; macrophage, CD14_Mono; monocyte CD14+, CD16_Mono; monocyte CD16+.

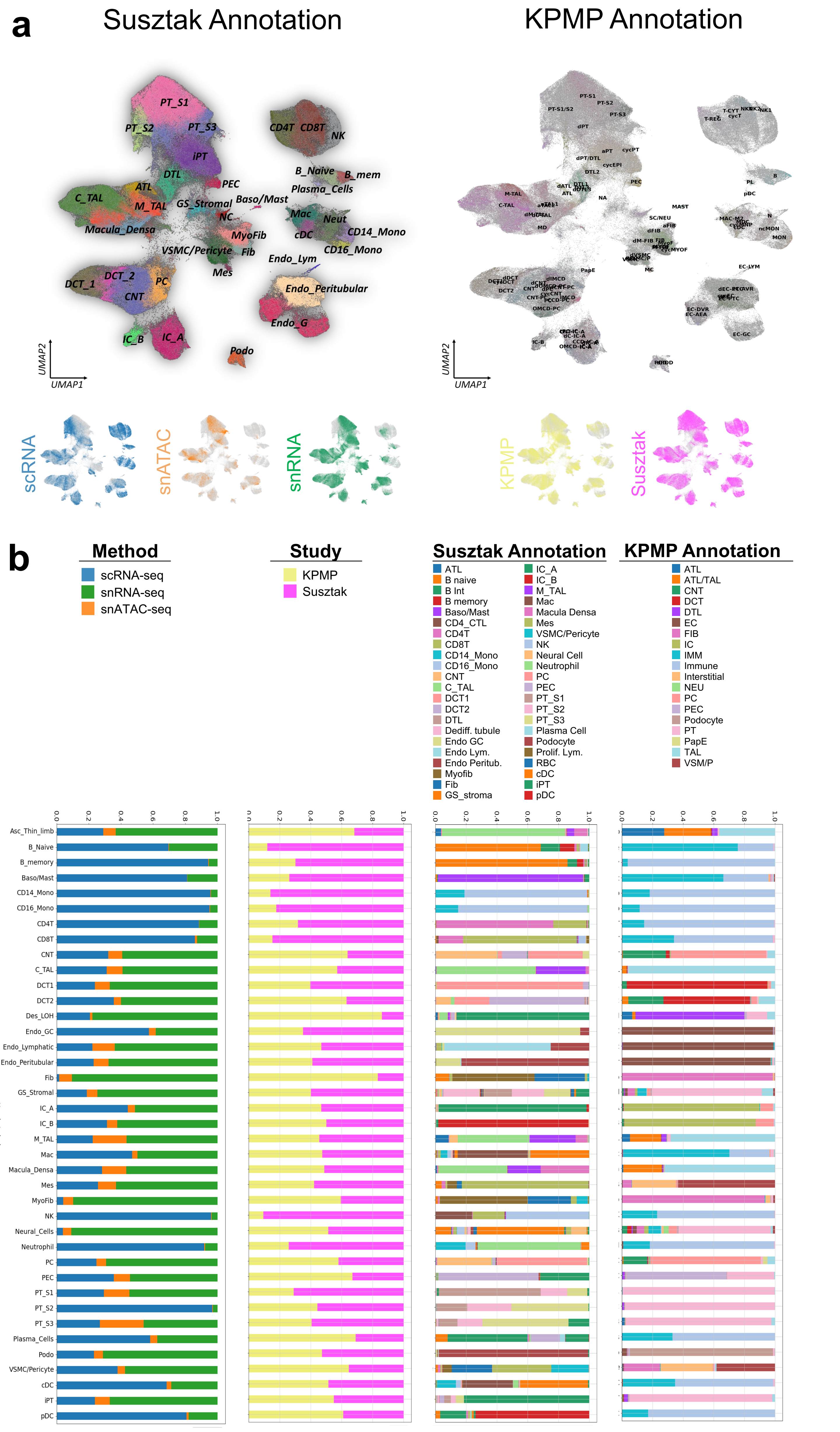


Supplementary Fig. 4

**Supplementary Fig. 4. Integrations of snRNAseq, scRNAseq, and snATACseq datasets from multiple sources**
**(a)** UMAP of integrated snRNAseq, scRNAseq, and snATACseq datasets (n=588,425 cells and nuclei) from Susztaklab and KPMP using the SCVI tool. Left panel shows the new annotation after integration the dataset and right panel indicates the original annotation used by KPMP.
**(b)** Bar charts showing cell abundance in each cell clusters (method, lab, present study annotation and KPMP annotation).

Endo_G; endothelial cells of glomerular capillary tuft, Endo_peritubular; endothelial cells of peritubular vessels, Endo_lymphatic; endothelial cells of lymphatic vessels, Mes; mesangial cells, GS_Stromal; glomerulosclerosis-specific stromal cells, VSMC/Myofib; vascular smooth muscle cells/myofibroblast, PEC; parietal epithelial cells, Podo; podocyte, PT_S1; proximal tubule segment 1, PT_S2; proximal tubule segment 2, PT_S3; proximal tubule segment 3, Injured_PT; injured proximal tubule cells, DLOH; thin descending loop of Henle, C_TAL; cortical thick ascending loop of Henle, M_TAL; medullary thick ascending loop of Henle, DCT; distal convoluted tubule, CNT; connecting tubule cells, PC; principal cells of collecting duct, IC_A; Type alpha intercalated cells, IC_B; Type beta intercalated cells, NK; natural killer cells, CD4T; T lymphocytes CD4+, CD8T; T lymphocytes CD8+, B_Naive; Naive B lymphocyte, B_memory; memory B lymphocyte, RBC; red blood cells, Baso/Mast; basophil or mast cells, pDC; plasmacytoid dendritic cells, cDC; classical dendritic cells, Mac; macrophage, CD14_Mono; monocyte CD14+, CD16_Mono; monocyte CD16+.

a

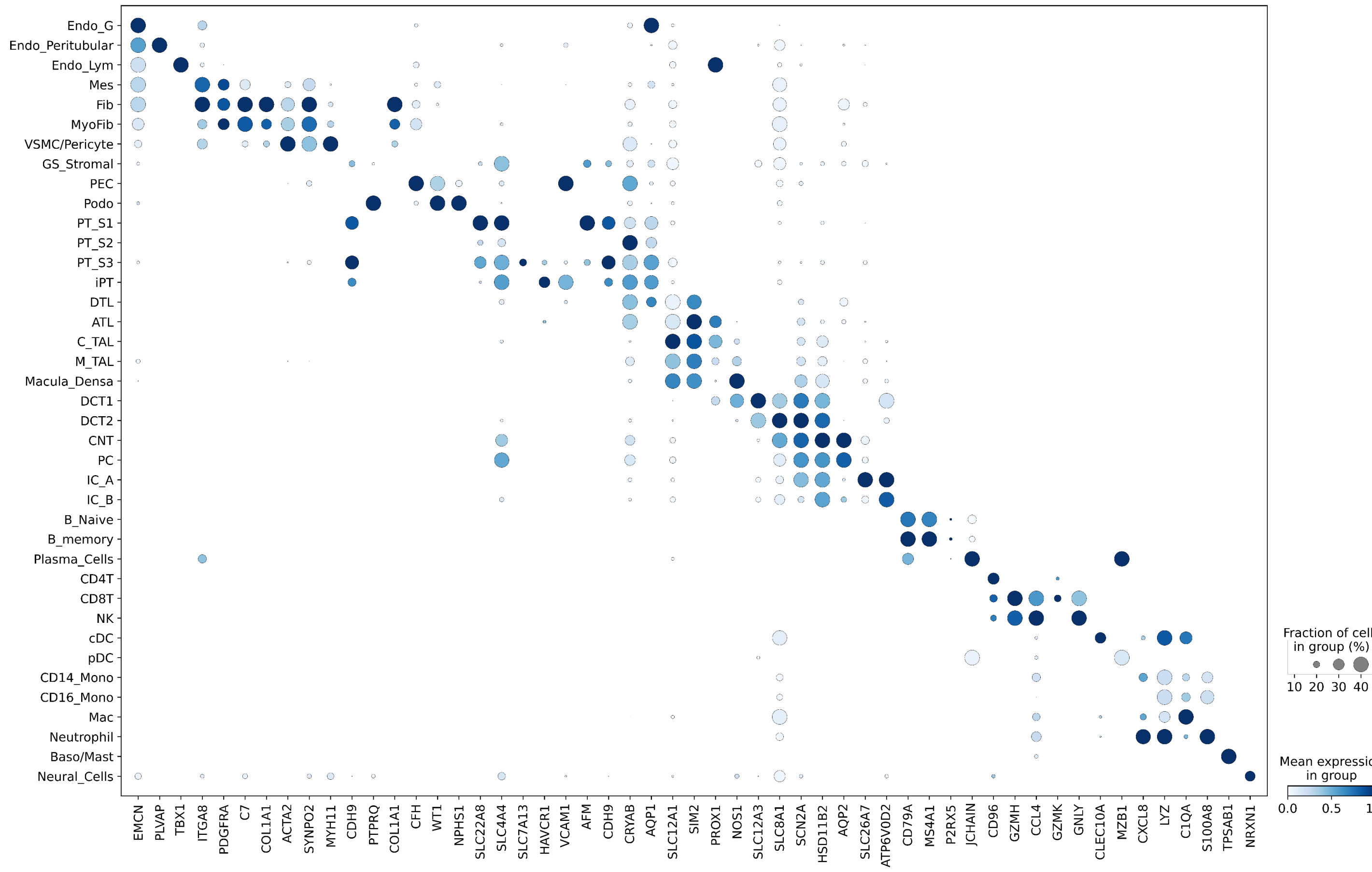


b

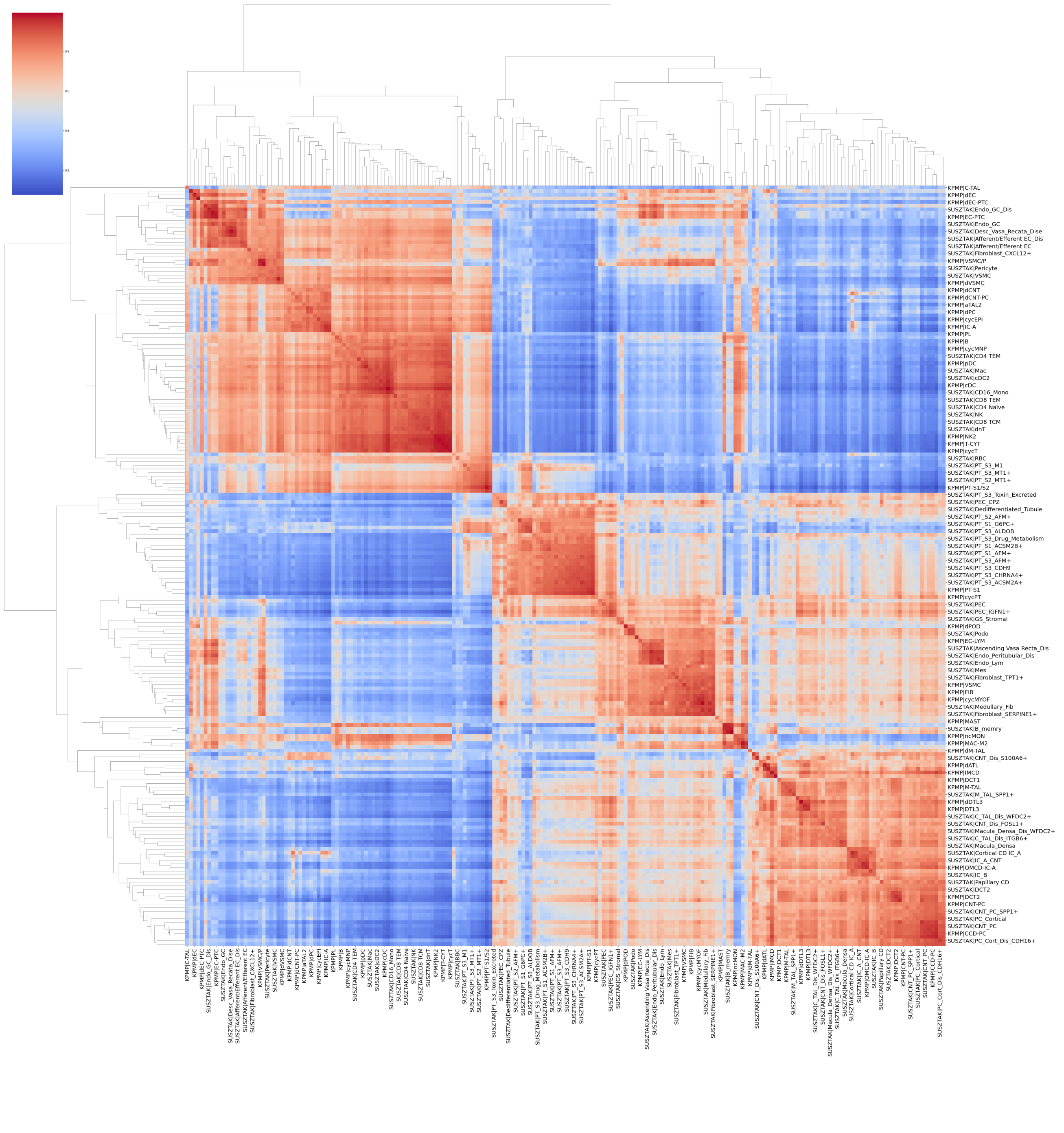

**Supplementary Fig. 5**

**Supplementary Fig. 5. Integrations of snRNAseq, scRNAseq, and snATACseq datasets from multiple sources**

**(a)** Dot plots of marker genes used for the annotation of 39 main cell types in the integrated dataset. The size of the dot indicates the percent positive cells, and the darkness of the color indicates average expression.

**(b)** Cell type consistency of the different datasets using MetaNeighbor. The heatmap displays the Area Under the Receiver Operating Characteristic (AUROC) scores among different cell subtypes in two different data sources (this study vs KPMP). Dendrograms were created via hierarchical clustering, based on Euclidean distances, and used average linkage for this purpose. Each row and each column are a cell type in one of the source data.

Endo_G; endothelial cells of glomerular capillary tuft, Endo_peritubular; endothelial cells of peritubular vessels, Endo_lymphatic; endothelial cells of lymphatic vessels, Mes; mesangial cells, GS_Stromal; glomerulosclerosis-specific stromal cells, VSMC/Pericyte; vascular smooth muscle cells/pericyte, PEC; parietal epithelial cells, Podo; podocyte, PT_S1; proximal tubule segment 1, PT_S2; proximal tubule segment 2, PT_S3; proximal tubule segment 3, Injured_PT; injured proximal tubule cells, DLOH; thin descending loop of Henle, C_TAL; cortical thick ascending loop of Henle, M_TAL; medullary thick ascending loop of Henle, DCT; distal convoluted tubule, CNT; connecting tubule cells, PC; principal cells of collecting duct, IC_A; Type alpha intercalated cells, IC_B; Type beta intercalated cells, NK; natural killer cells, CD4T; T lymphocytes CD4+, CD8T; T lymphocytes CD8+, dnT; double negative T cells, Prolif_Lym; proliferative lymphocyte, Th17; T helper 17 lymphocyte, B_Naive; Naive B lymphocyte, B_memory; memory B lymphocyte, RBC; red blood cells, Baso/Mast; basophil or mast cells, pDC; plasmacytoid dendritic cells, cDC; classical dendritic cells, Mac; macrophage, CD14_Mono; monocyte CD14+, CD16_Mono; monocyte CD16+.

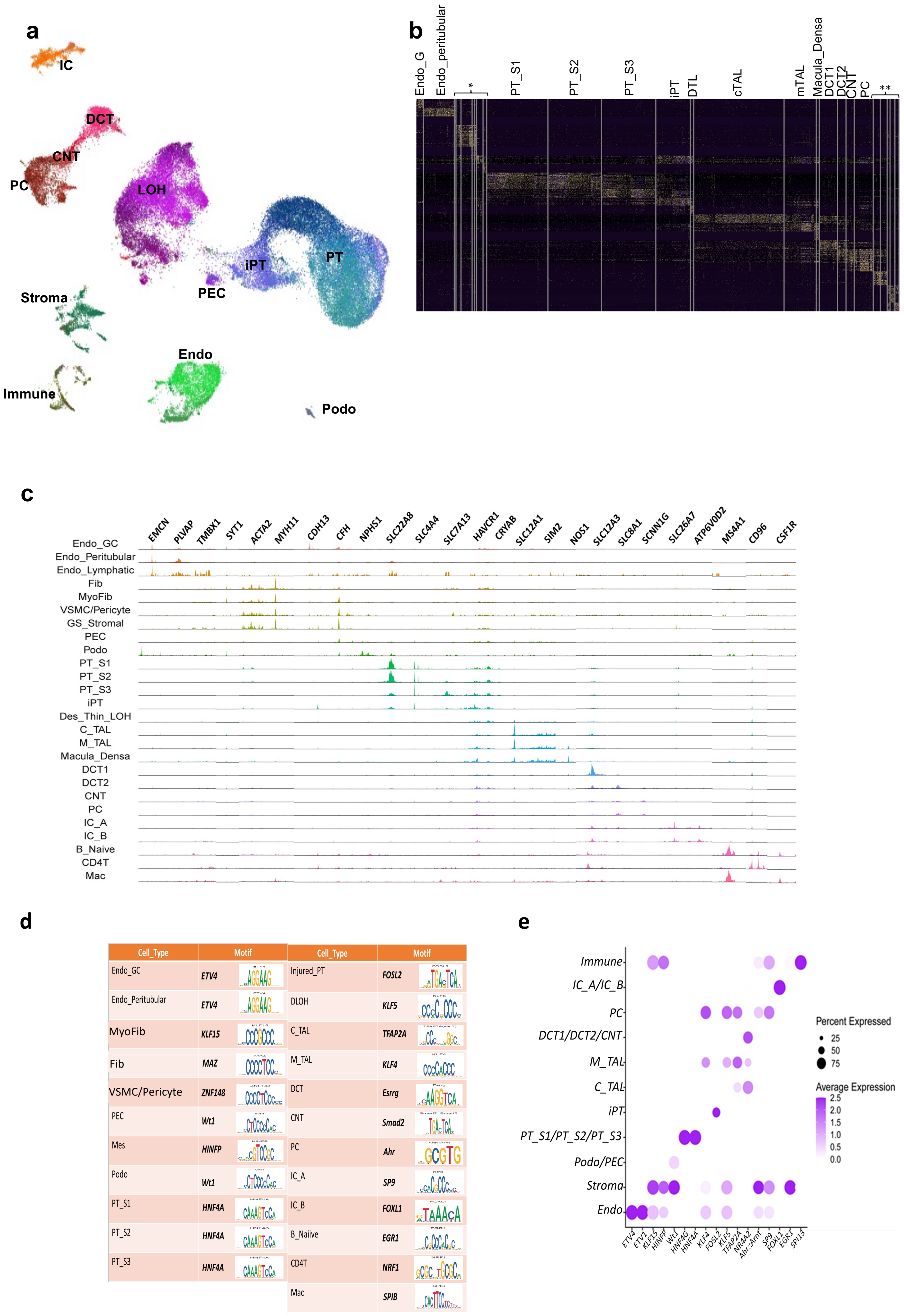


**Supplementary Fig. 6**

**Supplementary Fig. 6. Human kidney single nuclear open chromatin (snATACseq) data.**

**(a)** UMAP of 58,155 nuclei after QC filtering in snATACseq dataset. 26 clusters were identified.

**(b)** Heatmap of top 10 differentially accessible regions per cells in the snATACseq dataset. Rows indicate the top 10 differentially accessible regions and their degree of accessibility pre cluster and columns show the cell types. Populations indicated within the * region are Endo_lym, Fib, MyoFib, VSMC/Pericyte,GS_stromal, PEC, Podo (labeled left to right). Populations indicated in the ** region are IC_A, IC_B, B_naive, CD4T, and Mac, (labeled left to right).

**(c)** Accessibility of marker genes across annotations. Reads at each locus are shown as a with a higher peak indicating more open chromatin.

**(d)** Enriched motif and their activity per cluster in snATACseq dataset. The table shows the enriched motif in each cell type.

**(e)** The dot plot indicates the motifs activity in main clusters of snATACseq dataset. The size of the dot indicates the percent positive cells, and the darkness of the color indicates average activity.

Endo_GC; endothelial cells of glomerular capillary tuft, Endo_peritubular; endothelial cells of peritubular vessels, Endo_lymphatic; endothelial cells of lymphatic vessels, Mes; meseangial cells, GS_Stromal; glomerulosclerosis-specific stromal cells, VSMC/Pericyte; vascular smooth muscle cells/pericyte, PEC; parietal epithelial cells, Podo; podocyte, PT_S1; proximal tubule segment 1, PT_S2; proximal tubule segment 2, PT_S3; proximal tubule segment 3, Injured_PT; injured proximal tubule cells, DLOH; thin descending loop of Henle, C_TAL; cortical thick ascending loop of Henle, M_TAL; medullary thick ascending loop of Henle, DCT; distal convoluted tubule, CNT; connecting tubule cells, PC; principal cells of collecting duct, IC_A; Type alpha intercalated cells, IC_B; Type beta intercalated cells, NK; natural killer cells, CD4T; T lymphocytes CD4+, CD8T; T lymphocytes CD8+, dnT; double negative T cells, Prolif_Lym; proliferative lymphocyte, Th17; T helper 17 lymphocyte, B_Naive; Naive B lymphocytes, Mac; macrophage

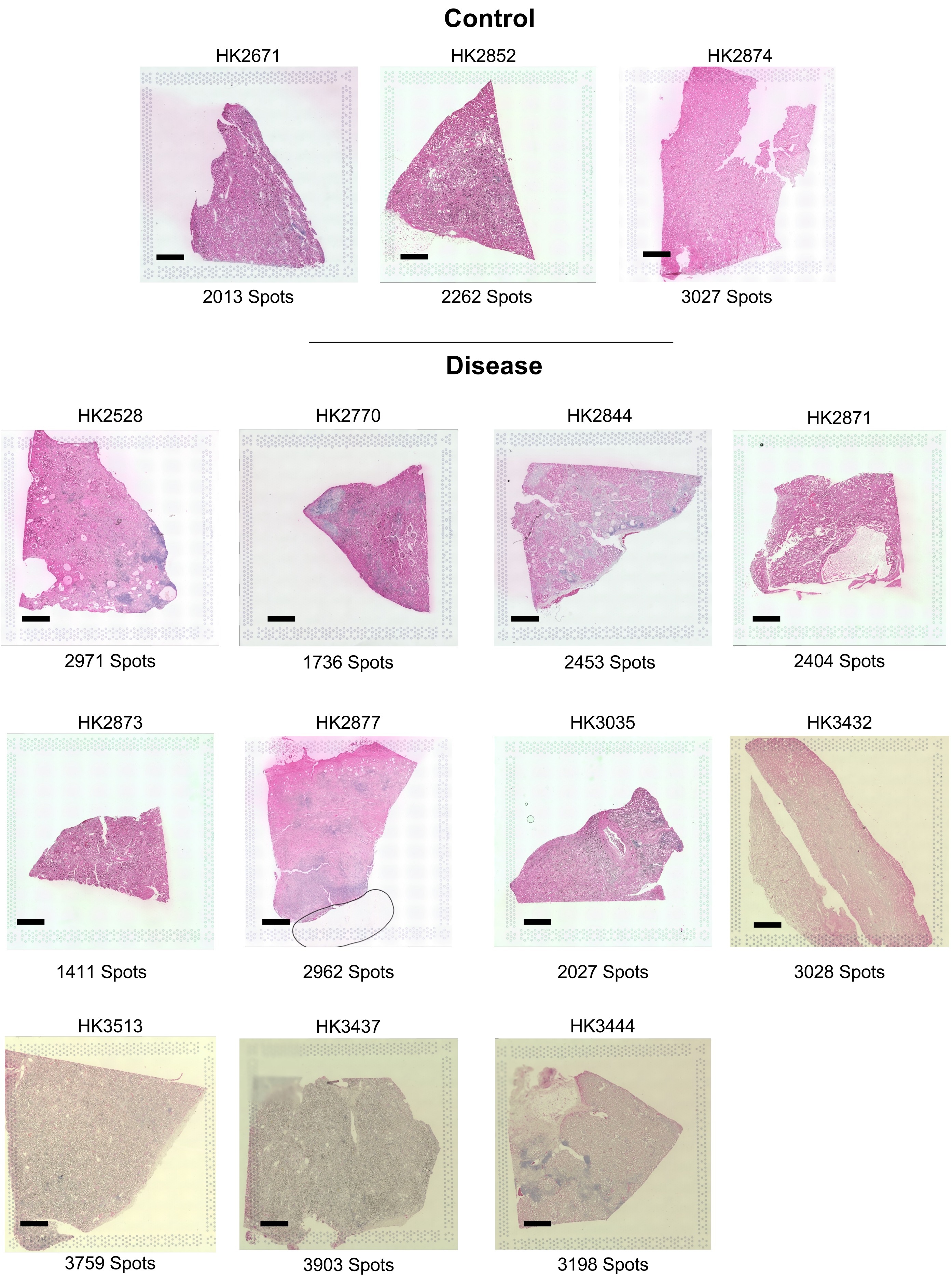


**Supplementary Fig. 7**

**Supplementary Fig. 7. Histological images of the Visium data.**

H&E imaging of the 3 control and 11 diseased samples. Each sample is H&E stained and the number of Visium spots with significant RNA quantity is indicated below. Scale bar is 1 mm.

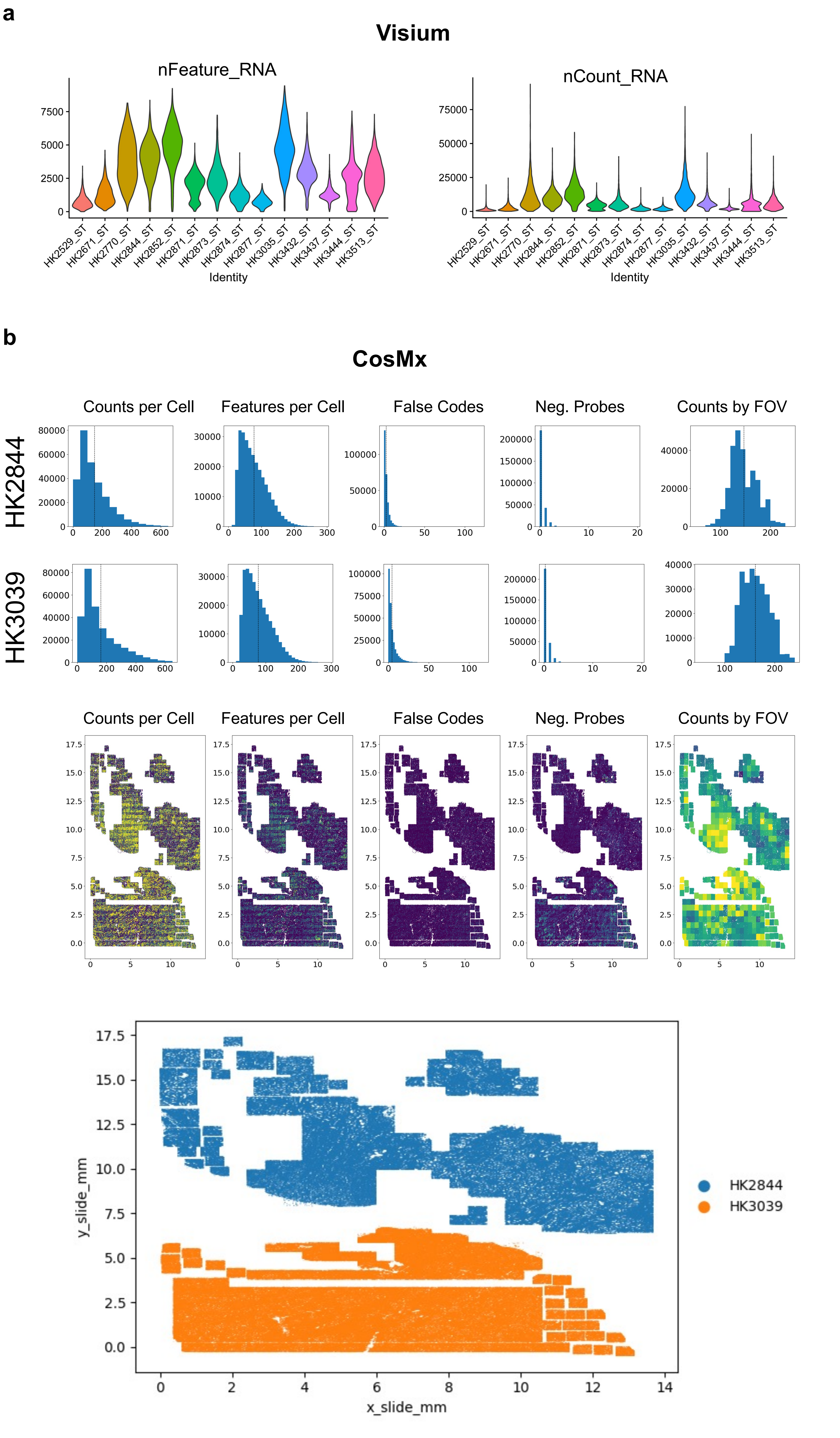


Supplementary Fig. 8

**Supplementary Fig 8. Quality Control metrics of spatial transcriptomic data.**

**(a)** QC parameters of the Visium spatial RNA assay. We show number of features and number of RNA counts for each individual sample.

**(b)** QC parameters for the CosMx analysis represented as histograms, which are parsed by sample. False codes indicates reads that do not correspond to a known gene barcode, and negative probe counts corresponds to reads to a probe that is designed not to anneal to a human RNA. Counts were aggregated by field of view (FOV) to inspect if a particular region of tissue performed poorly. These metrics are then shown in space on the slide. Lastly, we also show the location of each sample on the CosMx slide (below).

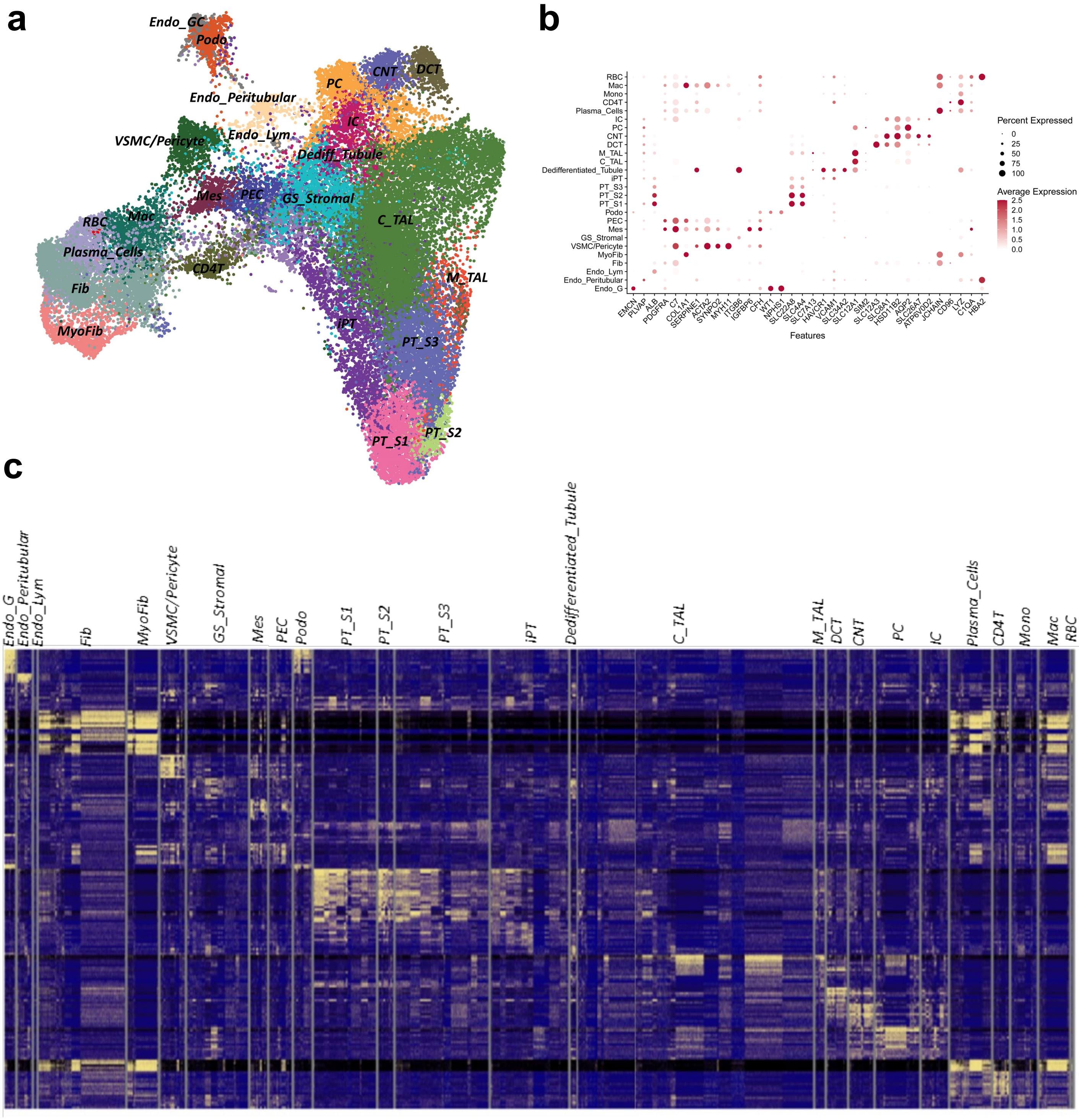


**Supplementary Fig. 9**

**Supplementary Fig. 9. Human kidney Visium spatial transcriptomics data**
**(a)** UMAP of the 37,436 spots, across all 14 samples. Spots were annotated based on markers shown in B, though multiple cell types likely exist within each spot.
**(b)** The dot plots of marker genes used for spots annotation in the spRNAseq. The size of the dot indicates the percent positive cells, and the darkness of the color indicates average expression.
**(c)** The heatmap of the top 10 differentially expressed genes in the identified spots in spRNAseq. Rows indicate the top 10 differentially expressed genes and their degree of accessibility pre cluster and columns show the cell types.

Endo_G; endothelial cells of glomerular capillary tuft, Endo_peritubular; endothelial cells of peritubular vessels, Endo_lymphatic; endothelial cells of lymphatic vessels, Mes; mesangial cells, GS_Stromal; glomerulosclerosis-specific stromal cells, Fib; Fibroblast, MyoFib; Myofibroblast, VSMC/Pericyte; vascular smooth muscle cells/Pericyte, PEC; parietal epithelial cells, Podo; podocyte, PT_S1; proximal tubule segment 1, PT_S2; proximal tubule segment 2, PT_S3; proximal tubule segment 3, Injured_PT; injured proximal tubule cells, DLOH; thin descending loop of Henle, C_TAL; cortical thick ascending loop of Henle, M_TAL; medullary thick ascending loop of Henle, DCT; distal convoluted tubule, CNT; connecting tubule cells, PC; principal cells of collecting duct, IC_A; Type alpha intercalated cells, IC_B; Type beta intercalated cells, NK; natural killer cells, CD4T; T lymphocytes CD4+, CD8T; T lymphocytes CD8+, B_Naive; Naive B lymphocyte, B_memory; memory B lymphocyte, RBC; red blood cells, Baso/Mast; basophil or mast cells, pDC; plasmacytoid dendritic cells, cDC; classical dendritic cells, Mac; macrophage, CD14_Mono; monocyte CD14+, CD16_Mono; monocyte CD16+.

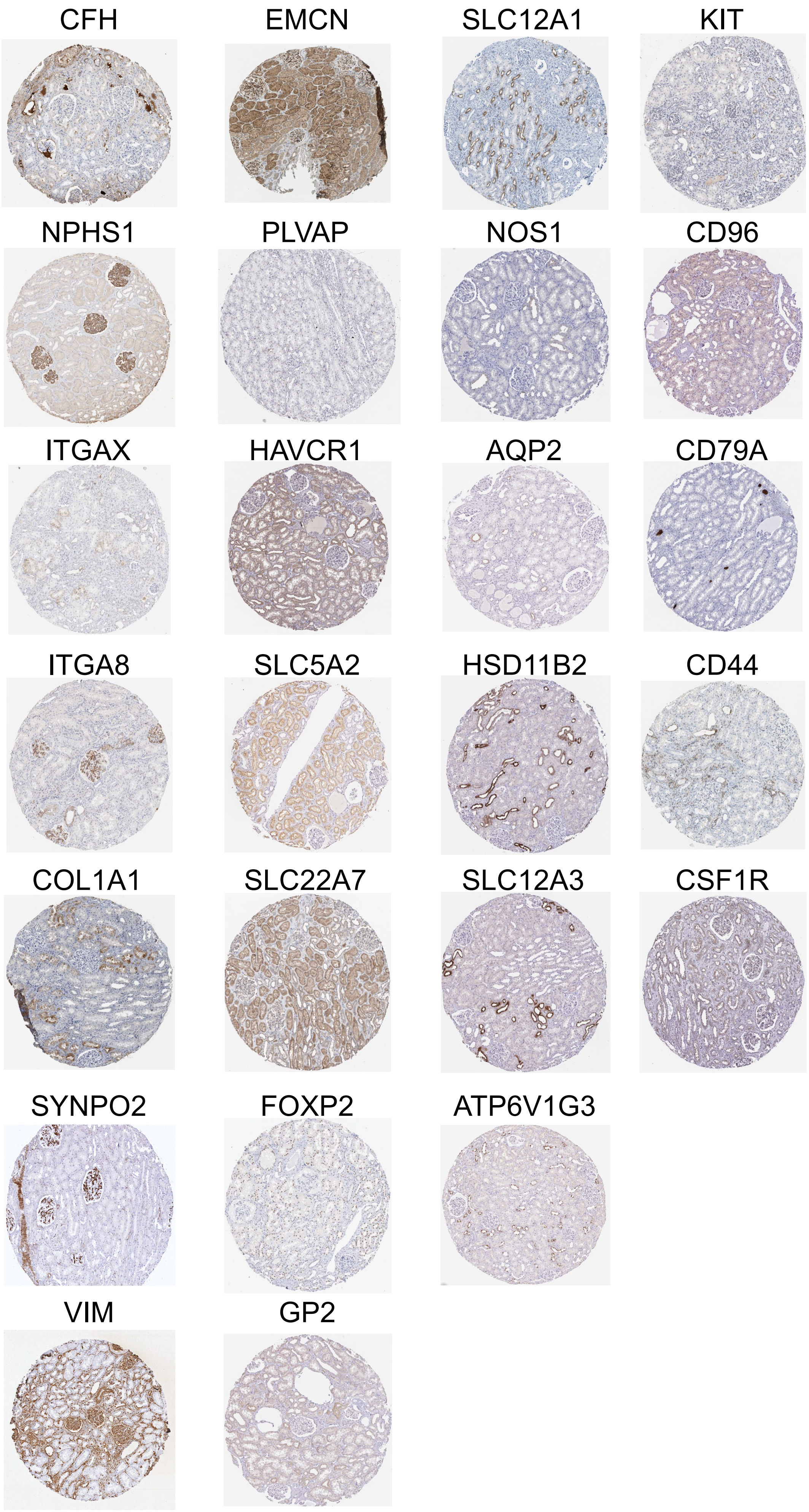


**Supplementary Fig. 10**

**Supplementary Fig. 10. Protein expression of human kidney cell type markers in the Human Protein Atlas**

Images were taken https://www.proteinatlas.org/.

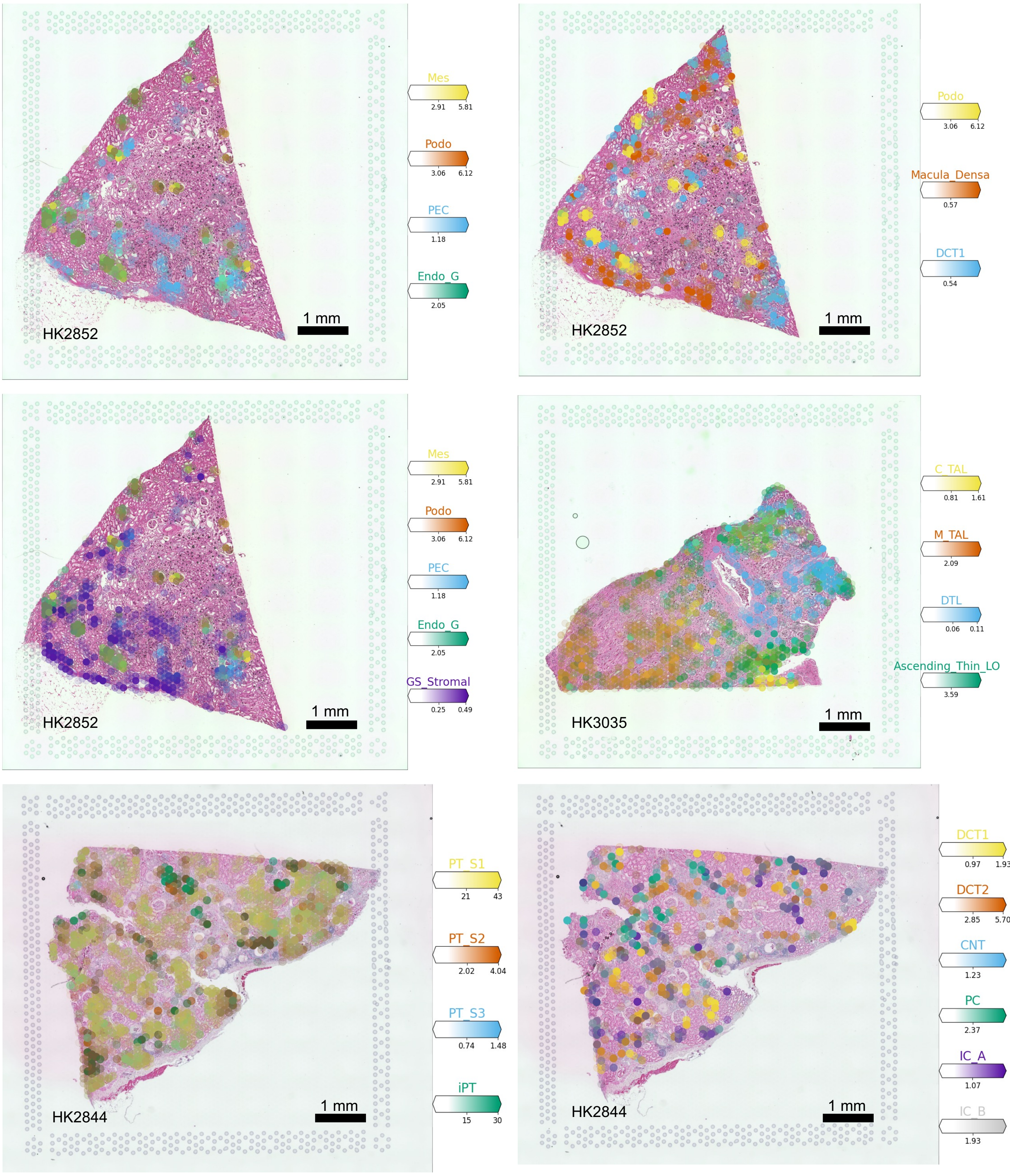


**Supplementary Fig. 11**

**Supplementary Fig. 11. Glomerular, proximal and distal tubule cells in each kidney sample**
(Cell2Location was used to map cells).

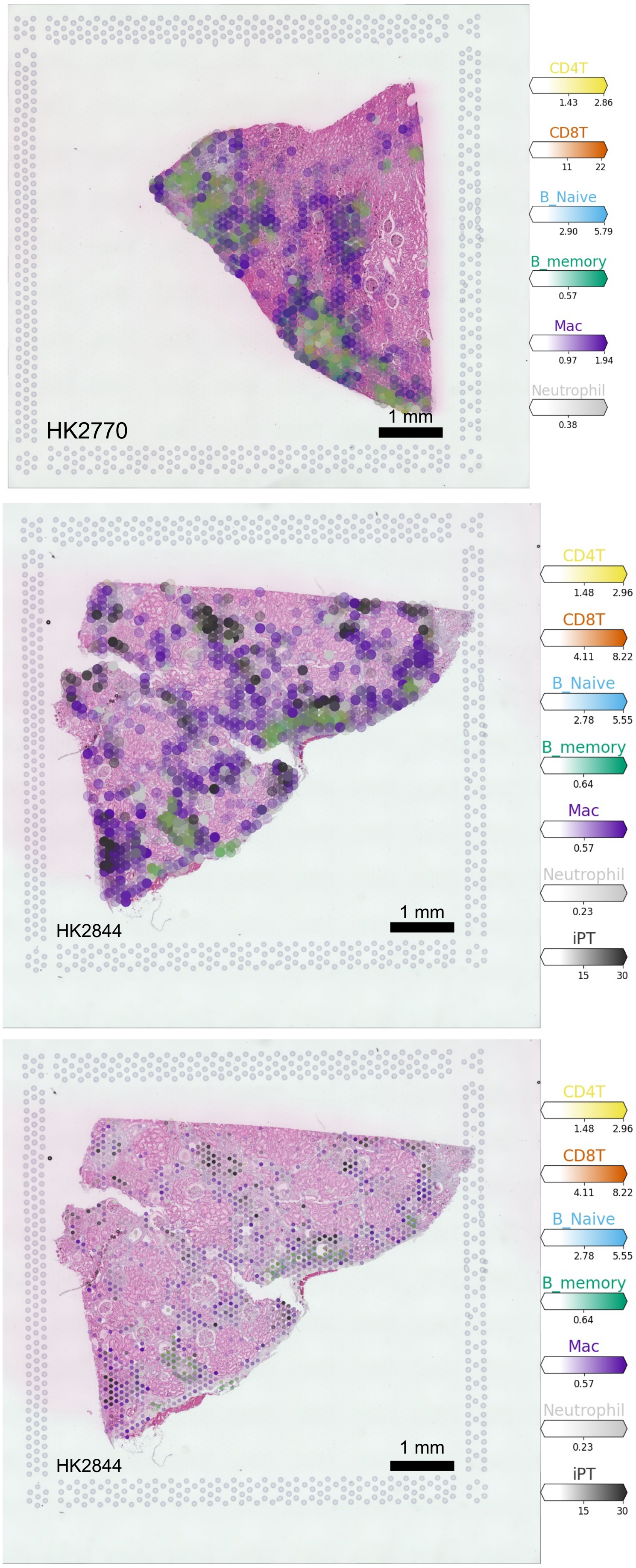


**Supplementary Fig. 12**

**Supplementary Fig. 12. Immune and stromal cells in each kidney sample** (Cell2Location was used to map cells).

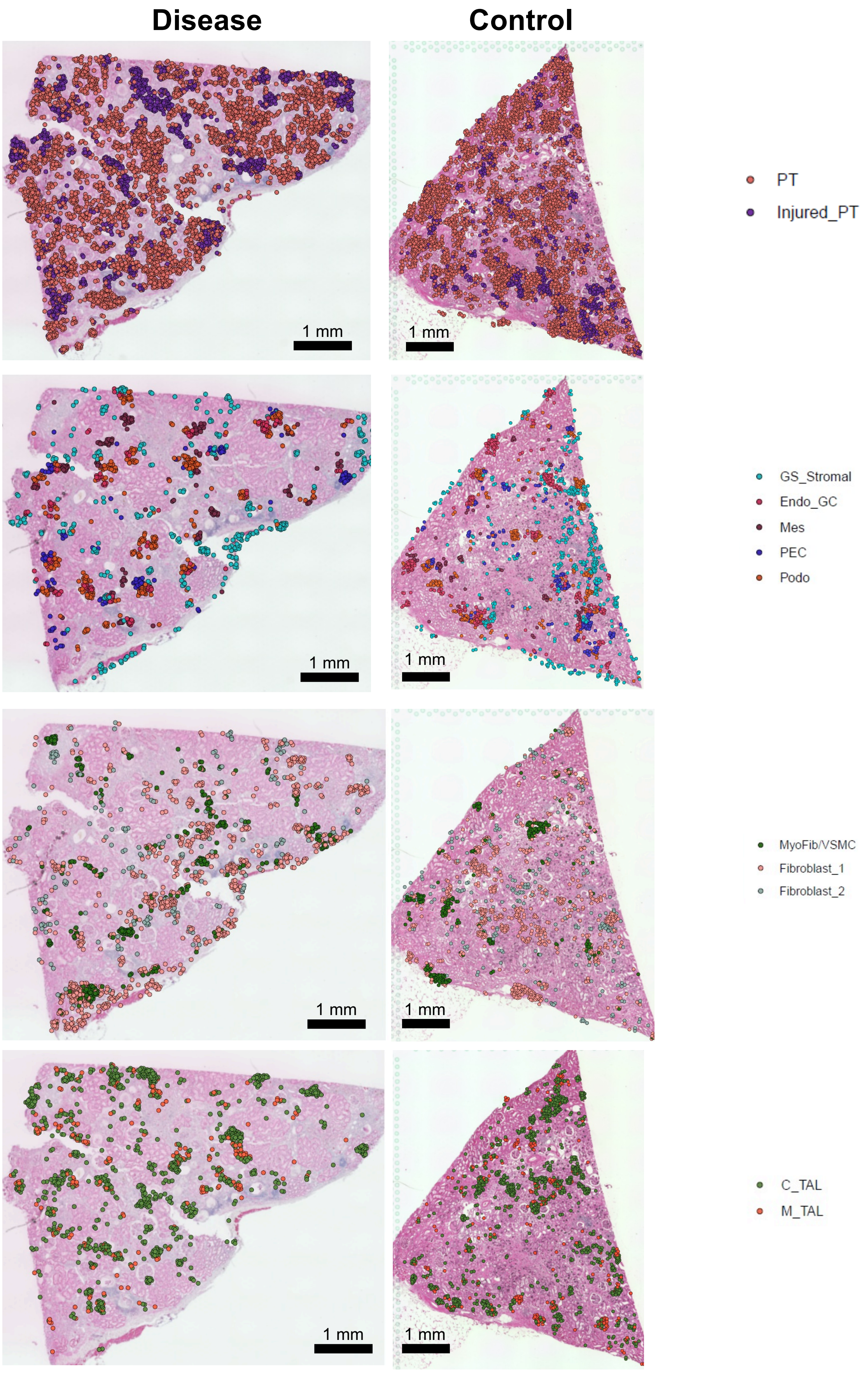


Supplementary Fig. 13

**Supplementary Fig. 13. Proximal tubule, glomerular, stromal and LOH cells in each kidney sample** (CellTrek was used to map cells).

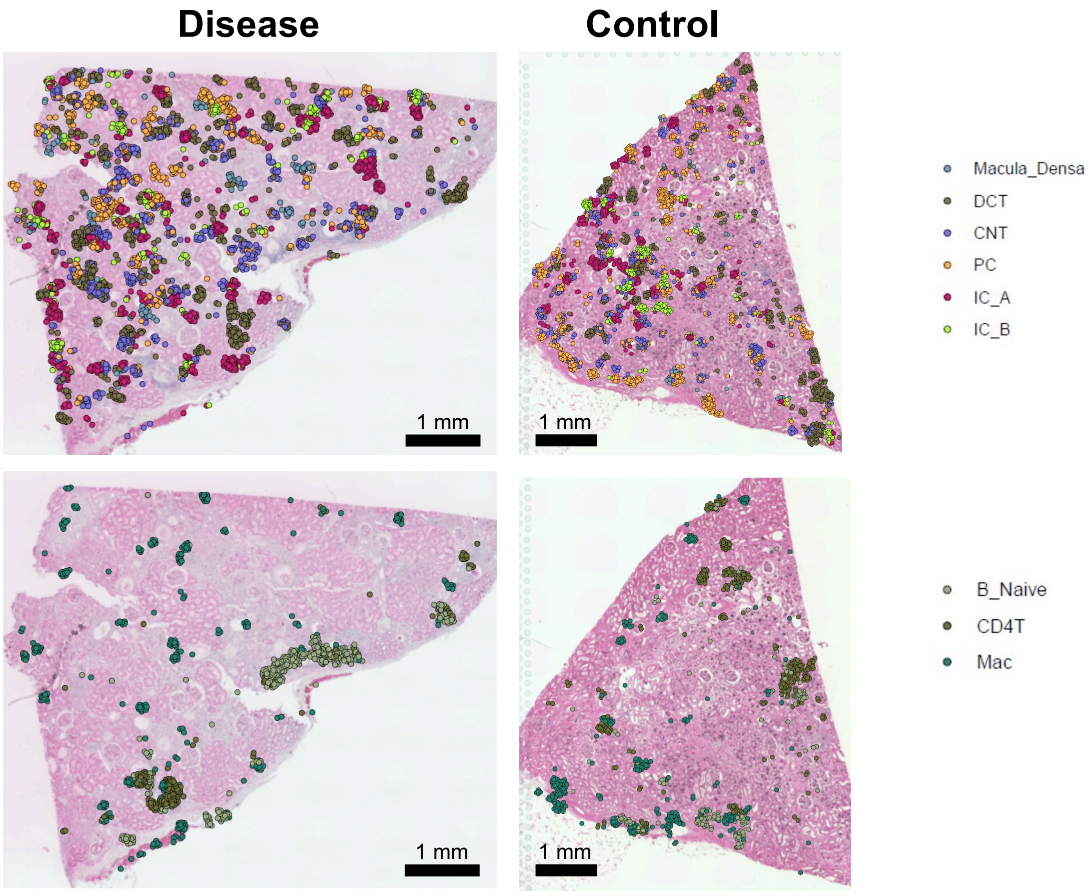


**Supplementary Fig. 14**

**Supplementary Fig. 14. Distal tubule and immune cells in each kidney sample** (CellTrek was used to map cells).

a

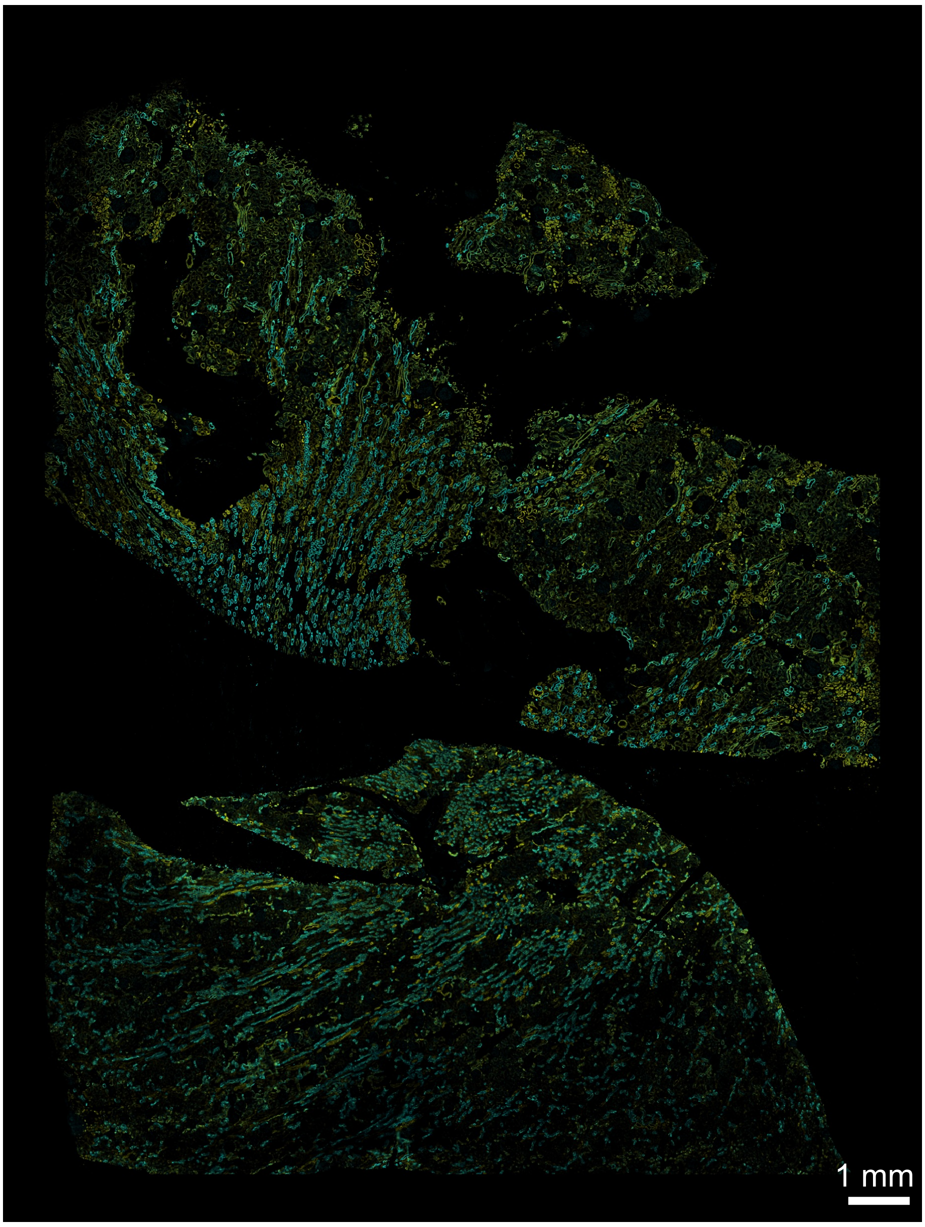


b

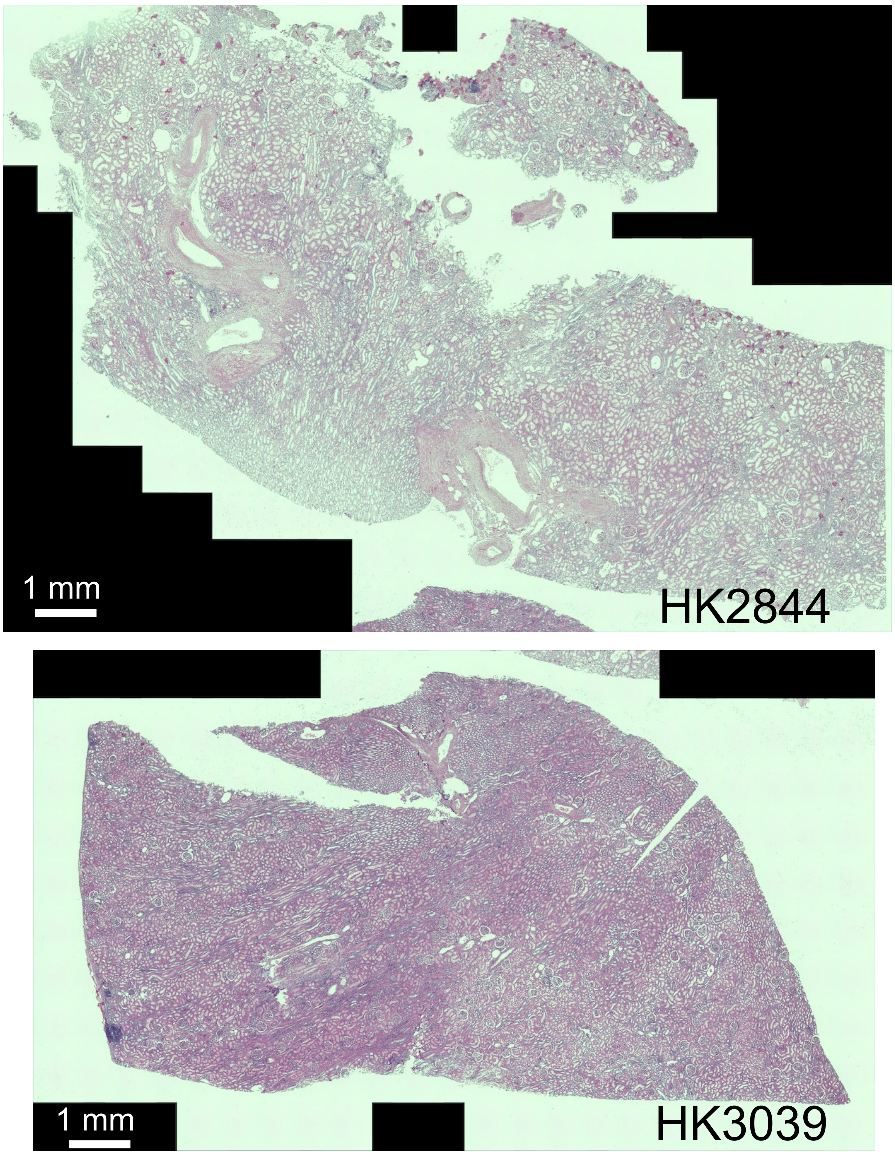


**Supplementary Fig. 15**

**Supplementary Fig 15. Tissue staining of the CosMx slides.**

**(a)** Immunostained CosMx slide. CosMx slides were stained with DAPI, Pan-Cytokeratin, CD-45, and membrane stain.

**(b)** After imaging the slide, the flowcell (sample) was removed and the slides were then H&E stained and re-imaged.

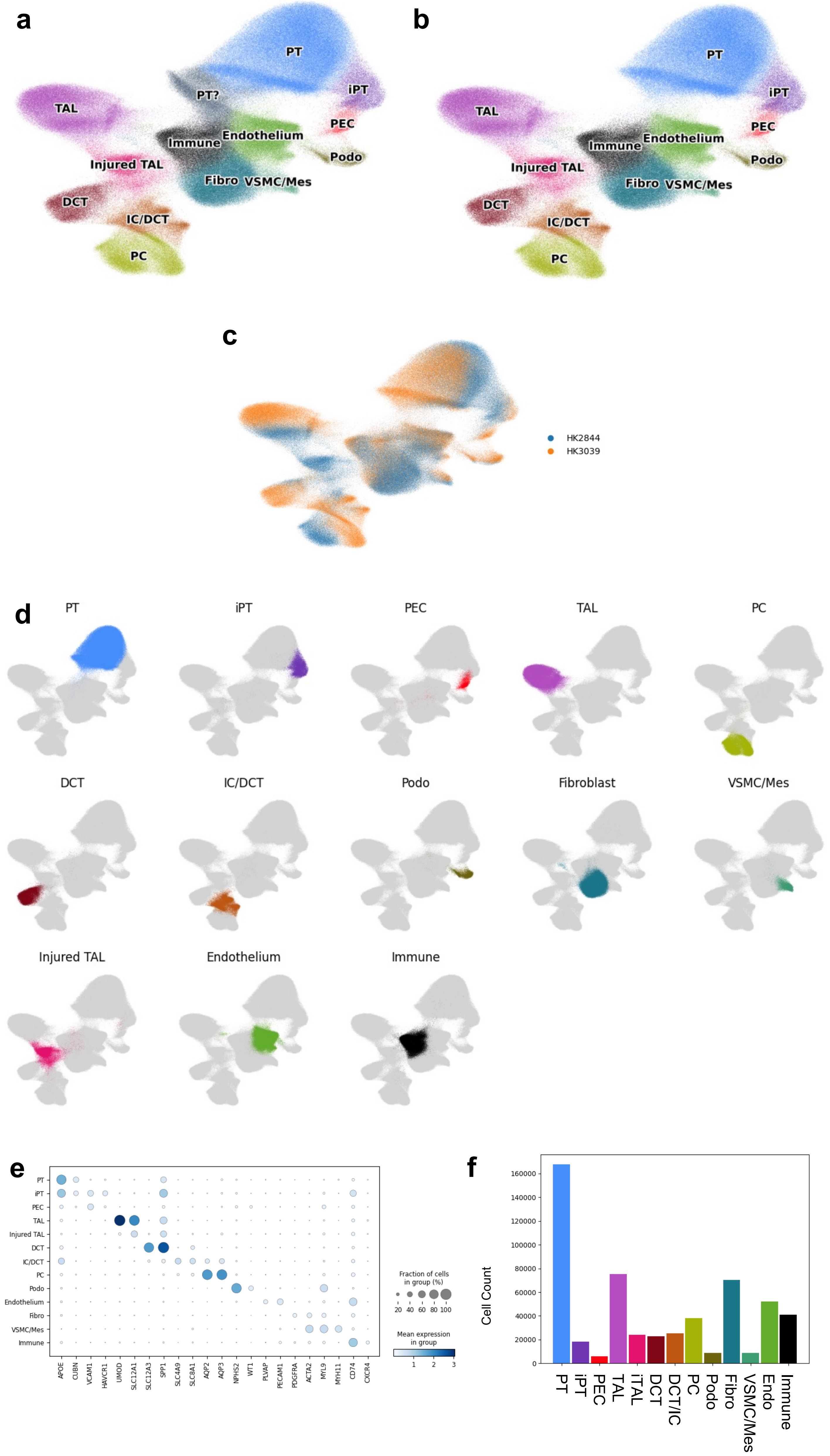


Supplementary Fig. 16

**Supplementary Fig. 16. CosMx Cell populations.**
**(a)** Original UMAP of all cells that passed QC along with their annotations.
**(b)** Cell Populations with annotatable markers of the CosMx data on UMAP.
**(c)** UMAP of all CosMx cells by sample.
**(d)** CosMx cell annotations across the UMAP with a single population being shown for a given UMAP demonstrating that these clusters are indeed relatively localized within the UMAP.
**(e)** Dot plot showing markers for each annotated CosMx population
**(f)** Annotation Frequency for each cell type.

VSMC/Mes; vascular smooth muscle cells/Mesangial, PEC; parietal epithelial cells, Podo; podocyte, PT; proximal tubule, Injured_PT; injured proximal tubule cells, TAL; thick ascending loop of Henle, iTAL; injured thick ascending limb, Fibro; fibroblast, PC; principal cell, Immune; immune cell, Endo; endothelial cell; DCT; distal convoluted tubule, DCT/IC;distal convoluted tubule/intercalated cell

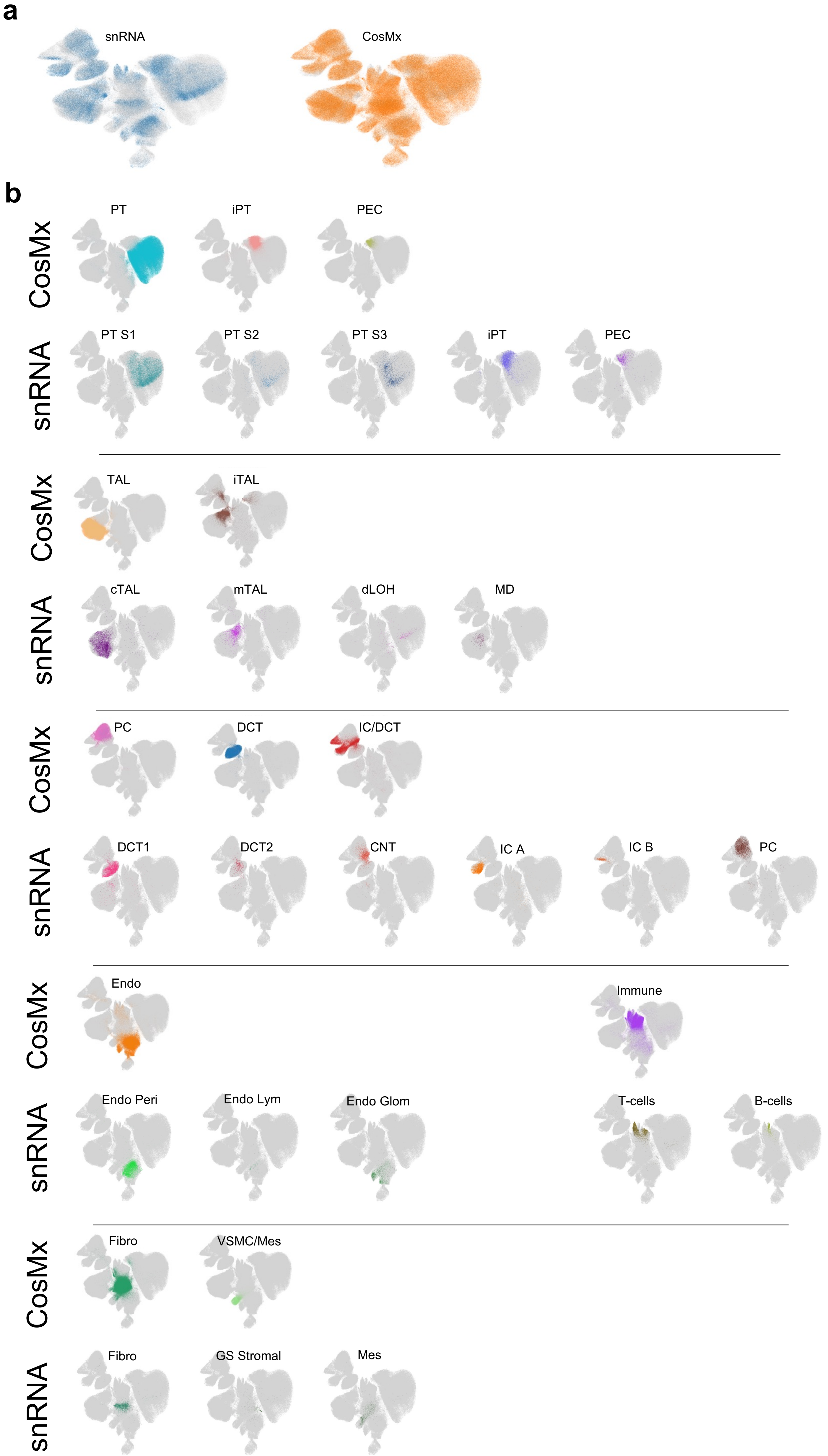


**Supplementary Fig. 17**

**Supplementary Fig. 17. CosMx SCVI integration with snRNAseq data.**
After integrating with the snRNAseq data, we compared annotations from our CosMx analysis and the original snRNAseq annotation.
**(a)** UMAP of integrated data demonstrating technology type of each cell within the UMAP.
**(b)** Comparison of CosMx annotations and snRNAseq annotations, demonstrating concordance of location within the integrated UMAP.

VSMC/Mes; vascular smooth muscle cells/Mesangial, PEC; parietal epithelial cells, Podo; podocyte,  PT; proximal tubule, Injured_PT; injured proximal tubule cells, TAL; thick ascending loop of Henle, iTAL; injured thick ascending limb, Fibro; fibroblast, PC; principal cell, Immune; immune cell, Endo; endothelial cell; DCT; distal convoluted tubule, DCT/IC;distal convoluted tubule/intercalated cell, IC A; intercalated alpha cells, IC B intercalated beta cells, CNT; connecting tubule, Endo_GC; endothelial cells of glomerular capillary tuft, Endo_peritubular; endothelial cells of peritubular vessels, Endo_lymphatic; endothelial cells of lymphatic vessels, Mes; meseangial cells, GS_Stromal; glomerulosclerosis-specific stromal cells, VSMC/Pericyte; vascular smooth muscle cells/pericyte,

a

b

c

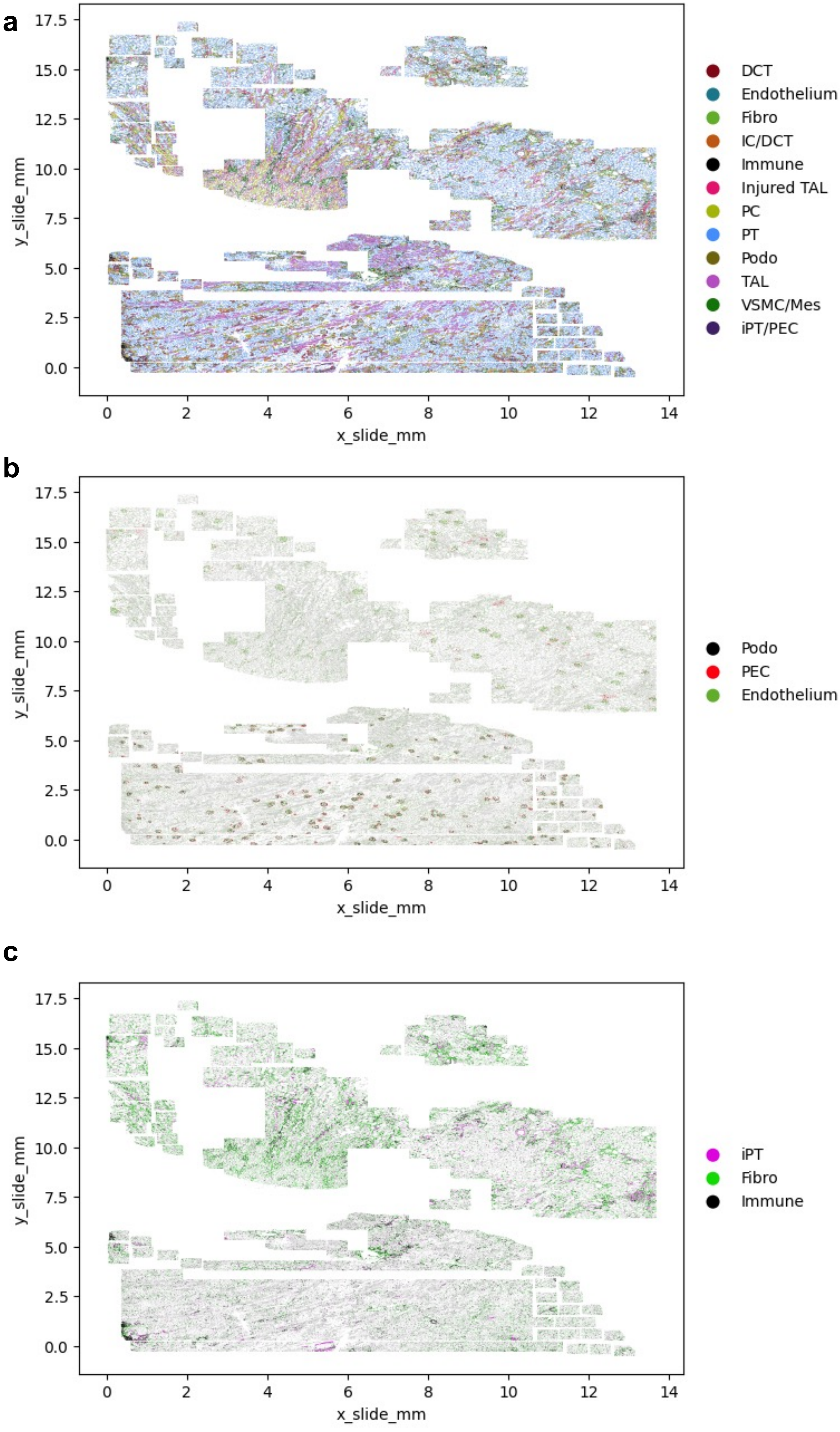


**Supplementary Fig. 18**

**Supplementary Fig. 18 Location of CosMx annotated cell types within the slide.**
**(a)** Location of annotated cell types within the two tissue sections.
**(b)** Location of glomerular cell subtypes
**(c)** Location of injured PT, fibroblasts and immune cells

VSMC/Mes; vascular smooth muscle cells/Mesangial, PEC; parietal epithelial cells, Podo; podocyte, PT; proximal tubule, Injured_PT; injured proximal tubule cells, TAL; thick ascending loop of Henle, iTAL; injured thick ascending limb, Fibro; fibroblast, PC; principal cell, Immune; immune cell, Endo; endothelial cell; DCT; distal convoluted tubule, DCT/IC;distal convoluted tubule/intercalated cell

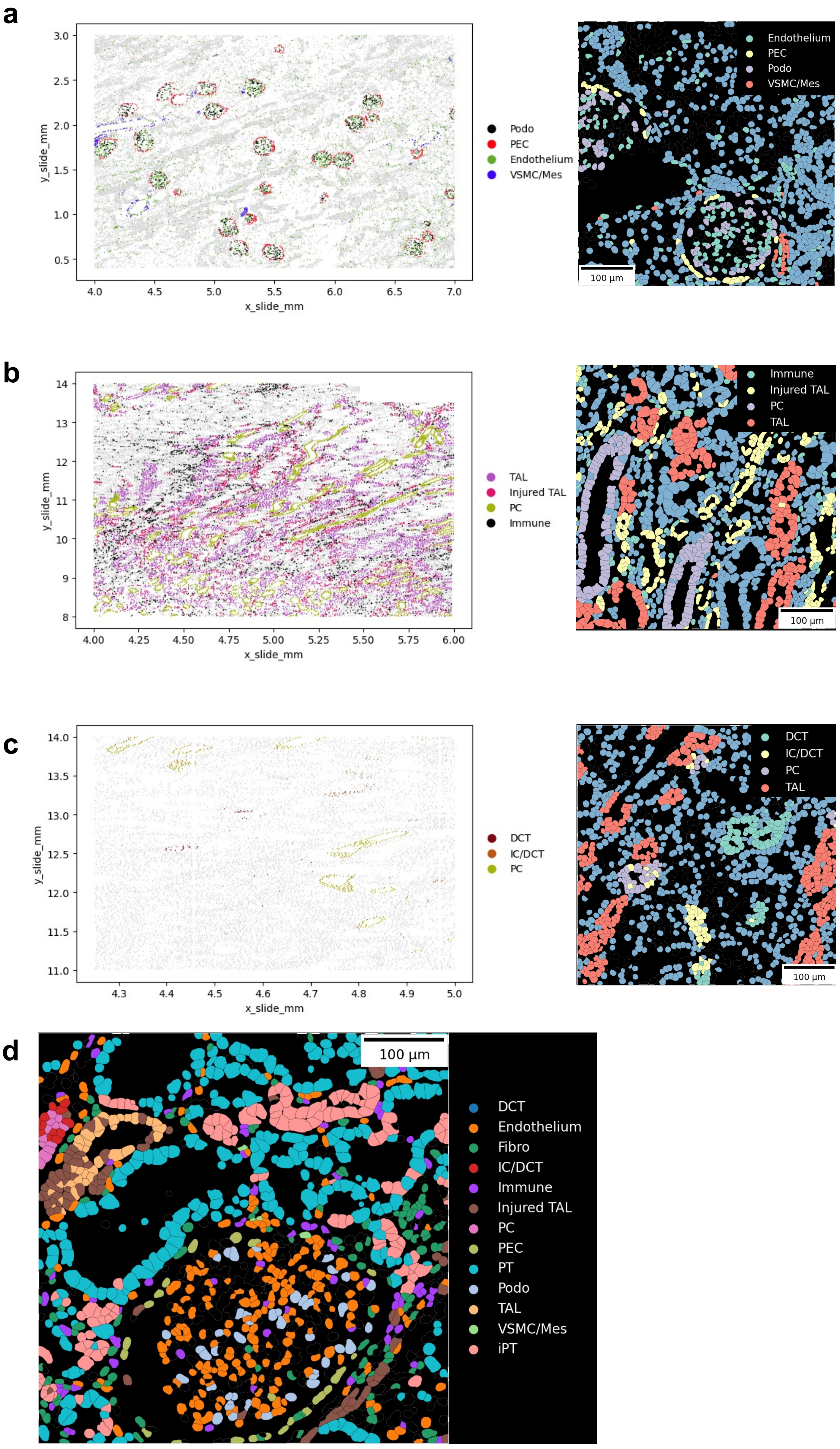


**Supplementary Fig. 19**

**Supplementary Fig. 19 Microanatomy of the CosMx slide**
**(a)** Location of glomerular cell types within a subsection of tissue, and in a single field of view (right)
**(b)** Location of injured thick ascending limb, healthy injured thick ascending limb, principal cells and immune cell types within a subsection of tissue, and in a single field of view (right).
**(c)** Location of distal nephron cell subtypes within a subsection of tissue and in a single field of view (right)
**(d)** Single field of view showing many cell types.

VSMC/Mes; vascular smooth muscle cells/Mesangial, PEC; parietal epithelial cells, Podo; podocyte, PT; proximal tubule, Injured_PT; injured proximal tubule cells, TAL; thick ascending loop of Henle, iTAL; injured thick ascending limb, Fibro; fibroblast, PC; principal cell, Immune; immune cell, Endo; endothelial cell; DCT; distal convoluted tubule, DCT/IC;distal convoluted tubule/intercalated cell

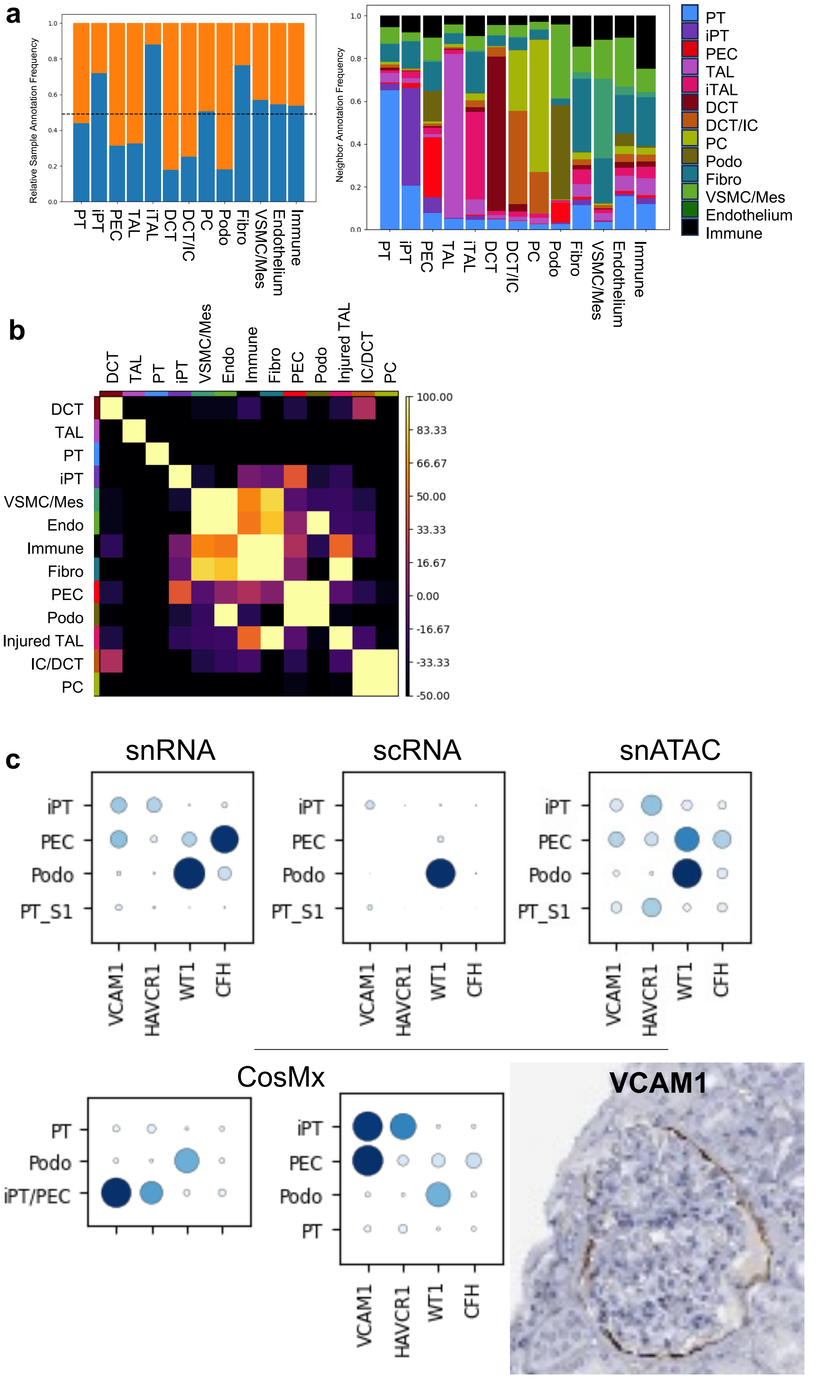


Supplementary Fig. 20

**Supplementary Fig. 20. Neighborhood characteristics of CosMx slide**

**(a)** Relative type cell frequency between each sample. Orange indicates HK3039 (healthy) and blue indicates HK2844 (diseased), and frequency of neighbor annotations for each cell type for a 20-micron neighborhood (right).

**(b)** Neighborhood enrichment by permuting annotations for the 20-micron neighborhood size. Lighter color indicates higher enrichment and co-localization of a given population.

**(c)** Dot plots for iPT, PEC, Podocytes and PT cells expression of iPT and PEC markers across genomics modalities and protein staining of VCAM1 in PECs from the human protein atlas: https://www.proteinatlas.org/.

VSMC/Mes; vascular smooth muscle cells/Mesangial, PEC; parietal epithelial cells, Podo; podocyte, PT; proximal tubule, Injured_PT; injured proximal tubule cells, TAL; thick ascending loop of Henle, iTAL; injured thick ascending limb, Fibro; fibroblast, PC; principal cell, Immune; immune cell, Endo; endothelial cell; DCT; distal convoluted tubule, DCT/IC;distal convoluted tubule/intercalated cell.

a

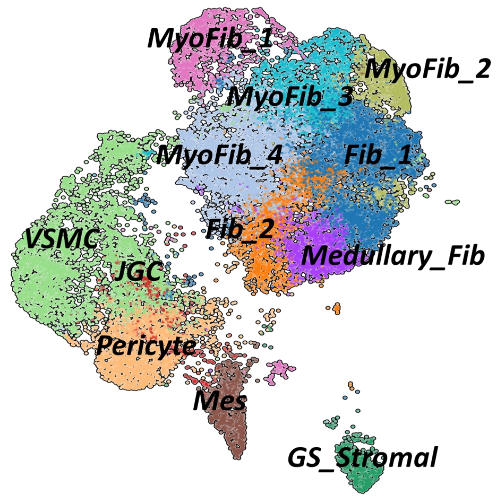

MyoFib_1
MyoFib_2
MyoFib_3
MyoFib_4
Fib_1
Fib_2
VSMC
JGC
Medullary_Fib
Pericyte
Mes
GS_Stromal


b

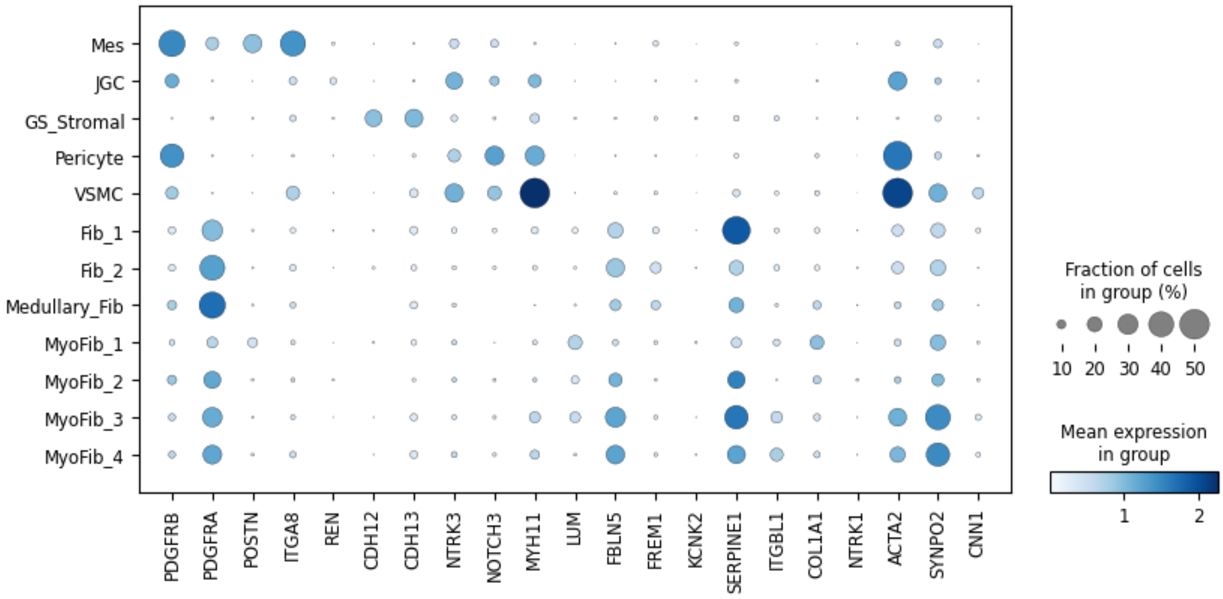

Mes
JGC
GS_Stromal
Pericyte
VSMC
Fib_1
Fib_2
Medullary_Fib
MyoFib_1
MyoFib_2
MyoFib_3
MyoFib_4
PDGFRB
PDGFRA
POSTN
ITGA8
REN
CDH12
CDH13
NTRK3
NOTCH3
MYH11
LUM
FBLN5
FREM1
KCNK2
SERPINE1
ITGBL1
COL1A1
NTRK1
ACTA2
SYNPO2
CNN1
Fraction of cells in group (%)
10 20 30 40 50
Mean expression in group
1 2


c

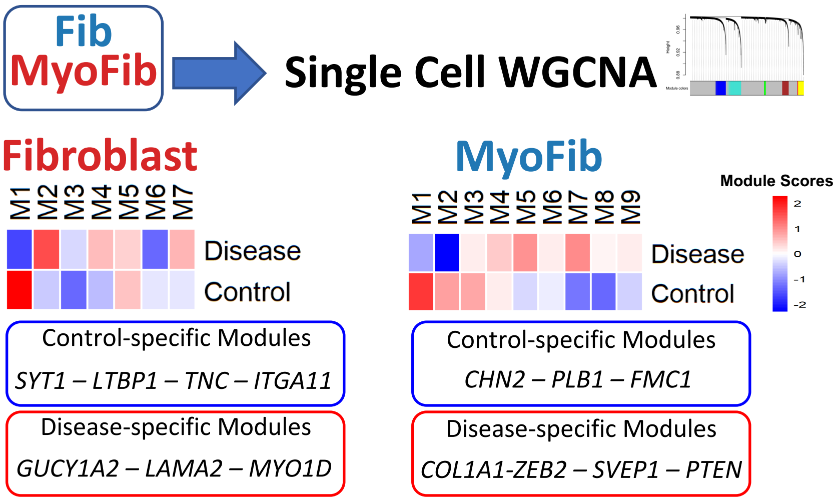

Fib
MyoFib
Single Cell WGCNA
Fibroblast
M1 M2 M3 M4 M5 M6 M7
Disease
Control
Control-specific Modules
SYT1 – LTBP1 – TNC – ITGA11
Disease-specific Modules
GUCY1A2 – LAMA2 – MYO1D
MyoFib
M1 M2 M3 M4 M5 M6 M7 M8 M9
Disease
Control
Module Scores
2
1
0
-1
-2
Control-specific Modules
CHN2 – PLB1 – FMC1
Disease-specific Modules
COL1A1-ZEB2 – SVEP1 – PTEN

**Supplementary Fig. 21. Sub-clustering of stromal cells in the integrated dataset**
**(a)** UMAP of 41,422 stromal cells in the integrated dataset.
**(b)** The dot plots of marker genes used for cell type annotation. The size of the dot indicates the percent positive cells, and the darkness of the color indicates average expression.
**(c)** The single cell Weighted correlation network analysis (WGCNA) on fibroblast and myofibroblast identified specific modules in control and disease samples. The heatmaps shows the identified modules in each cell types and their scores. The lower panel shows the representative genes with high scores in each cell type.

Mes; Mesangial cells, GS_Stromal; glomerulosclerosis associated stromal cells, JGC; juxta-glomerular cells, VSMC; vascular smooth muscle cell, Fib; fibroblast, MyoFib; myofibroblast.

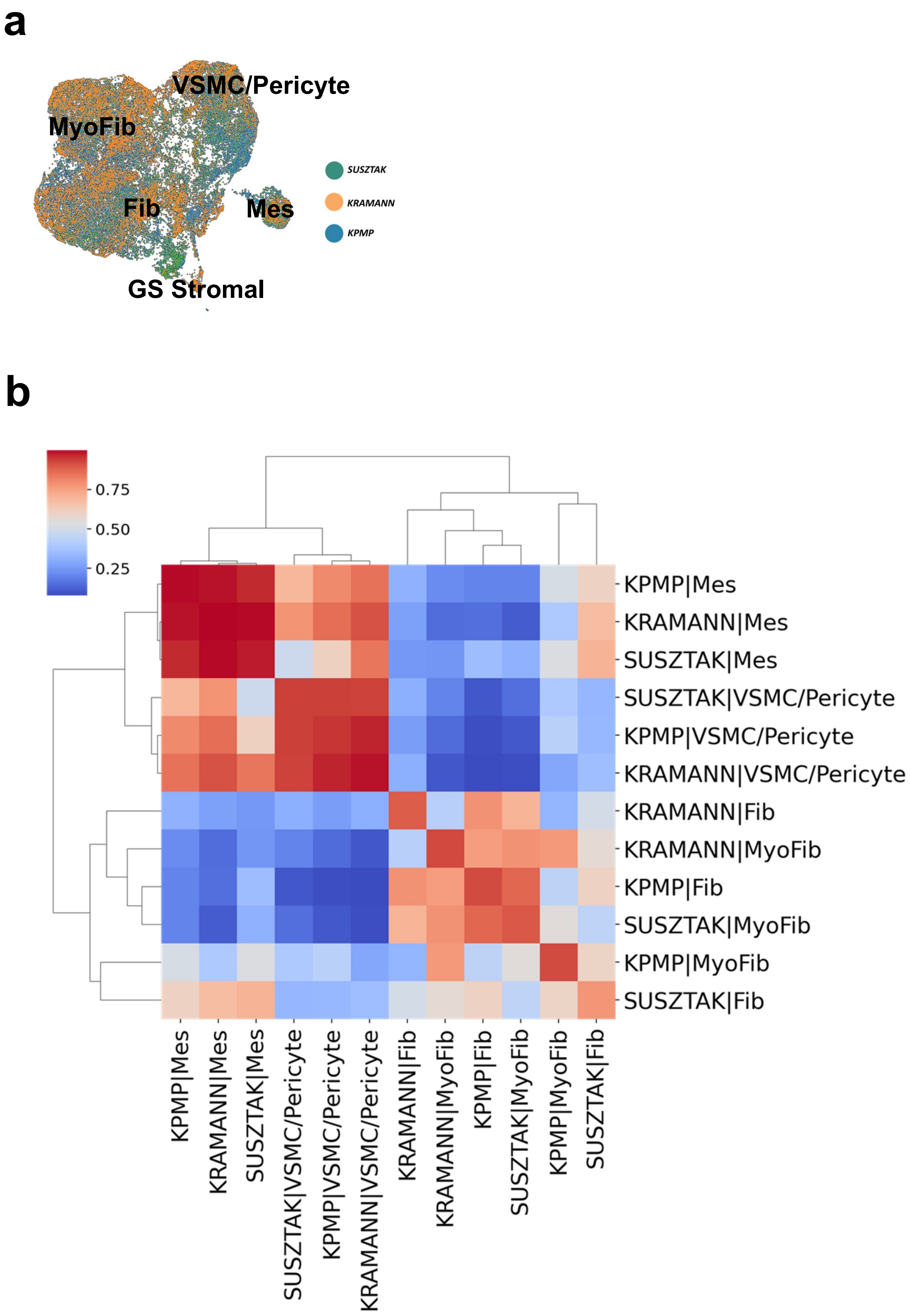


Supplementary Fig. 22

**Supplementary Fig. 22. The integration of stromal cells with prior studies**

**(a)** UMAP of integration of 86,481 stromal cells of current study prior papers by KPMP and Kramann *et al.* The color indicates the data source.

**(b)** Heatmap display of MetaNeighbor analysis of the similarities of the identified stromal cell types. The heatmap displays the Area Under the Receiver Operating Characteristic (AUROC) scores among different cell subtypes in three different sources of datasets, which were determined using a set of highly variable genes (HVG). Dendrograms were created via hierarchical clustering, based on Euclidean distances, and used average linkage for this purpose. Each row and column is a cell type in the source data.

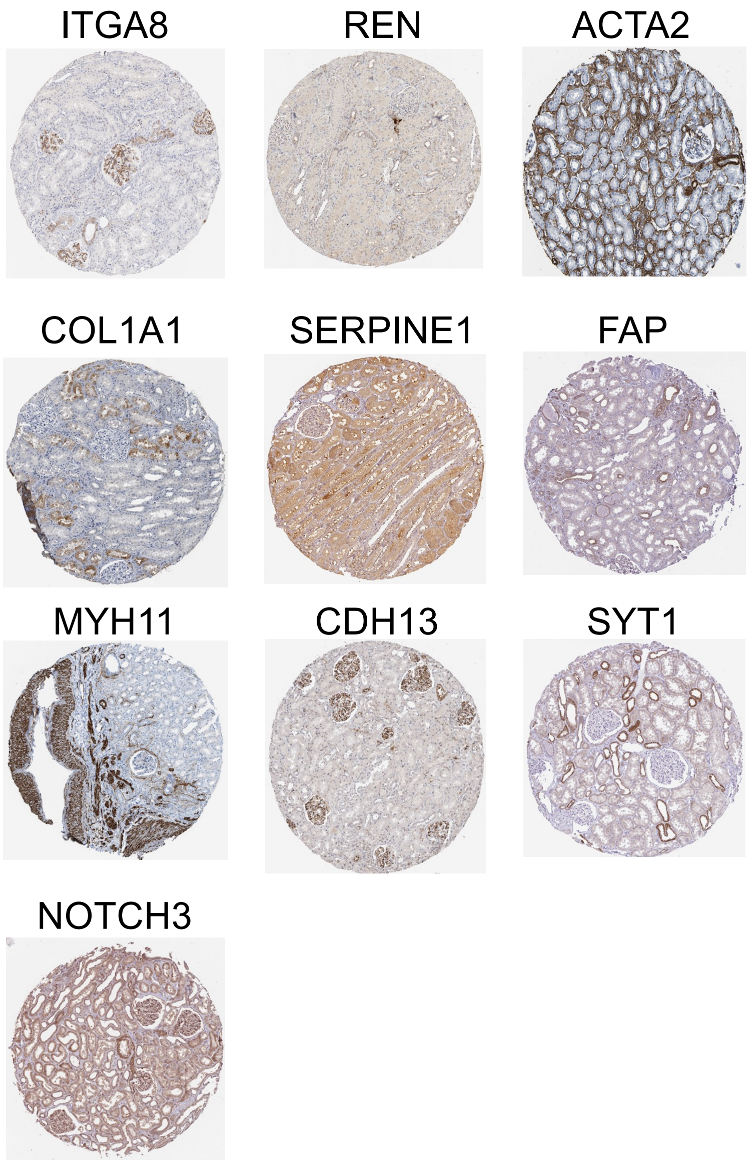


**Supplementary Fig. 23**

**Supplementary Fig. 23. Expression of stromal markers in the Human** Protein Atlas kidney samples. Images were taken from https://www.proteinatlas.org/.

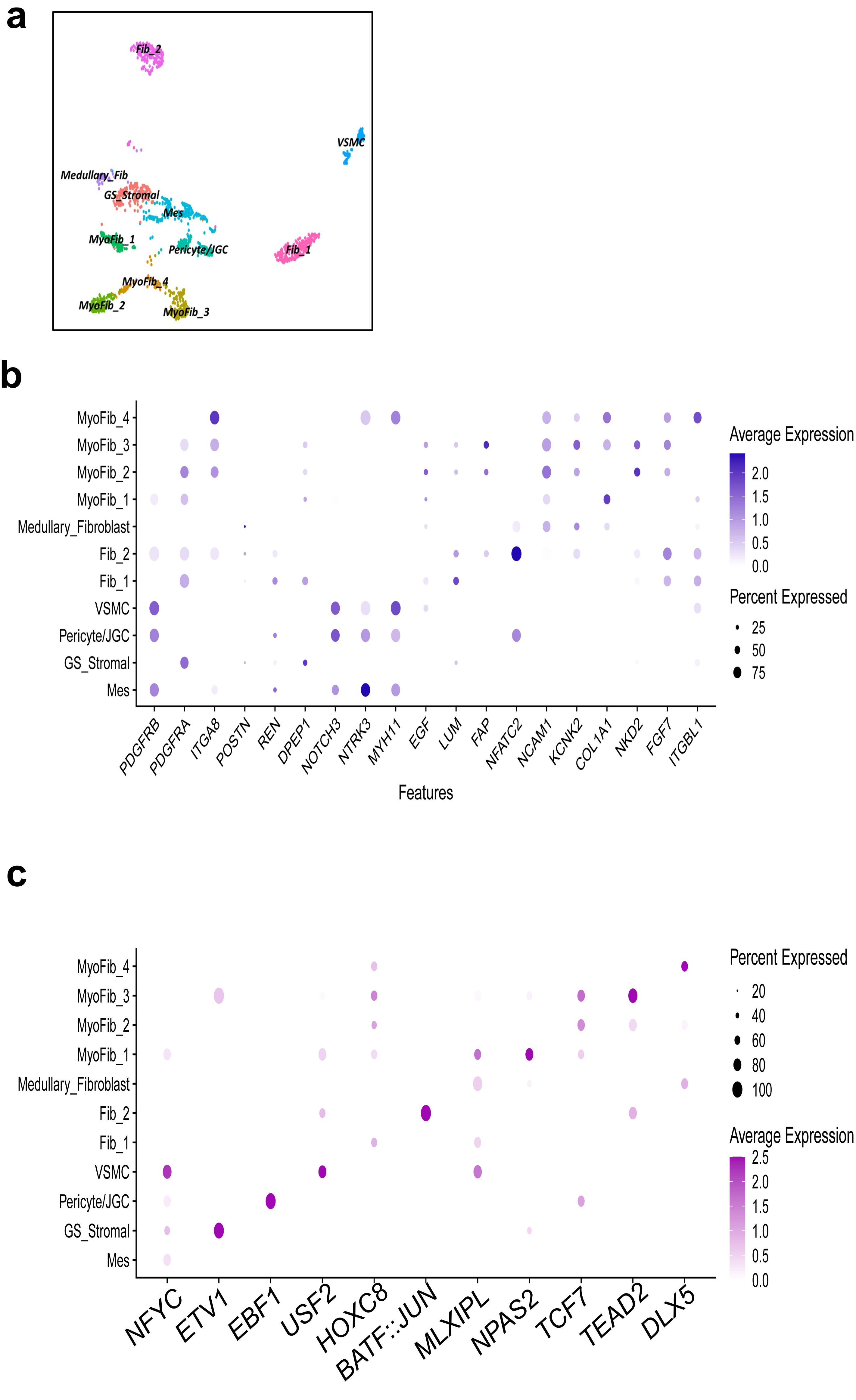


**Supplementary Fig. 24**

**Supplementary Fig. 24. Stromal sub-clustering in human snATACseq data.**
**(a)** UMAP of 5,822 stromal cells in human snATACseq dataset.
**(b)** The dot plot of gene markers used to annotate the different stromal cell types in snATACseq dataset. The dot plot shows the gene activity in different cell types. The size of the dot indicates the percent positive cells, and the darkness of the color indicates average expression.
**(c)** The dot plot of motif activity of enriched motifs in each identified stromal cluster. The size of the dot indicates the percent positive cells, and the darkness of the color indicates average expression.

Mes; Mesangial cells, GS_Stromal; glomerulosclerosis associated stromal cells, JGC; juxta-glomerular cells, VSMC; vascular smooth muscle cell, Fib; fibroblast, MyoFib; myofibroblast.

**a**

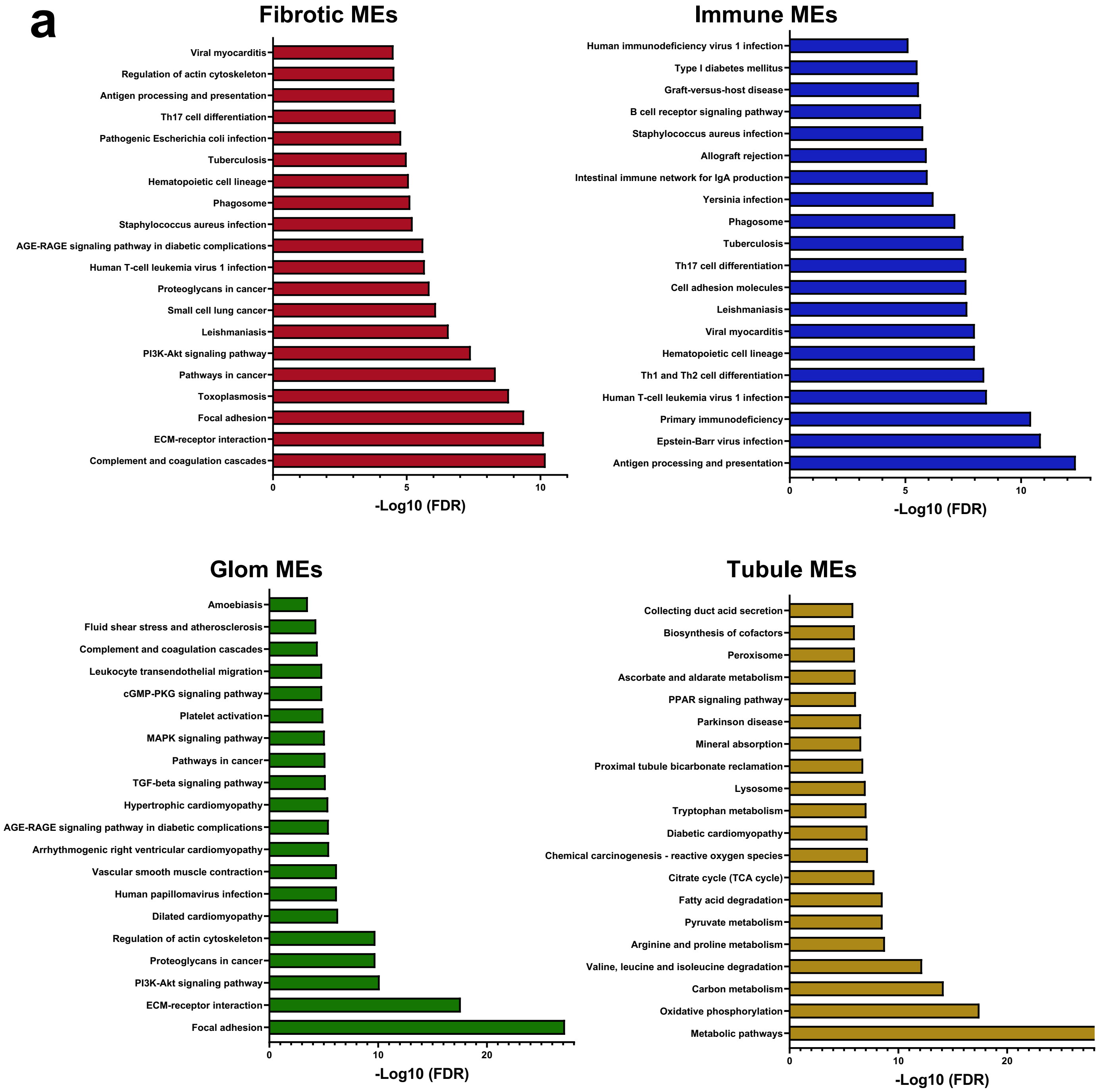


**b**

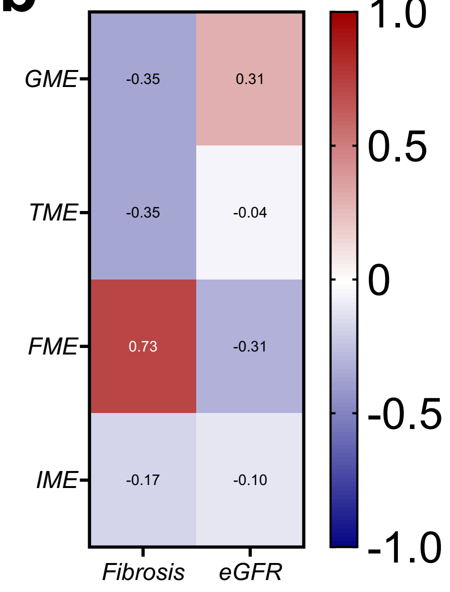


**Supplementary Fig. 25**

**Supplementary Fig. 25. Gene ontology analysis of microenvironment gene signature**

**(a)** Gene ontology analysis (DAVID) of the microenvironment gene signature. The X axis shows the -Log10 (FDR) of the identified pathways and Y axis lists the pathways.

**(b)** Person correlation of the number of spots assigned to each microenvironment and eGFR and fibrosis.

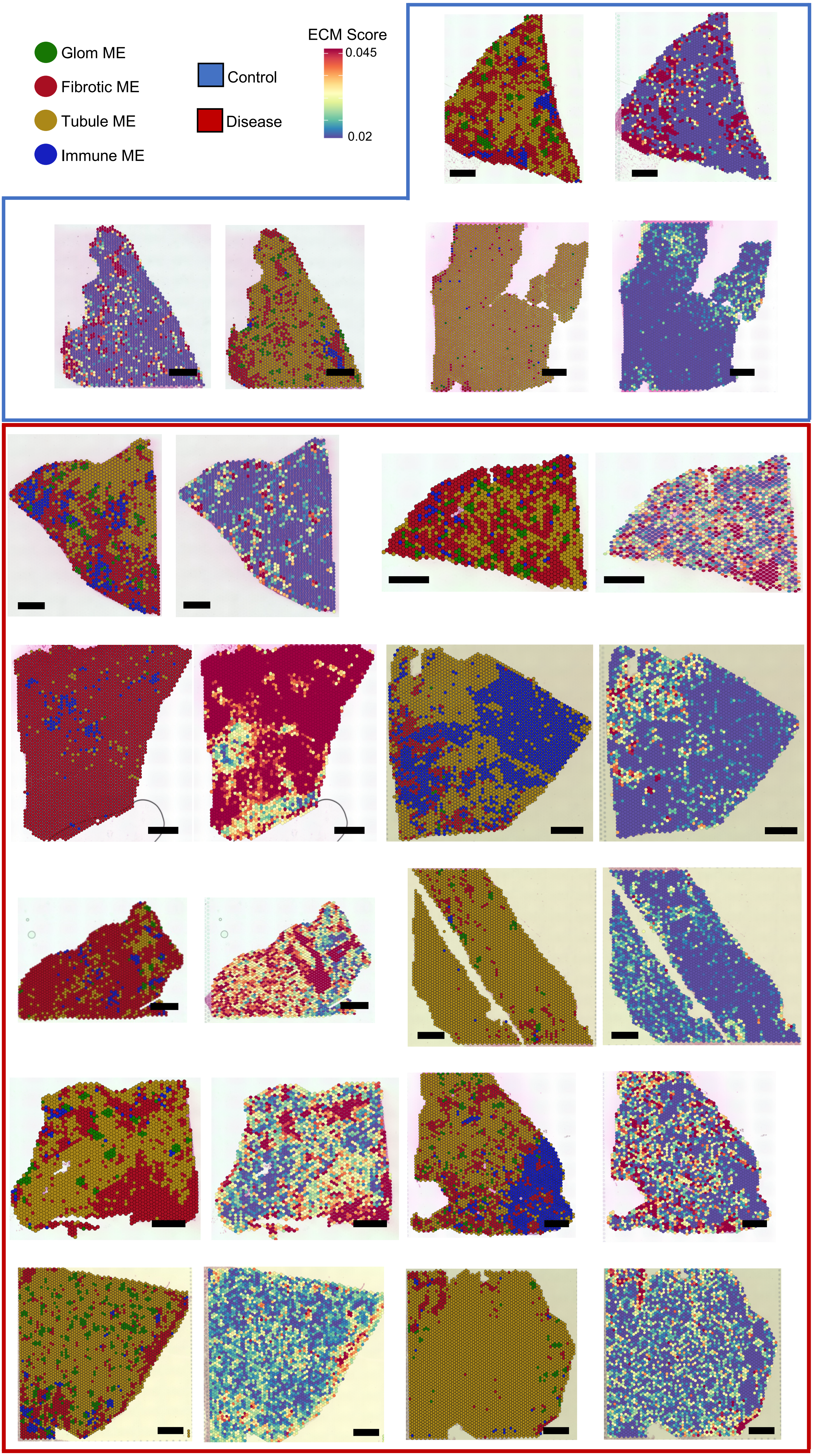


Supplementary Fig. 26

**Supplementary Fig. 26. The spatial microenvironments in the human kidney.** Four spatial microenvironments were identified in human kidney samples using matrix factorization: glomerular (green), tubule (brown), fibrotic (red) and immune (blue). The ECM score (calculated by aggregate expression collagen and other glycoprotein, and proteoglycan) in Visium data shown next to the tissue microenvironment information. Scale bar is 1 mm in length.

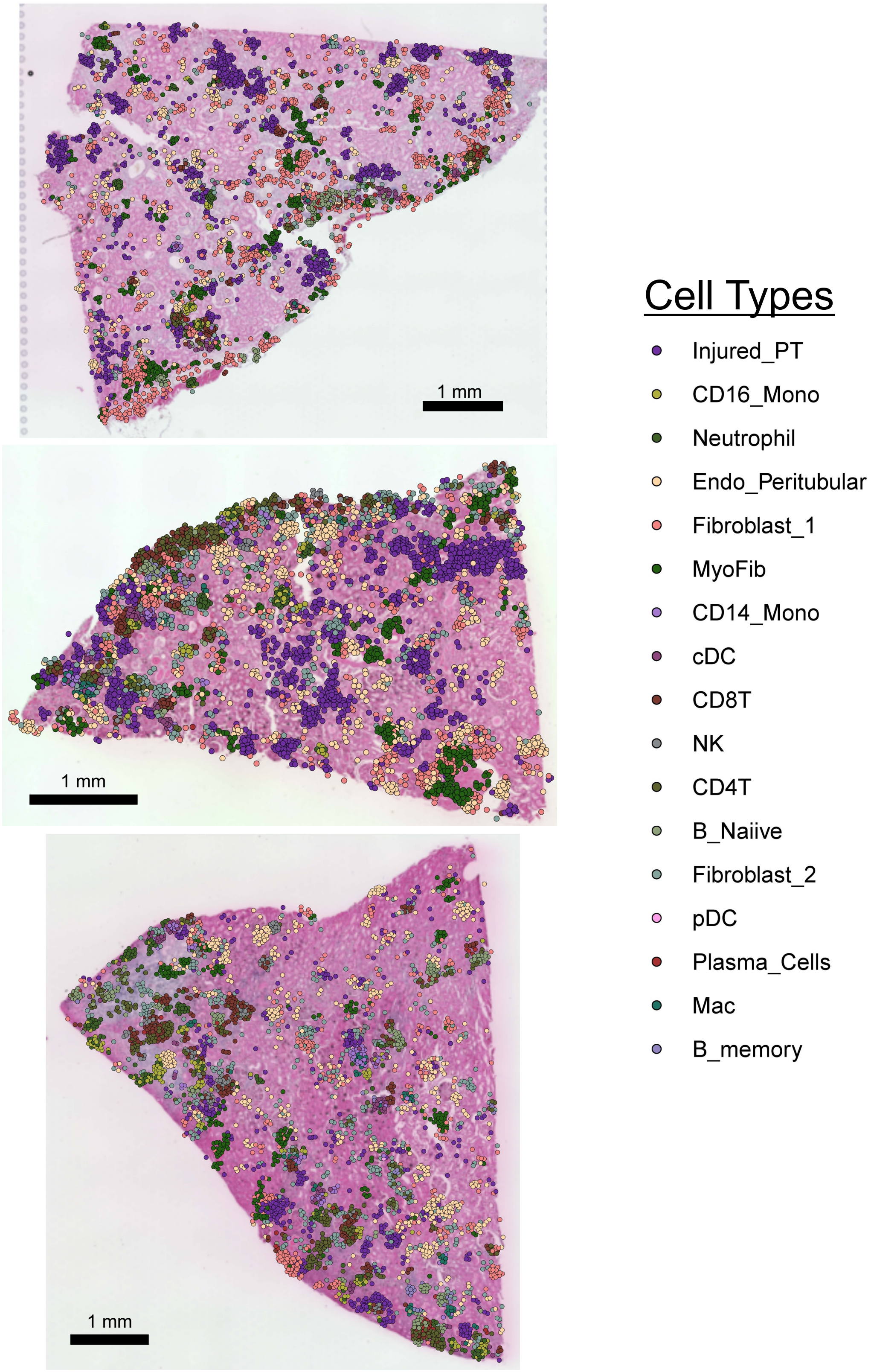


**Supplementary Fig. 27**

**Supplementary Fig. 27. Different cell types in kidney microenvironment.** Left panel kidney microenvironment in tissue samples, right panel shows cell abundance mapped using Cell2location (endothelial cell, injured_PT cells, stromal cells and immune cells)

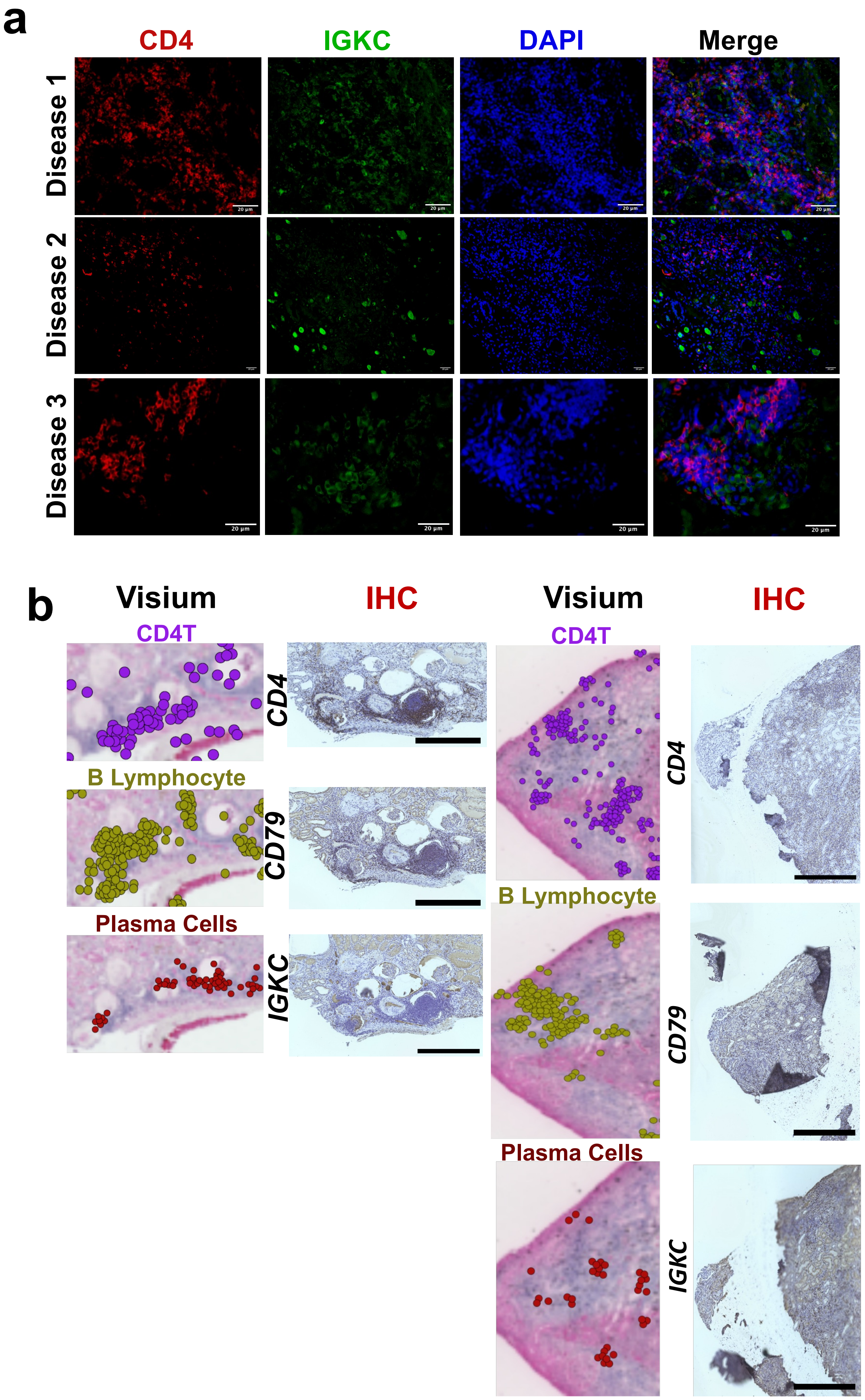


Supplementary Fig. 28

**Supplementary Fig. 28. Immunostaining of the lymphocyte and plasma cells in regions of fibrosis-associated immune cells aggregates.**

**(a)** Immunofluorescent staining of T-cell markers (CD4) and plasma cells (IGKC) in diseased human kidney samples.

**(b)** Spatial RNAseq (10x Visium with CellTrek imputation) and immunohistochemistry staining of T-cell, B-cell, and plasma cells in diseased human kidney samples in the regions of fibrosis-associated immune cells aggregates. Scale bar is 1 mm in length.

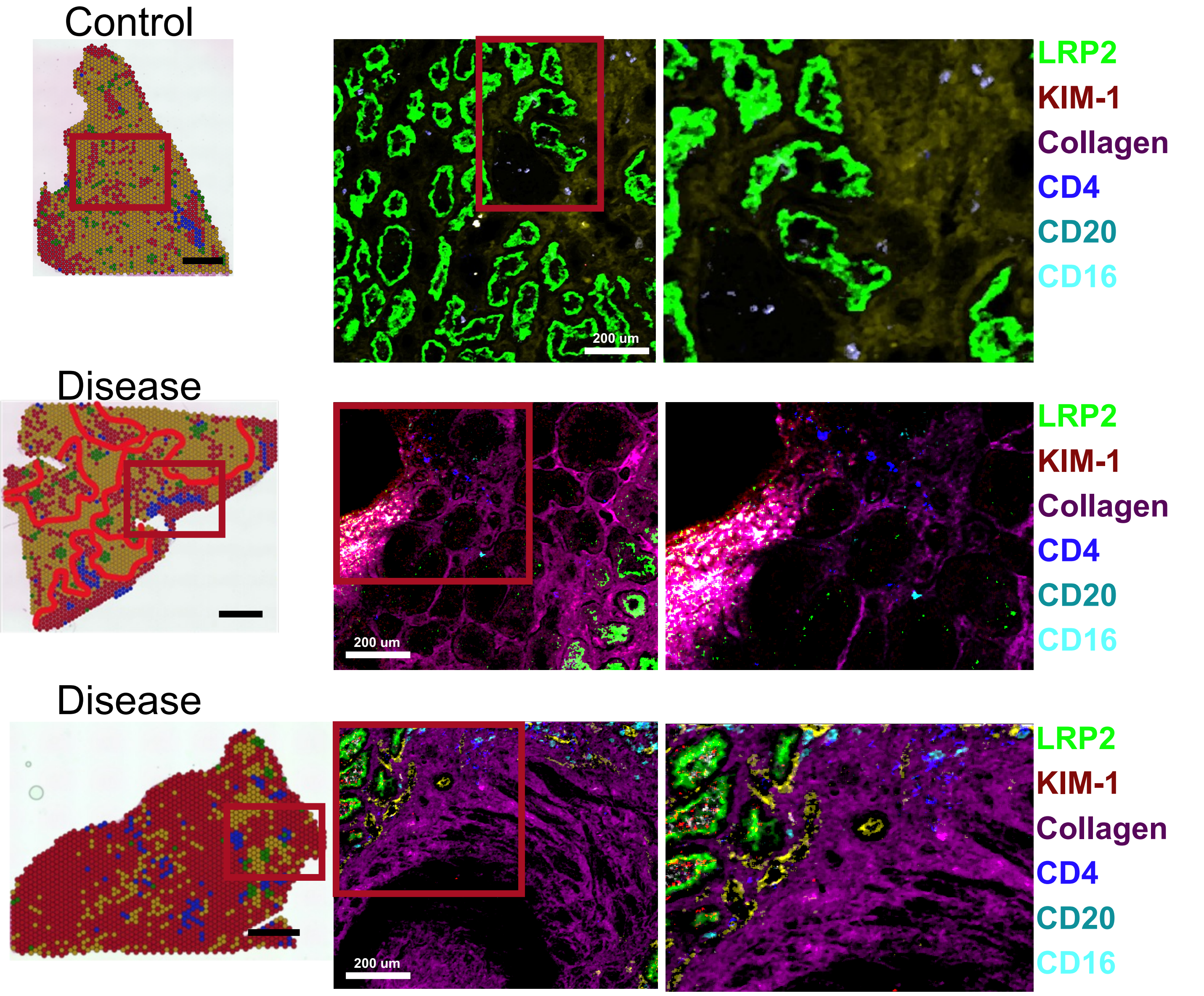


**Supplementary Fig. 29**

**Supplementary Fig. 29. *In situ* mass spectrometry of the control, and diseased samples to verify cell types in fibrotic microenvironments.**
The left panel shows the Visium 10x data cellular microenvironment. The right panel shows the IMC staining of the area, markers and colors are listed on the right. Scale bar is 1 mm unless otherwise indicated.

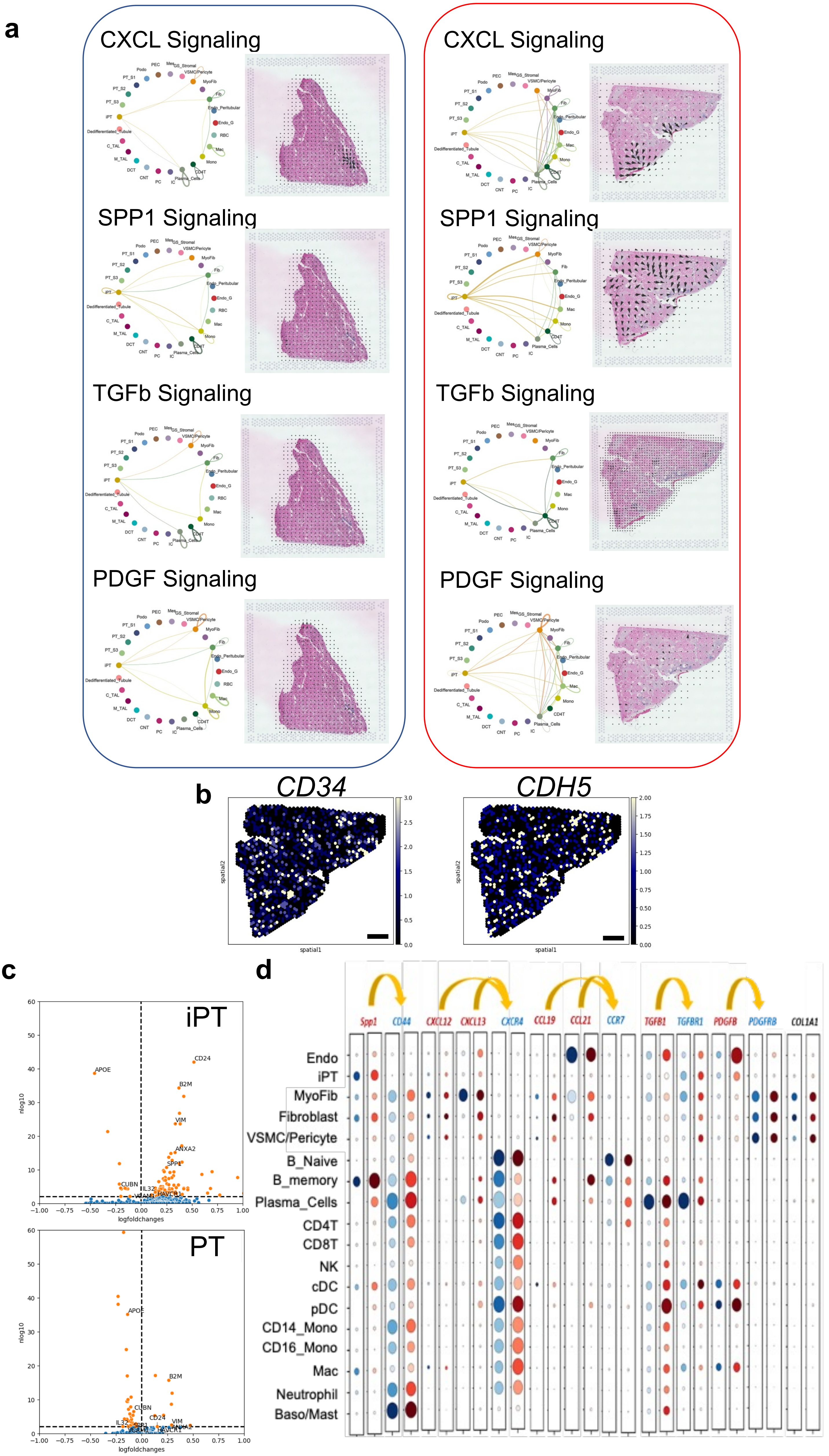


Supplementary Fig. 30

**Supplementary Fig. 30. Cell-cell interaction analysis in the spRNAseq dataset in fibrotic microenvironment.**

**(a)** Weighted total interaction strength of the CXCL, SPP1, TGFb, and PDGF pathways in control and diseased samples in the fibrotic microenvironment (left panel). The spatial location of the identified cell-cell communications pathways (*CXCL*, *SPP1*, *TGFb*, and *PDGF*) in control and diseased sample in the fibrotic microenvironment were shown in the right panel. The arrows indicate the source and targets of the identified pathways.

**(b)** Expression of *CD34* and *CDH5* as the markers of high endothelial venules (HEVs) in the fibrotic microenvironments. Scale bar is 1 mm in length.

**(c)** Volcano plot of differentially expressed genes from CosMx data. Cells with an immune neighbor within 20 microns were compared against cells without an immune neighbor for both the PT and iPT population. Logfold change > 0 indicates increased expression in cell with an immune neighbor, while logfoldchange < 0 indicates increased expression in cells without an immune neighbor. Log10(pvalue) is indicated by the y-axis. Genes with an adjusted p-value < 0.01 are marked in orange.

**(d)** The dot plot of expression of ligands and receptors in regions of FME in integrated sn/scRNAseq and snATACseq data. The size of the dot indicates the percent positive cells and the darkness of the color indicates average expression (right panel). The gray indicates control and red indicates diseased group.

Endo; endothelial cells, MyoFib; myofibroblast, Fib; fibroblast, VSMC/Pericyte; vascular smooth muscle cells and pericyte, Podo; podocyte, Mes; mesangial cells, PT; proximal tubule cells, iPT; injured proximal tubule cells, LOH; loop of Henle, DCT; distal convoluted tubule, CNT; connecting tubule, PC; principal cells of the collecting duct.

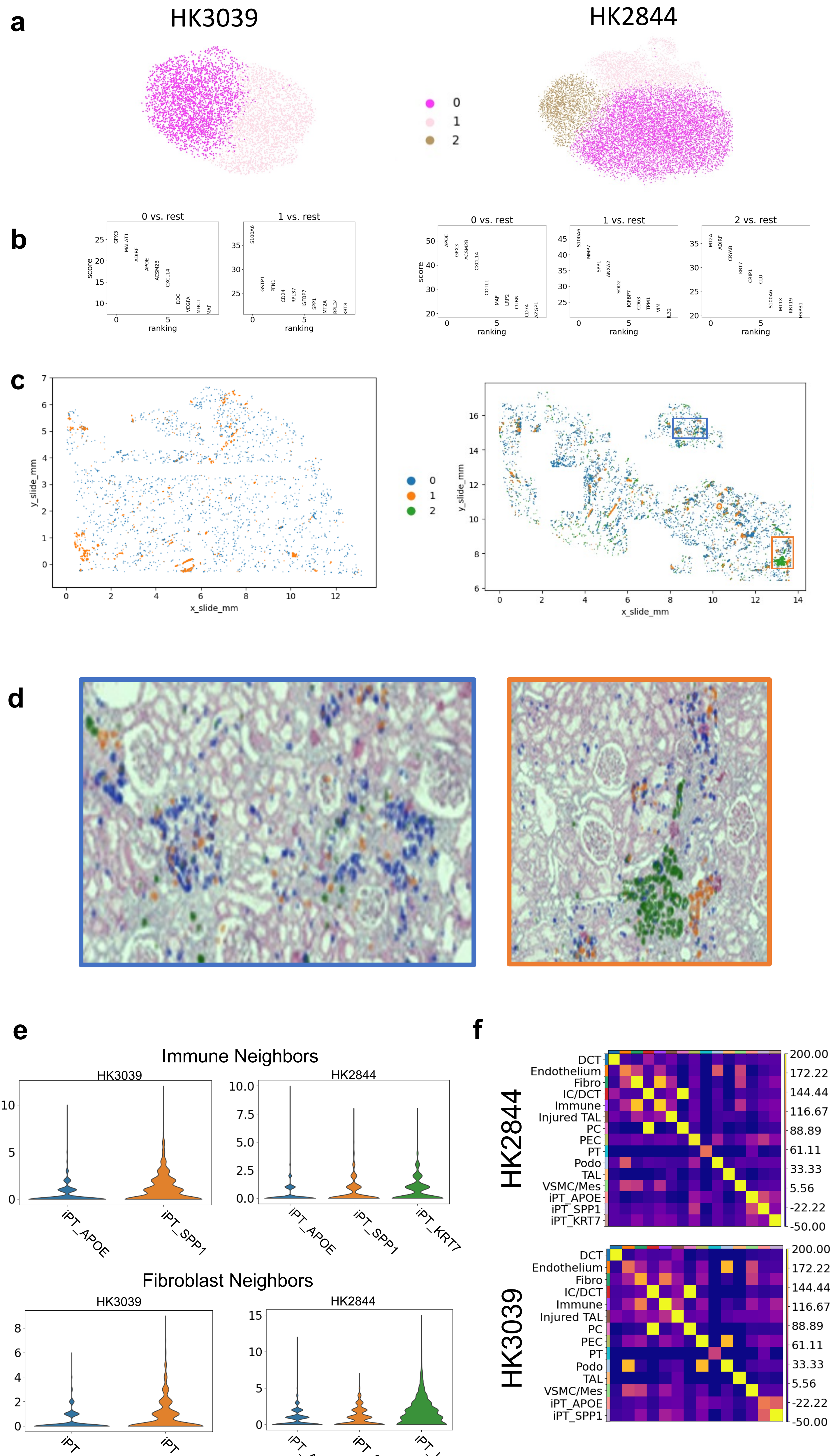


**Supplementary Fig. 31**

**Supplementary Fig. 31 Spatial characteristics of injured PT subclusters on CosMx.**

**(a) The CosMx iPT population for each sample was subclustered using a leiden algorithm with resolution of 0.3.**

**(b) The top 10 differentially expressed genes for each subcluster.**

**(c) iPT subcluster localization within the entire slide. Views of specific regions indicated by inset boxes are shown in Supplementary Fig. 31d.**

**(d) iPT subclusters visualized on H&E. Subset images showing populations on H&E. Blue cells correspond with cluster 0 (iPT_APOE), orange with cluster 1 (iPT_SPP1) and green with cluster 2 (iPT_KRT7).**

**(e) Frequency of immune and fibroblast neighbors for each iPT subtype within each sample within 20 microns is shown below. We performed testing using a Wilcoxon rank sum test between each population within a sample. These subtypes had significantly different immune neighbors and fibroblast neighbors with each sample. HK3039 fibroblasts: iPT_APOE vs iPT_SPP1, p-value = 7e-39. HK3039 immune cells: iPT_APOE vs iPT_SPP1, p-value = 4e-67. HK2844 fibroblasts iPT_APOE vs iPT_SPP1, p-value = 9e-9, iPT_KRT7 vs iPT_SPP1, p-value = 9e-46. HK2844 immune cells iPT_APOE vs iPT_SPP1, p-value = 3e-16, iPT_KRT7 vs iPT_SPP1, p-value = 9e-11.**

**(f) iPT subcluster neighborhood enrichment within a 20-micron neighborhood size. Lighter color indicates higher enrichment and co-localization of a given population.**

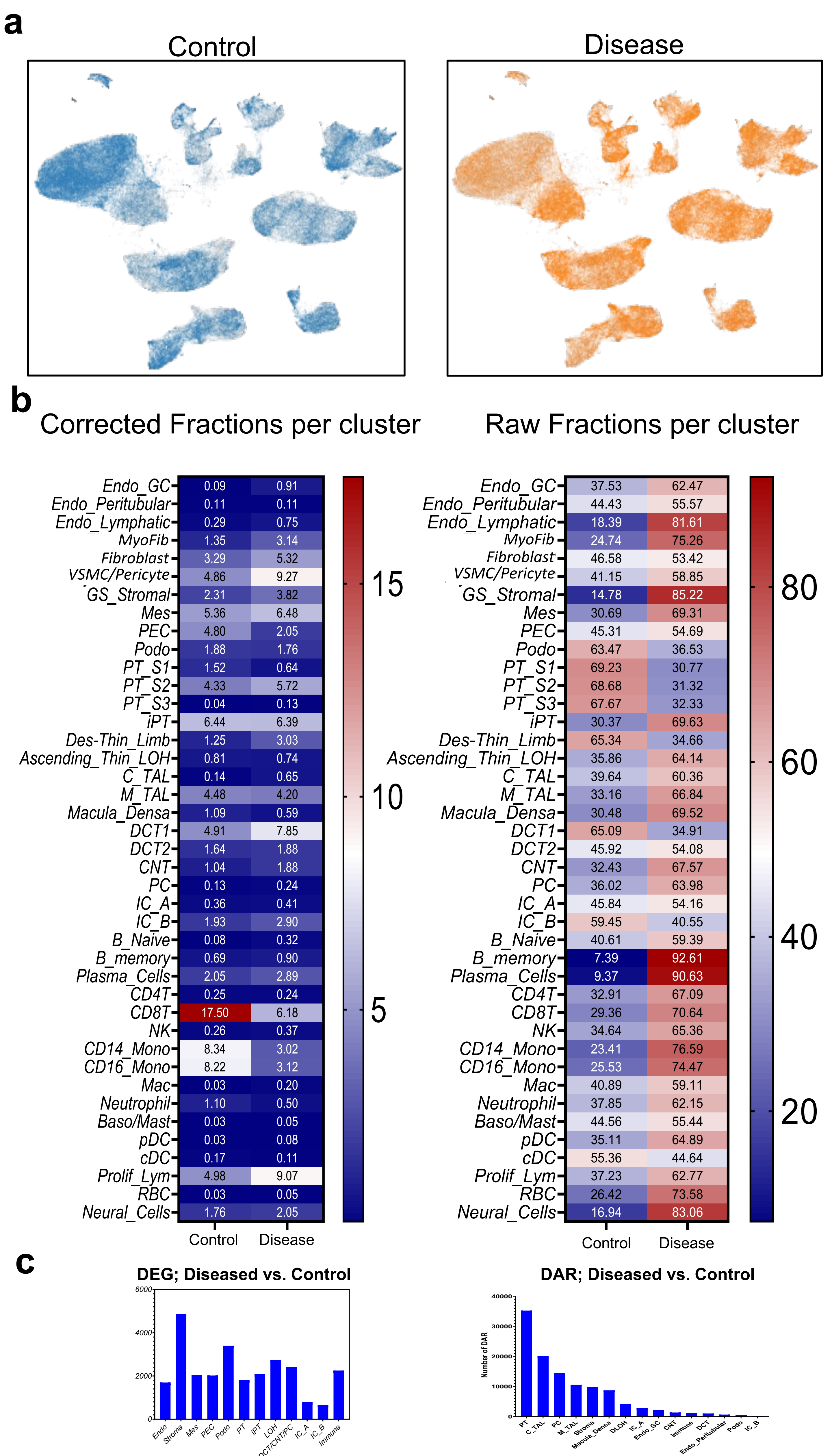


Supplementary Fig. 32

**Supplementary Fig. 32. Comparison of control and diseased samples in the integrated sn/scRNAseq and snATACseq.**

**(a)** UMAP distribution of control and diseased samples in the integrated snRNA/scRNAseq dataset.

**(b)** Heatmaps of cell type distribution in the control and diseased conditions.

**(c)** Heat of gene expression changes correlating with eGFR and fibrosis in each cell type. Red indicates larger number of correlating cells.

Endo; endothelial cells Mes; mesangial cells, Stroma; stromal cells, PEC; parietal epithelial cells, Podo; podocyte, PT; proximal tubule cells, Injured_PT; injured proximal tubule cells, LOH; loop of Henle, DCT; distal convoluted tubule, CNT; connecting tubule cells, PC; principal cells of collecting duct, IC_A; Type alpha intercalated cells, IC_B; Type beta intercalated cells.

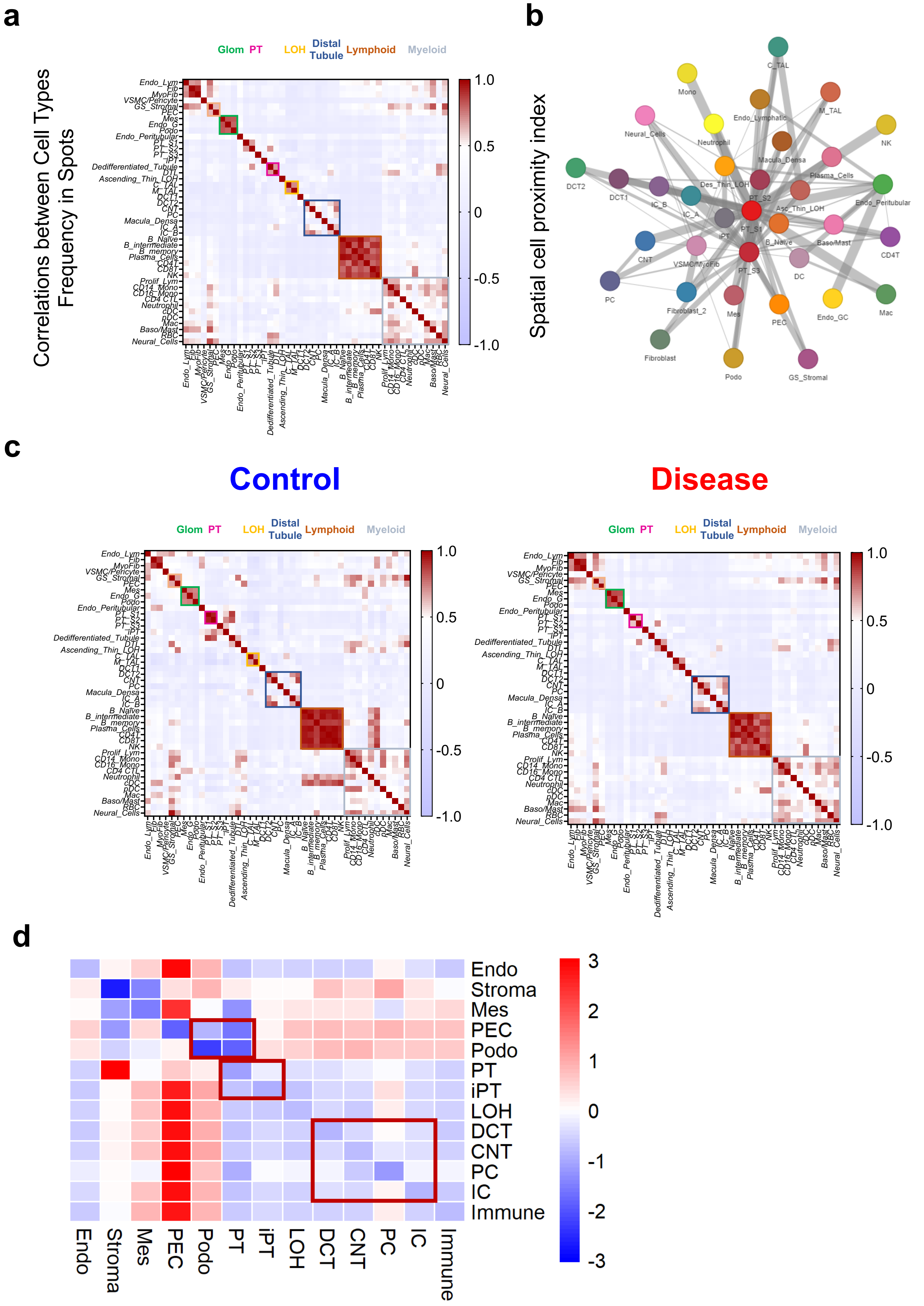


Supplementary Fig. 33

**Supplementary Fig 33. Cell type correlations and distances  of Cell Trek data.**

**(a)** Identification of neighborhoods based on spatial location of the CellTrek imputations.

**(b)** Spatial cell proximity index of CellTrek cell types.

**(c)** Neighborhoods parsed by Control and Disease sample types.

**(d)** Cell type distances between different cell types. Glomerular, proximal tubular and distal tubular neighborhoods demonstrated by red boxes.

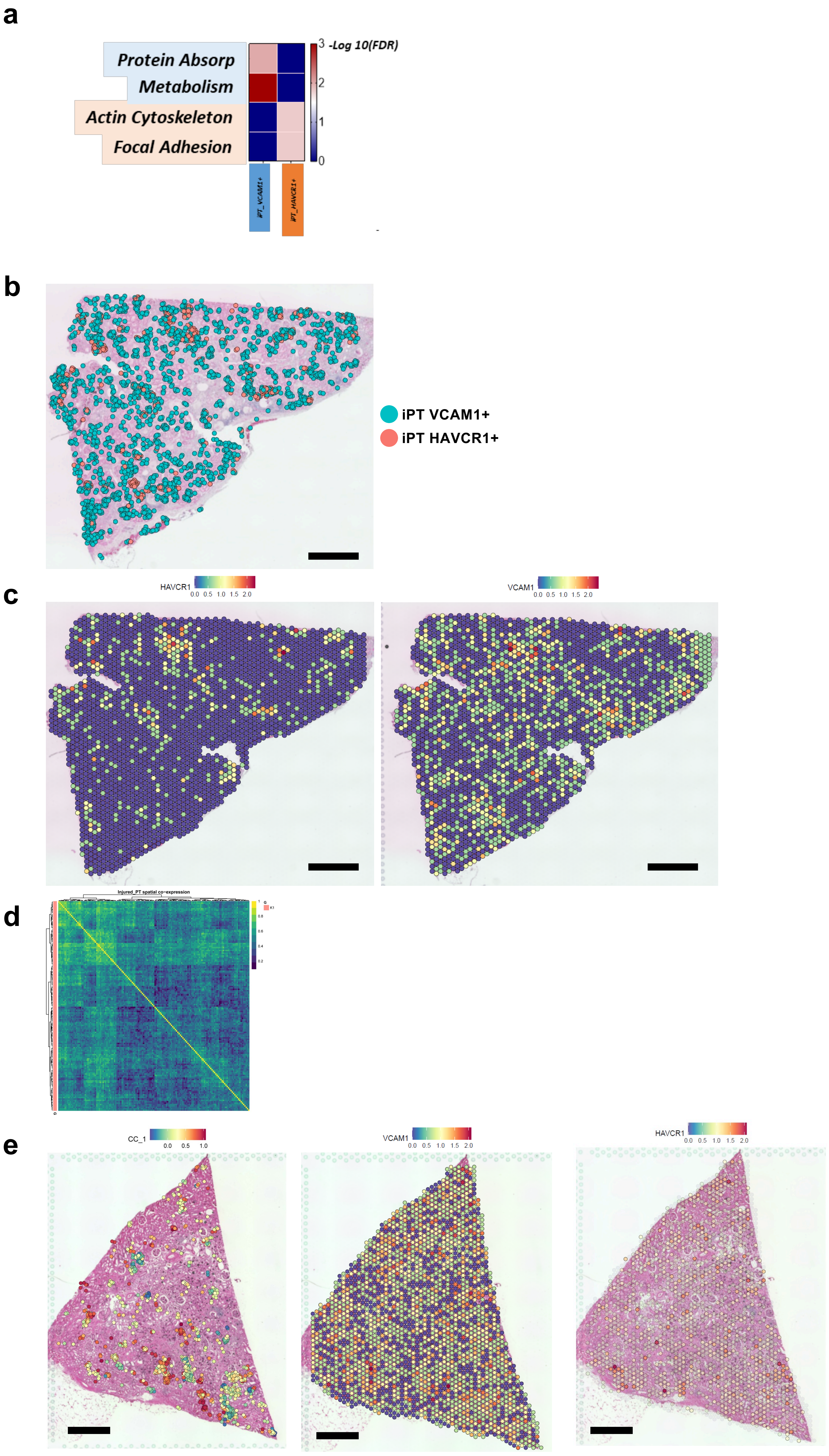


Supplementary Fig. 34

**Supplementary Fig. 34. Injured PT cells in the control and diseased human kidney samples.**

**(a)** Gene ontology enrichment for top pathways in iPT-VCAM1+ and iPT-HAVCR1+ cells.

**(b)** The spatial location of iPT_VCAM1+ (blue) and iPT_HAVCR1+ (red) cells in a diseased kidney sample based on CellTrek imputation. Scale bar is 1 mm in length.

**(c)** Expression of *HAVCR1* and *VCAM1* (color indicates level of expression). Scale bar is 1 mm in length.

**(d)** The gene co-expression network indicates one type of injured_PT in control human kidneys, upper panel shows the heatmap of co-expressed genes.

**(e)** Expression of *CC1* in imputed iPT from CellTrek, and raw *VCAM1*, and *HAVCR1* expression in a healthy human sample. Scale bar is 1 mm in length.

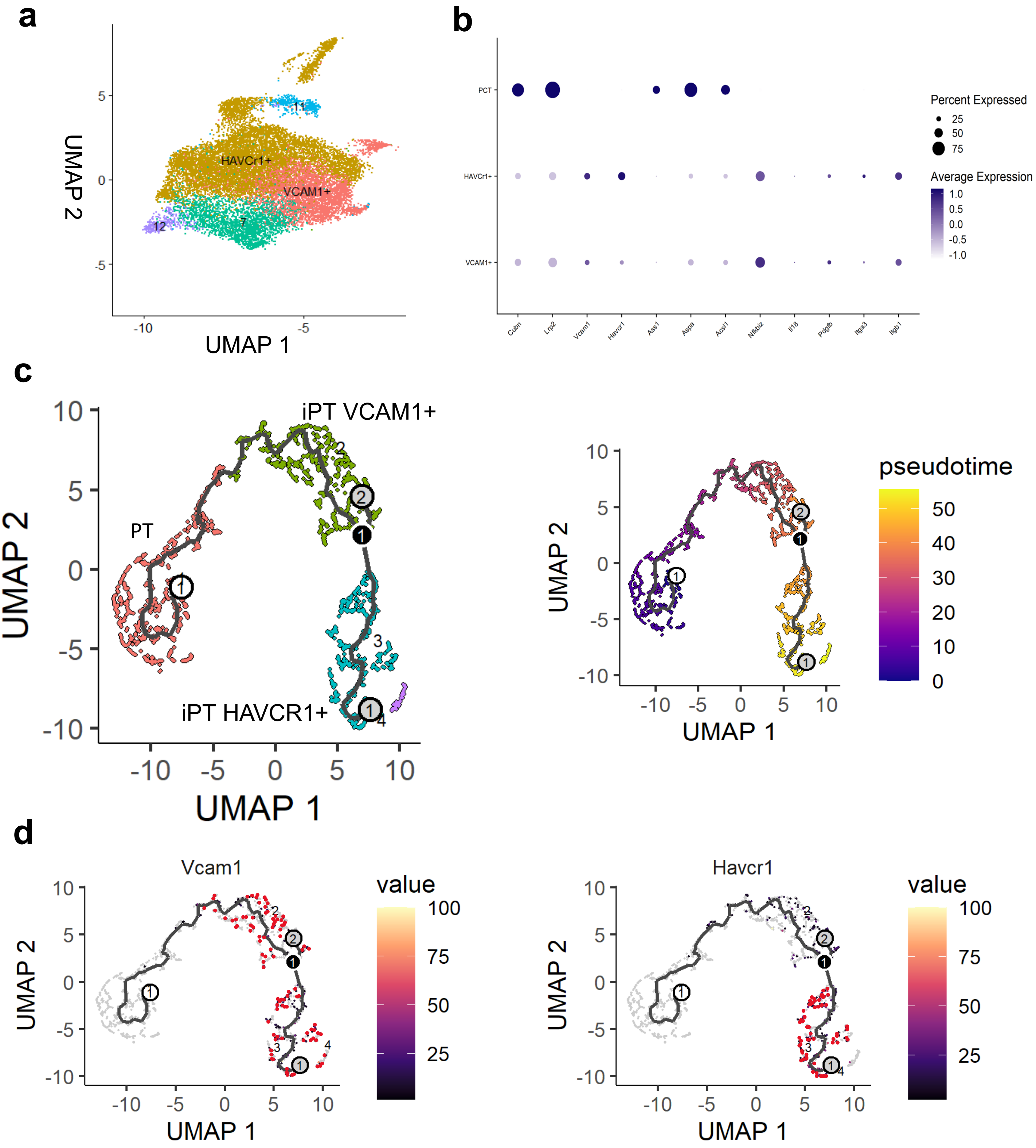


**Supplementary Fig. 35**

**Supplementary Fig. 35. Injured PT cells in mouse diabetic kidney disease samples.**

**(a)** UMAP of (n=28,579) of proximal convoluted tubule and two types of injured PT cell nuclei (left panel).

(**b**) The dot plots of marker genes used for the annotation of PCT, iPT_VCAM1+, and iPT_HAVCR1+ in the integrated dataset. The size of the dot indicates the percent positive cells, and the darkness of the color indicates average expression.

(**c**) Monocle cell trajectory of PT and iPT cells sub-clustering trajectory from PT to iPT-VCAM1+ and iPT-HAVCR1+ in mouse snRNAseq (left panel). Cells are colored by pseudotime (right panel) .

**(d)** Expression of the *Vcam1* and *Havcr1* along the trajectory.

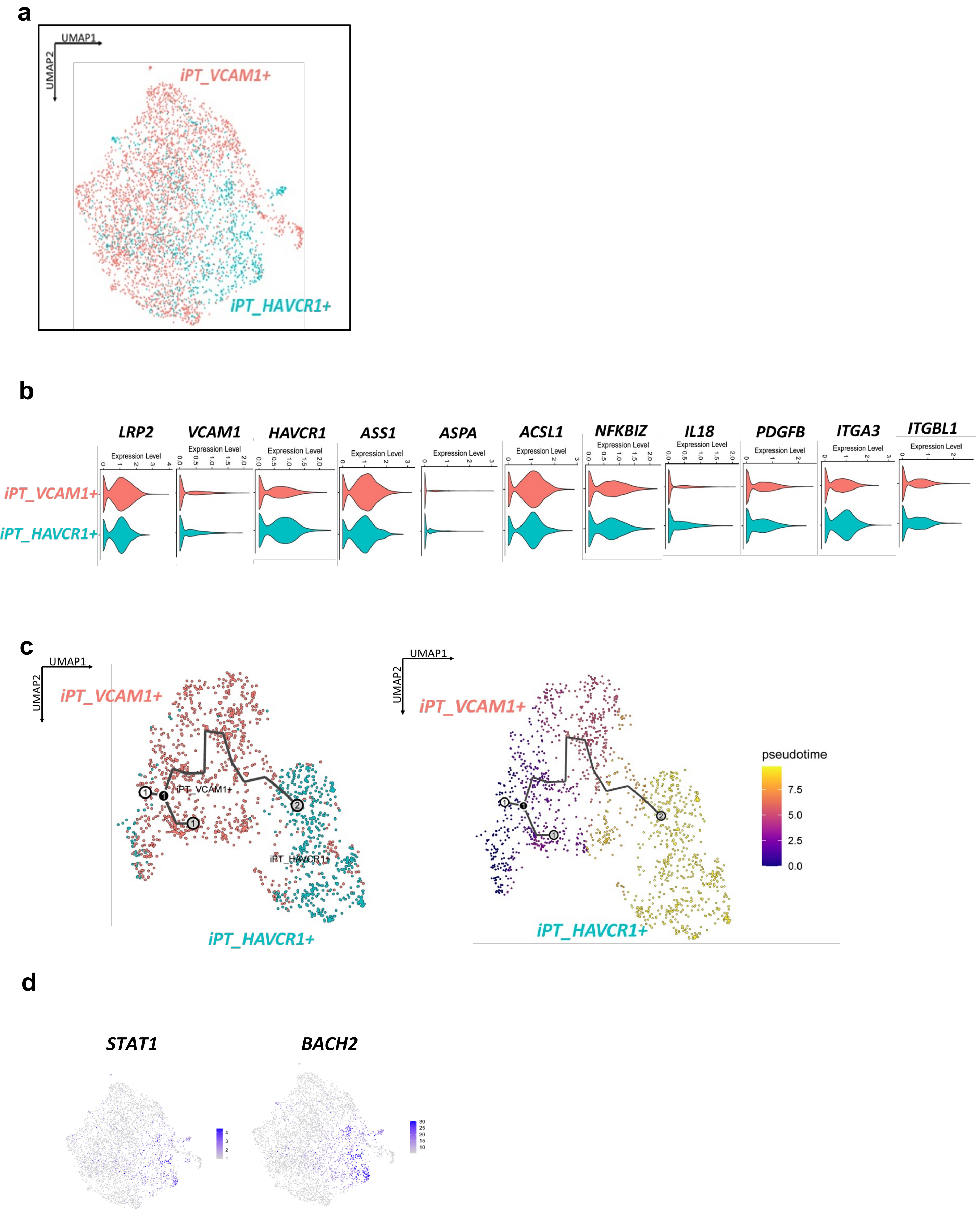


Supplementary Fig. 36

**Supplementary Fig. 36. Injured PT cells in the snATACseq data.**

(**a**) UMAP of 24,256 injured PT sub-cluster in human snATACseq dataset.

(**b**) Violin plot of expressions of VCAM1, HAVCR1 and other genes in in two types of iPTs in snATACseq dataset.

(**c**) Monocle trajectory analysis of PT and injured_PT cells in the snATACseq dataset. The heatmap shows trajectory analysis based on chromatin accessibility.

**(d)** Feature plots of transcription factor enriched motif in iPT_HAVCR1+ cell type.

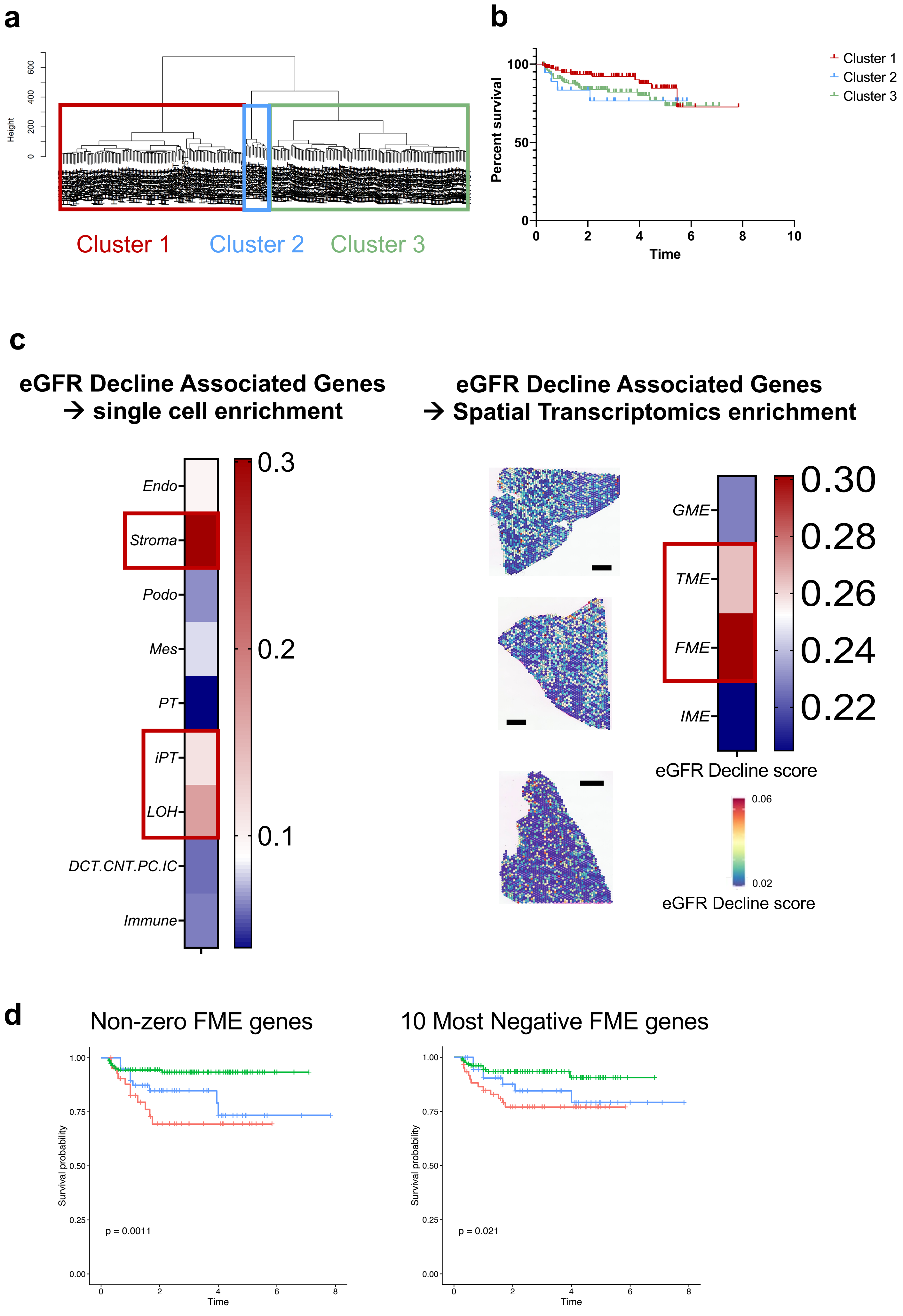


**Supplementary Fig. 37**

**Supplementary Fig. 37. FME gene expression predicts kidney outcomes**

**(a)** Hierarchical clustering of 245 human kidney tubule samples based on the expression of 1,100 randomly picked genes.

**(b)** Kaplan Meier analysis with Log-rank test was used to compare the survival of the 3 different clusters. Renal survival was defined as cases reaching end stage renal disease or greater than 40% eGFR decline.

**(c)** Single cell expression enrichment of genes associated with eGFR decline. The heatmap shows the cell type enrichment of genes associated with eGFR decline (red indicates more genes with cell type expression). Endo; endothelial cells, Stroma; stromal cells, PEC; parietal epithelial cells, Podo; podocyte, PT; proximal tubule cells, LOH; loop of Henle, DCT; distal convoluted tubule, CNT; connecting tubule, PC; principal cells of collecting duct, IC_A; Type alpha intercalated cells. Spatial expression and microenvironment enrichment of genes associated with eGFR decline. GME: glomerular, TME: tubule, FME: fibrosis, IME: immune microenvironment.

**(d)** Using the LASSO regression of all FME genes against eGFR, Kaplan Meier Analysis was re-performed using clustering of subsets of gene-- those with a Lasso coefficient that is non-zero and the genes with the most negative coefficients (Supplementary Table 13).
